# Identifying genetic variants that influence the abundance of cell states in single-cell data

**DOI:** 10.1101/2023.11.13.566919

**Authors:** Laurie Rumker, Saori Sakaue, Yakir Reshef, Joyce B. Kang, Seyhan Yazar, Jose Alquicira-Hernandez, Cristian Valencia, Kaitlyn A Lagattuta, Annelise Mah-Som, Aparna Nathan, Joseph E. Powell, Po-Ru Loh, Soumya Raychaudhuri

## Abstract

To understand genetic mechanisms driving disease, it is essential but difficult to map how risk alleles affect the composition of cells present in the body. Single-cell profiling quantifies granular information about tissues, but variant-associated cell states may reflect diverse combinations of the profiled cell features that are challenging to predefine. We introduce GeNA (Genotype-Neighborhood Associations), a statistical tool to identify cell state abundance quantitative trait loci (csaQTLs) in high-dimensional single-cell datasets. Instead of testing associations to predefined cell states, GeNA flexibly identifies the cell states whose abundance is most associated with genetic variants. In a genome-wide survey of scRNA-seq peripheral blood profiling from 969 individuals,^1^ GeNA identifies five independent loci associated with shifts in the relative abundance of immune cell states. For example, rs3003-T (p=1.96×10^-11^) associates with increased abundance of NK cells expressing TNF-α response programs. This csaQTL colocalizes with increased risk for psoriasis, an autoimmune disease that responds to anti-TNF treatments. Flexibly characterizing csaQTLs for granular cell states may help illuminate how genetic background alters cellular composition to confer disease risk.

## Main Text

### Background

Genome-wide association studies (GWASs) have identified thousands of disease-associated loci.^2,3^ Examination of biologic samples from human population cohorts can reveal associations of these same loci to molecular, cellular and tissue traits, offering insight into disease development processes to inform novel treatment strategies.^4^ Many previous studies have linked variants to molecular phenotypes by identifying quantitative trait loci (QTLs) that change the expression^5,1,6,7^ (eQTL) or splicing^8–10^ (sQTL) of a single gene, the abundance of a protein in serum^11,12^ or cells^13–15^ (pQTL), or the epigenetic state^16–20^ (caQTL) of a single genomic site. However, molecular QTLs alone do not fully explain the essential alterations in tissue function that ultimately lead to disease.^21–23^ We must, therefore, also identify how disease-associated variants change tissue function, as reflected by altered abundances of functional cell states.

Previous investigators have explored genetic links to the abundance of predefined blood cell types using flow cytometry, yielding associations that often colocalize with disease risk.^15,24–31^ High-dimensional single-cell assays present a new opportunity to expand upon flow cytometry-based approaches. Through unbiased profiling of thousands of features per cell, these datasets reveal a more granular landscape of tissue composition within which we might detect a greater variety of genetically associated changes. Furthermore, high-dimensional profiling technologies can readily be applied to tissues that are difficult to disaggregate or lack a broad repertoire of markers for flow-sorting, as illustrated by single-nuclear profiling of brain and muscle samples and of frozen biobanked tissue.^32–34^ However, these high-dimensional single-cell datasets present new statistical challenges. Instead of modeling genetic associations to a one-dimensional dependent variable (e.g., expression of one gene or abundance of one predefined cell type), analysis of these datasets requires a flexible approach capable of detecting genetic associations to many cell states, each defined by different combinations of the profiled cell features.

We introduce GeNA (Genotype-Neighborhood Associations), a tool to conduct genome-wide surveys for cell state abundance quantitative trait loci (csaQTLs) in high-dimensional single-cell data. We use “cell state” broadly to connote any group of cells with shared features. For example, variants could alter the abundance of a canonical discrete cell type like naïve B cells or cells with a shared active gene expression program like IL-2-responding B cells. Genetic associations to the abundance of cytokines^11,35^ also suggest that a single genetic variant may associate with the abundance of cell states within multiple cell types that respond to a shared signaling molecule. In simulations, GeNA has strong statistical power to identify associations to many cell states while demonstrating well-controlled type I error. In a genome-wide csaQTL survey within a real single-cell dataset, GeNA reveals novel csaQTLs that replicate in independent datasets and colocalize with immune-mediated disease risk loci. GeNA provides insight into tissue compositional changes at single-cell resolution that may contribute to disease development.

### Overview of GeNA

GeNA takes as input a cohort of genotyped individuals with single-cell profiling of one sample per individual (**Methods;** **Fig. 1**). GeNA uses a framework from our previous work^36^ to take a nearest neighbor graph representation of the single cells and quantify cellular fractional abundance per individual across many small regions of the cell state space, termed neighborhoods. The cell abundance distributions per individual are represented in a samples-by-neighborhoods matrix (Neighborhood Abundance Matrix, NAM). PCA is used to define the top principal components of the NAM (NAM-PCs). NAM-PCs reflect the primary axes of cell state abundance variation across individuals and aggregate information across neighborhoods. Previously, we employed NAM-PCs to detect cell state associations to a single clinical attribute, such as treatment response, with the tool Covarying Neighborhood Analysis (CNA).^36^ As in CNA, rather than test abundance associations for individual neighborhoods, which would impose a high multiple-testing burden, GeNA performs a single association test per variant to the top NAM-PCs, with control for sample-level confounders like demographic variables and technical effects. In contrast to CNA, in GeNA we employ a new statistical model that now enables detection of genotype associations to NAM-PCs at genome-wide scale.

**Figure 1:**
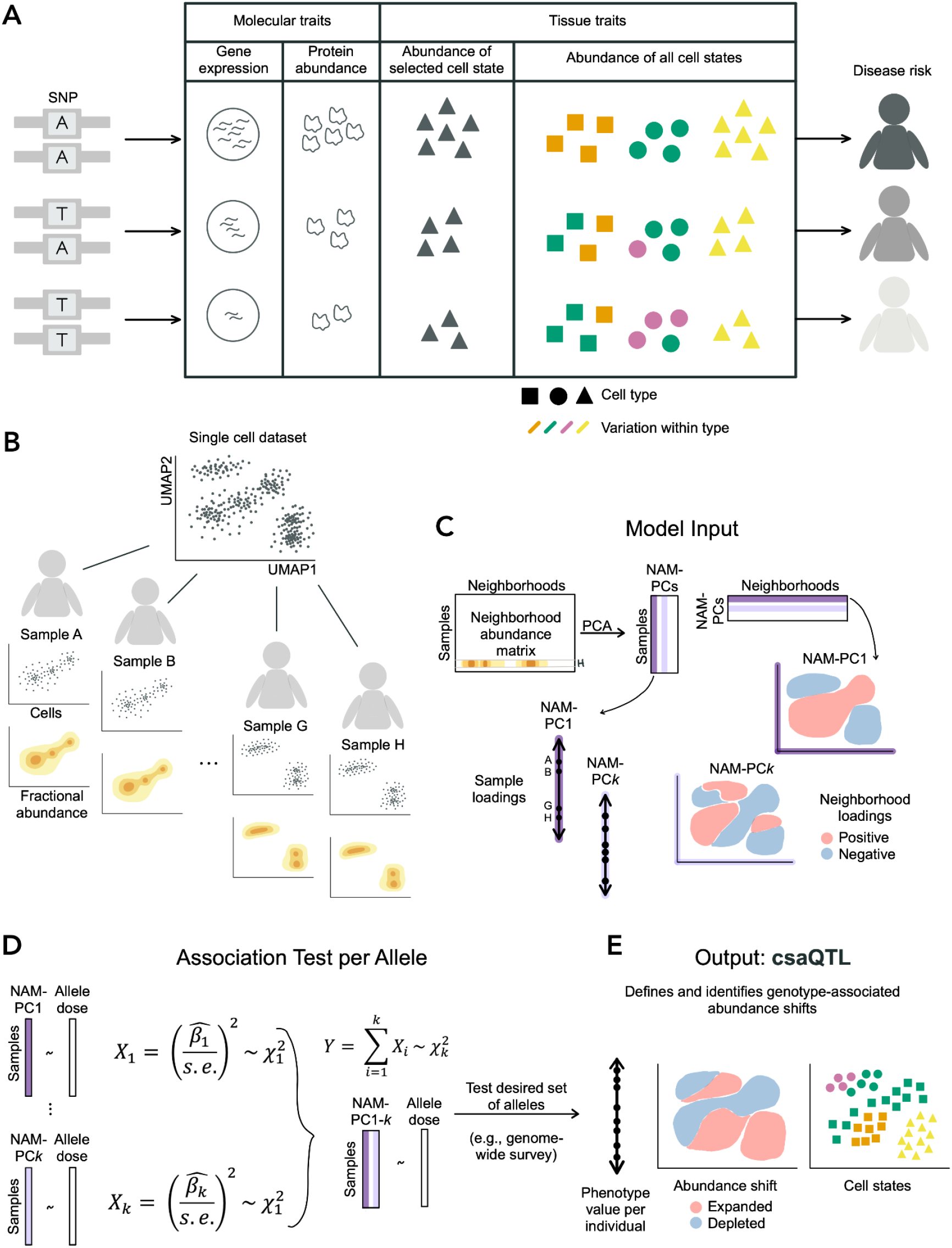
Method schematic. (**A**) Consider a variant associated with disease risk. Intermediate traits that may also associate with the variant and even mediate the genetic risk include well-studied molecular traits (e.g., transcript or protein abundance) as well as changes in the abundance of cells with varying character and function, illustrated here by variation in shape and color. Previous genetic studies of cell state abundance traits have quantified target cell states (e.g., triangle cell type) using flow cytometry. High-dimensional profiling may reveal genetically associated variation in the abundance of cell states researchers may not anticipate or cannot flow-sort (e.g., green character within the circle cell type). Detection of such associations requires granular information about cell variation and a flexible method for detecting variant-associated cell states. (**B**) For a given single-cell dataset, we use the landscape of cells observed for each sample to compute a granular distribution of fractional cellular abundance across the total cell state space, conceptually illustrated here in two dimensions. We illustrate a single axis of compositional variation across the four samples shown. (**C**) The Neighborhood Abundance Matrix stores the fractional abundance of cells from each sample in each neighborhood. Sample H is highlighted as an example. Principal components analysis of this object yields sample and neighborhood loading information on NAM-PCs. We illustrate with NAM-PC1 how these loadings would reflect the compositional axis illustrated in (B). We show another possible component (NAM-PC*k*) to illustrate that NAM-PCs can capture co-variation across transcriptionally distant regions. (**D**) GeNA uses a test statistic Y, which follows a chi-squared distribution with *k* degrees of freedom, to detect an association between allele dose for a given SNP and any systematic change in tissue cellular composition captured by NAM-PC1-*k* (**Methods**). (**E**) We illustrate how the cell state abundance shift shown in (A) might be revealed using GeNA.

To discover csaQTLs, GeNA tests whether the alternative allele dose for each single nucleotide polymorphism (SNP) is associated with any linear combination of sample loadings on the top *k* NAM-PCs. For a given SNP, for each NAM-PC GeNA computes a chi-squared statistic with one degree of freedom to reflect the relationship between genotype and that NAM-PC. The sum of these statistics across NAM-PC1-*k* follows a chi-squared distribution with *k* degrees of freedom, from which we obtain a p-value per SNP (**Methods**). Each SNP is tested only once, so we consider loci with p<5×10^-8^ associations genome-wide significant.

For each detected csaQTL, we define the lead SNP as the SNP with the strongest association to any cell state abundance shift. To identify the cell states impacted by the lead SNP, we compute the abundance correlation per neighborhood to allele dose (“neighborhood-level phenotype”). We define the sample-level phenotype value per individual as the linear combination of sample loadings on NAM-PC1-*k* from the fitted model for the lead SNP (**Methods**). This per-sample value reflects the degree to which the SNP-associated pattern of cell state abundance characterizes that sample’s total cell abundance across all states.

Replication in independent datasets is crucial for any genotype-trait association. We devised an approach to project a neighborhood-based cell state abundance phenotype into an independent dataset using reference mapping (**Methods**). This approach supports replication testing for any phenotype found using NAM-PCs, including GeNA csaQTLs and CNA case-control analyses (**Supp. Note**).

GeNA requires <25 minutes and <17 GB for a genome-wide survey (6.4M SNPs) in a dataset of >800,000 cells from >950 individuals. We have released open-source scripts implementing GeNA (**URLs**).

### Performance assessment with simulations

We used a real single-cell dataset and simulated genotype values to assess GeNA’s calibration and statistical power by estimating type I and type II error, respectively. This published dataset contains 822,552 scRNA-seq profiles of peripheral blood mononuclear cells (PBMCs) from 969 individuals (“OneK1K cohort”),^1^ approximately half of whom have documented clinical metadata. To assess type I error, we simulated random genotypes in Hardy-Weinberg equilibrium without true associations to the single-cell data (**Methods**). We tested for csaQTL effects within five major cell type groups (NK, T, B, myeloid, and all cells). Among other quality control measures (**Methods**), we excluded samples with <25 cells, leaving N_T_=968, N_B_=910, N_NK_=935, N_myeloid_=523 and N_all cells_=969 donors. We observed well-calibrated p-values, with p<0.05 in 18,465/400,000 trials (type I error rate at α=0.05 of 0.046±0.0063). Type I error was consistent across cell types, *k* values, and minor allele frequencies (MAFs; **Supp.** Fig. 1-4). To estimate statistical power, we defined real cell-state abundance traits within each major cell type and simulated many genotypes associated with each trait reflecting increasing amounts of noise (**Methods**). In flow cytometry studies, real SNPs have explained 0.8-13.8% (5th-95th percentile)^15^ of variance in the abundance of tested cell states. GeNA demonstrates 37% and 86% power at p<5×10^-8^ to detect associations for SNPs that explain 6% and 12% of phenotypic variance, respectively (**Supp.** Fig. 5-6). Notably, within the simulated T cell GWAS GeNA detected associations to many distinct simulated traits, including 14 traits with pairwise sample-level phenotype r^2^<0.2 (**Supp.** Figs. 7-8). Therefore, these simulations demonstrate that in a single GWAS GeNA can detect multiple csaQTLs that each associate with a distinct trait, without requiring parameter tuning.

### csaQTLs detected in the OneK1K dataset

We performed five total csaQTL GWASs in the OneK1K dataset: one to detect genetic associations to cell state shifts across all PBMCs, and one each within T, B, NK and myeloid cells. As in our simulations, without parameter tuning GeNA identified multiple real csaQTLs associated with uncorrelated traits (r^2^<0.2 between sample-level trait values) in one single-cell dataset (**Supp.** Figs. 9-10). We detected five csaQTLs passing genome-wide significance (p<5×10^-8^; **Fig. 2A; Supp. Table 1; Supp.** Fig. 11-13): four associated with shifts in NK cell states (**Fig. 2B-D,F**) and one associated with shifts in myeloid cell states (**Fig. 2E**). We permuted genotype values across samples 10^6^ times per csaQTL lead SNP; no permuted genotype resulted in a GeNA p<5×10^-8^. While two csaQTLs were associated with shifts in the relative abundance of cell states that correspond to the published clusters in this dataset (2q13, 15q25.1; **Fig. 2B, 2E**), three csaQTLs affect phenotypes not captured by the published clusters (**Fig. 2C-D, 2F**). Consistent with this observation, when we conducted GWASs of the published clusters’ abundances, we identified the 2q13 and 15q25.1 csaQTLs but not the other three (**Methods; Supp. Table 2, Supp.** Fig. 14-15). Also correspondingly, the 2q13 and 15q25.1 csaQTLs found by GeNA directionally replicate genotype-phenotype associations previously identified using flow cytometry,^15,37^ while the remaining three GeNA csaQTLs represent novel associations (**Supp. Table 3**).

**Figure 2:**
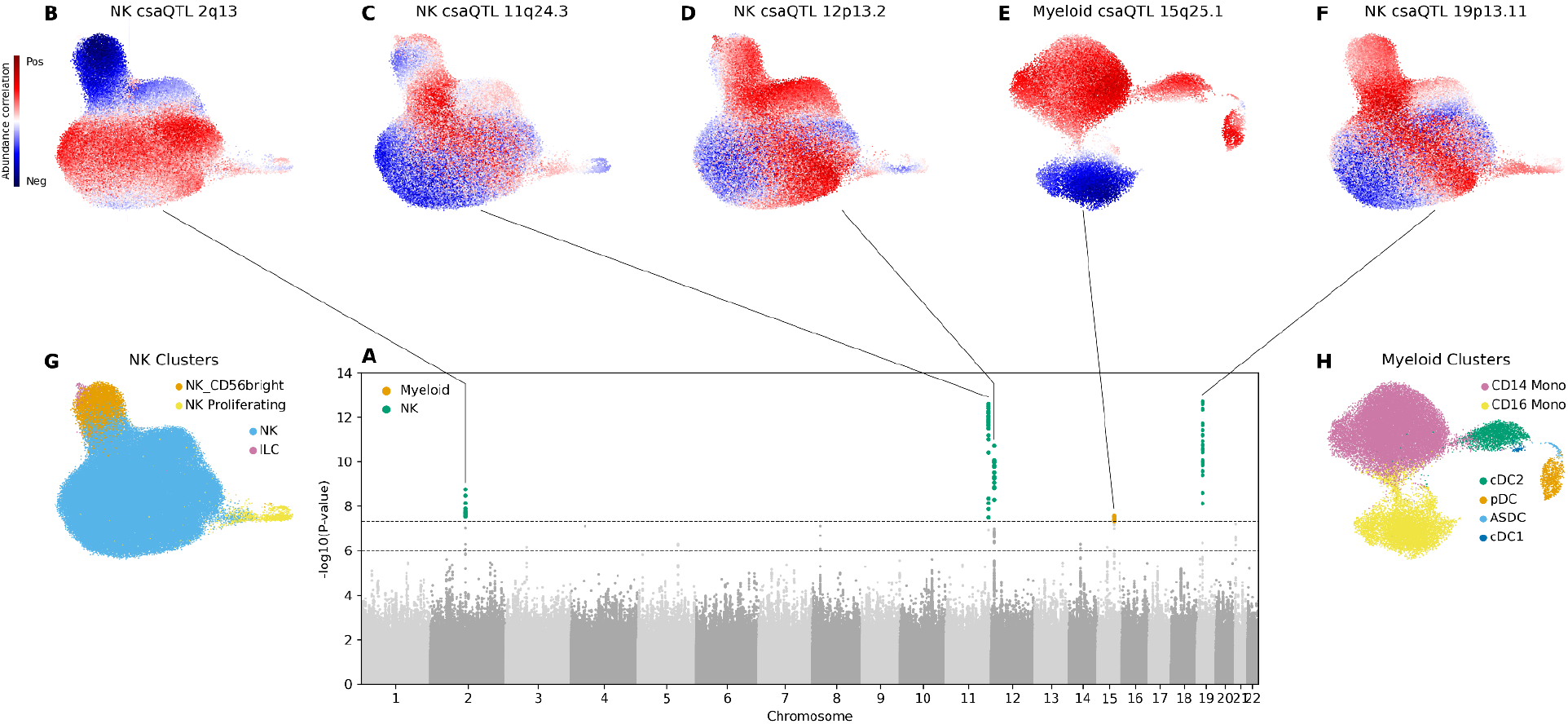
csaQTLs detected in the OneK1K dataset. (**A**) Superimposed Manhattan plots for csaQTL GWASs among NK cell states and among myeloid cell states. A genome-wide significance threshold of p<5×10^-8^ is indicated by a dashed line. SNPs with p-values less than this threshold are colored according to their source GWAS: NK cells (green) or myeloid (orange). Another dashed line indicates a p<1×10^-6^ threshold for suggestive associations. (**B-F**) Cell abundance correlation per neighborhood to dose of alternative allele is shown in UMAPs for each of the five genome-wide significant loci: (**B**) NK csaQTL 2q13, (**C**) NK csaQTL 11q24.3, (**D**) NK csaQTL 12p13.2, (**E**) myeloid csaQTL 15q25.1, and (**F**) NK csaQTL 19p13.11. (**G-H**) Cell type cluster labels from Yazar*, Alquicira-Hernandez*, Wing*, *et al.*^1^ for (**G**) NK and (**H**) myeloid cells are shown for reference.

To confirm the novel associations, we examined five replication cohorts with PBMC scRNA-sequencing representing 428 total individuals (**Supp. Table 4**).^38–40^ The novel csaQTLs replicate (meta-analysis p<0.05) with directional concordance (**Methods; Supp.** Fig. 16).

To characterize the cellular composition changes associated with each locus, the anchor cell for each neighborhood can serve as an estimate for the transcriptional state represented by the neighborhood. We quantified the correlation of the neighborhood-level phenotype to expression for individual genes, then used those correlation values per gene as input to gene set enrichment analysis (**Methods; Supp. Tables 5-6**). We then identified molecular and clinical traits that colocalize^41^ with the csaQTLs (**Methods; Supp. Tables 7-8**).

For the csaQTL at 12p13.2, for example, lead SNP rs3003-T associates with expansion of activated NK cell states expressing TNF-α, IFN-γ, IL-2 and IL-6 response genes (**Supp. Tables 5-6**; **Fig. 3A-E**), suggesting increased sensitivity to or abundance of these cytokines. This csaQTL colocalizes with an eQTL for *KLRC1* (eQTL p=1×10^-41^ in OneK1K NK cells, Pr_coloc_=96% probability of a shared causal variant; **Methods**) and colocalizes with risk for psoriasis^42,43^ (GWAS p=4.17×10^-9^, Pr_coloc_=91%), where rs3003-T associates with increased psoriasis risk (**Fig. 3F**). Notably, elevation of TNF-α and IFN-γ is well-documented in psoriasis.^44^ The combination of clinical metadata, genotyping and immune profiling available for the OneK1K cohort enables us to evaluate whether the csaQTL effect is fully explained by the presence of psoriasis disease itself. However, in a OneK1K subcohort with documented absence of autoimmune disease, this csaQTL is still evident (N=454, p=3×10^-4^; **Methods**; **Supp. Table 9**). Importantly, anti-TNF medications have established efficacy in psoriasis treatment^44^ and JAK inhibitors that blunt IFN-γ response have shown promise in clinical trials, suggesting that TNF-α and IFN-γ contribute to psoriasis pathology.^45^ This example locus illustrates that csaQTLs may help connect relevant biologic processes to disease risk loci.

**Figure 3:**
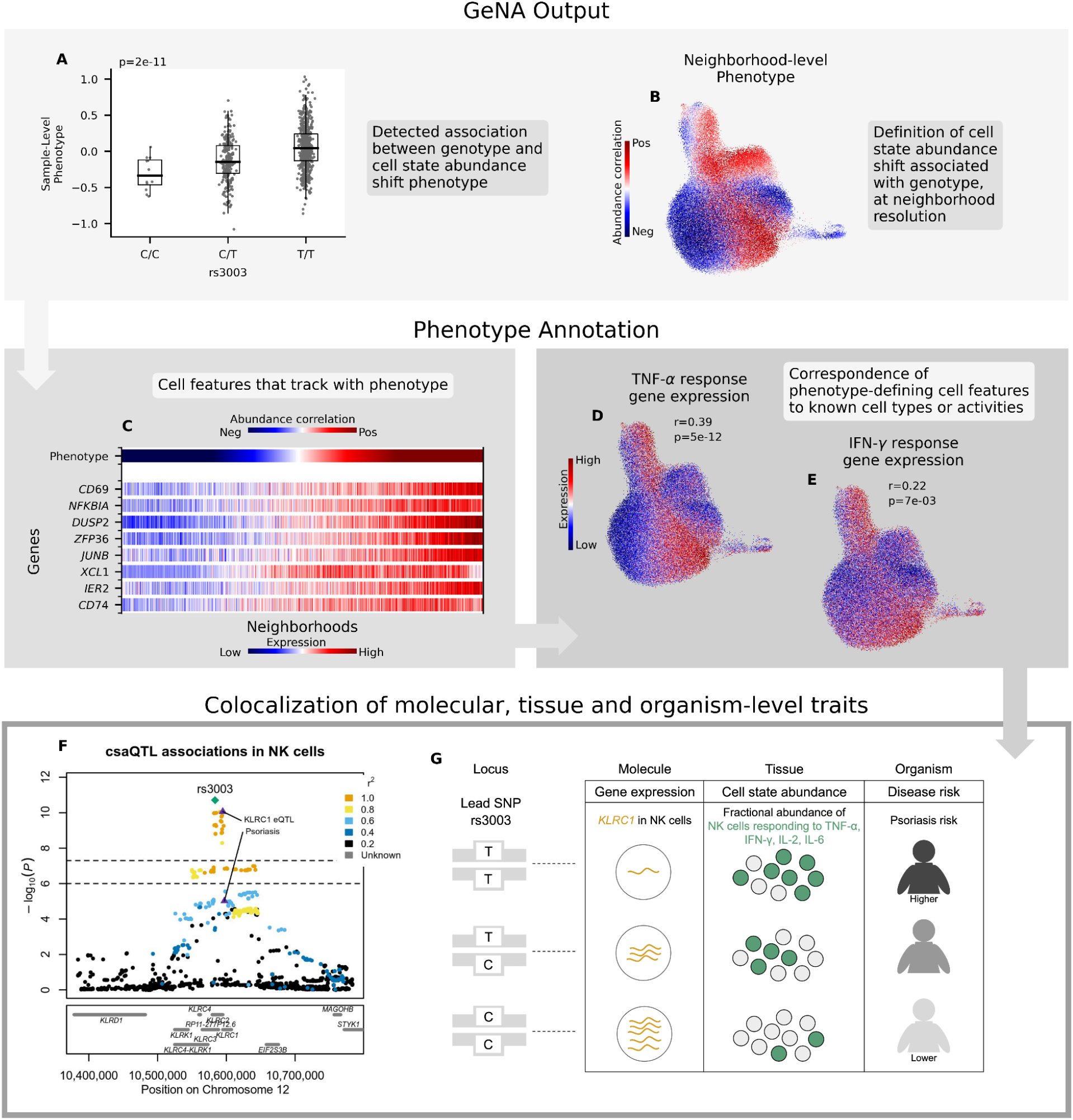
Characterization of the csaQTL at 12p13.2. (**A-B**) GeNA output: (A) Boxplot of sample-level phenotype values for each individual organized by genotype at the lead SNP. We also show the GeNA p-value. (B) UMAP of NK cells colored by neighborhood-level phenotype value (i.e., correlation between cell abundance and dose of alternative allele per neighborhood). (**C**) Heatmap of expression across neighborhoods for genes with strong correlations in expression to csaQTL neighborhood-level phenotype. Neighborhoods are arrayed along the x-axis by phenotype value. The phenotype-correlated genes include general markers of NK activation (*CD69*, *NFKBIA)* as well as TNF-α (*DUSP2, ZFP36, JUNB, IER2*) and IFN-γ (*CD74*, *XCL1*) response genes. (**D-E**) Gene set enrichment analysis identified significant activation of TNF-α and IFN-γ response pathways in association with the csaQTL phenotype. We show UMAPs of NK cells colored by summed expression of (D) TNF-α response genes and (E) IFN-γ response genes. We report the Pearson’s r across neighborhoods between phenotype values and summed expression within the gene set. We also show the FDR-adjusted gene set enrichment p-value. (**F**) Locus zoom plot with one marker per tested SNP, genomic position along the x-axis, and GeNA p-value on the y-axis. Each SNP marker is colored by linkage disequilibrium (LD) value relative to the lead SNP. The csaQTL lead SNP is labeled with a green diamond. The psoriasis risk and *KLRC1* eQTL lead SNPs are labeled with purple triangles. (**G**) Diagram of genotypes for the csaQTL lead SNP and colocalizing associations to molecular, tissue and organism-level traits at this locus.

As another example, we highlight the csaQTL at 15q25.1, where the GeNA lead SNP rs3826007-T was previously found by several studies to be associated with decreased count of flow-sorted monocytes (all GWAS p<5×10^-83^, lead SNPs LD≥0.95 to rs3826007-T).^24,25,46^ Subsequently, another study flow-sorted CD14+ and CD16+ monocytes separately, revealing an underlying association of this csaQTL to selective depletion of CD16+ monocytes^15^ (GWAS p=5×10^-9^, Pr_coloc_=99% with GeNA csaQTL). Similarly, single-cell profiling could uncover differential genetic impacts on granular cell states that may underlie previously detected associations. However, GeNA reveals that rs3826007-T associates with similar abundance changes across all CD16+ monocyte neighborhoods (**Supp.** Fig. 17).

This example locus also illustrates that csaQTLs may help generate biological hypotheses that connect to existing molecular trait associations. The rs3826007-T allele encodes a glycine to aspartic acid missense change in *BCL2A1* that is likely to be deleterious (SIFT^47^ score 0.036, PolyPhen2^48^ scores HDVIV 1 and HVAR 1) and predicted to produce a dysfunctional form of the BCL2A1 protein. BCL2A1 is an essential pro-survival factor^49^ and *BCL2A1* is preferentially expressed in CD16+ monocytes among PBMCs (differential expression p<1×10^-10^; **Methods; Supp.** Fig. 18). Interestingly, rs3826007-T was also previously found to be associated with decreased *BCL2A1* expression in whole blood^50^ (eQTL p=9.3×10^-52^, Pr_coloc_=98%; **Supp. Table 8**). Therefore, this csaQTL may yield a dysfunctional BCL2A1 protein, upon which CD16+ monocytes in particular are dependent for survival, leading to decreased CD16+ monocyte fractional abundance and thereby decreased *BCL2A1* expression in whole blood. This csaQTL also colocalizes with risk for primary sclerosing cholangitis (PSC)^51^ (GWAS p=1×10^-6^; Pr_coloc_=98%), a disease in which monocytes have been proposed to play important roles,^52^ and the csaQTL is evident in a subcohort of OneK1K individuals with documented absence of autoimmune disease (N=247, p=7×10^-5^; **Methods; Supp. Table 9**).

We applied a similar approach to characterize the other csaQTLs detected in the OneK1K dataset (**Supp.** Fig. 19-21), which revealed additional colocalizing associations to molecular traits and disease risk. The csaQTL at 11q24.3 colocalizes with a suggestive eQTL for the *ETS1* transcription factor^53^ (**Supp.** Fig. 19) and the csaQTL at 2q13 colocalizes with a pQTL for the abundance of chemokine CXCL16 in serum^12^ (**Supp.** Fig. 21). The csaQTL at 19p13.11 colocalizes with risk for asthma^54^ (GWAS p=1×10^-8^; Pr_coloc_=98.5%) and type 1 diabetes^55^ (GWAS p=7×10^-7^; Pr_coloc_=98.8%), and the csaQTL at 2q13 colocalizes with risk for epithelial ovarian cancer^56^ (GWAS p=2×10^-8^; Pr_coloc_=80%). For each csaQTL and corresponding disease with a colocalizing association, the csaQTL effect is evident within a OneK1K subcohort with documented absence of the corresponding disease (**Supp. Table 9**).

Because we observed colocalizing cis-eQTLs for some csaQTLs, we considered whether the csaQTLs might be driven primarily by cis-eQTL effects (i.e., the csaQTL expands or depletes cell states characterized by genes close to the csaQTL). To evaluate this possibility, for each csaQTL we created a custom version of the OneK1K dataset in which we removed expression information for all genes within a two-megabase window centered on the lead SNP then applied dataset quality control and graph construction and ran GeNA (Methods). We observed that each csaQTL was strongly sustained (p<3×10^-8^), with fidelity between the discovery and masked datasets in neighborhood-level (r^2^≥0.95) and sample-level (r^2^≥0.91) phenotypes (Supp. Table 10). Therefore, the csaQTLs reveal cell state abundance shifts that are not explained solely by cis-eQTLs. GeNA csaQTLs hold promise for implicating not just individual molecules but broad cellular functions in genetic disease risk.

### Shared effects of disease risk loci highlight disease-relevant cell states

Individual genetic variants associated with a given disease may exert their effects through shared biologic pathways. We sought to test whether SNPs associated with a single polygenic disease might alter similar cell states and to directly compare genetically associated states with peripheral blood changes observed during the disease itself. Perez *et al*. published the first large case-control study for an autoimmune disease—systemic lupus erythematosus (SLE)— with single-cell profiling of peripheral blood (1.2M PBMCs, 162 cases, 99 controls).^38^ Genetic background explains 43-66%^57^ of risk for SLE, a chronic illness with limited treatment options in which >50% of diagnosed individuals develop organ damage within 10 years.^58^ Chen *et al*.^59^ published a polygenic risk score (PRS) to estimate any individual’s total lupus genetic risk^60^ based on their genotypes at 95 genome-wide significant loci. Using this PRS, we estimated lupus genetic risk per OneK1K cohort individual and tested for PRS-associated cell states across all cells and within four major cell types using CNA^36^ (**Methods; Supp. Table 11**). To avoid PRS values acting as a proxy for clinically documented disease status, we included only OneK1K individuals with a documented absence of SLE. We further tested whether found associations persisted among OneK1K individuals with a documented absence of any autoimmune disease.

While no individual SLE risk SNP was identified as a significant csaQTL by GeNA, several SNPs in the SLE PRS are strongly correlated with the abundance of IFN-α-responding myeloid cell states, including SNPs near interferon-related genes^61,62^ (**Fig. 4A-B**). When the effects of individual SNPs were aggregated in the PRS, individuals with higher SLE genetic risk^59^ (but without disease) were found to share significant expansion of myeloid cell states responding to IFN-α (CNA Global p_FDR-adj._=0.04; GSEA p-value=1.5×10^-9^; **Fig. 4C-D**). Substantial existing evidence supports the importance of interferon in SLE,^63^ including the proximity of many SLE GWAS loci to interferon-related genes^64^ and the recent approval of interferon-targeting SLE treatments.^65^ To our knowledge, however, we offer the first direct evidence that genetic risk for SLE increases IFN-α signaling among individuals without a SLE diagnosis. Expansion of IFN-α-responding cell states also differentiates the peripheral blood of SLE patients from controls (**Fig. 4E-F**; **Supp.** Fig. 22). Our analyses contribute complementary evidence to support a causal role for IFN-α in lupus development. csaQTL analyses may help illuminate the convergence of effects from distinct disease risk loci on shared functional pathways.

**Figure 4:**
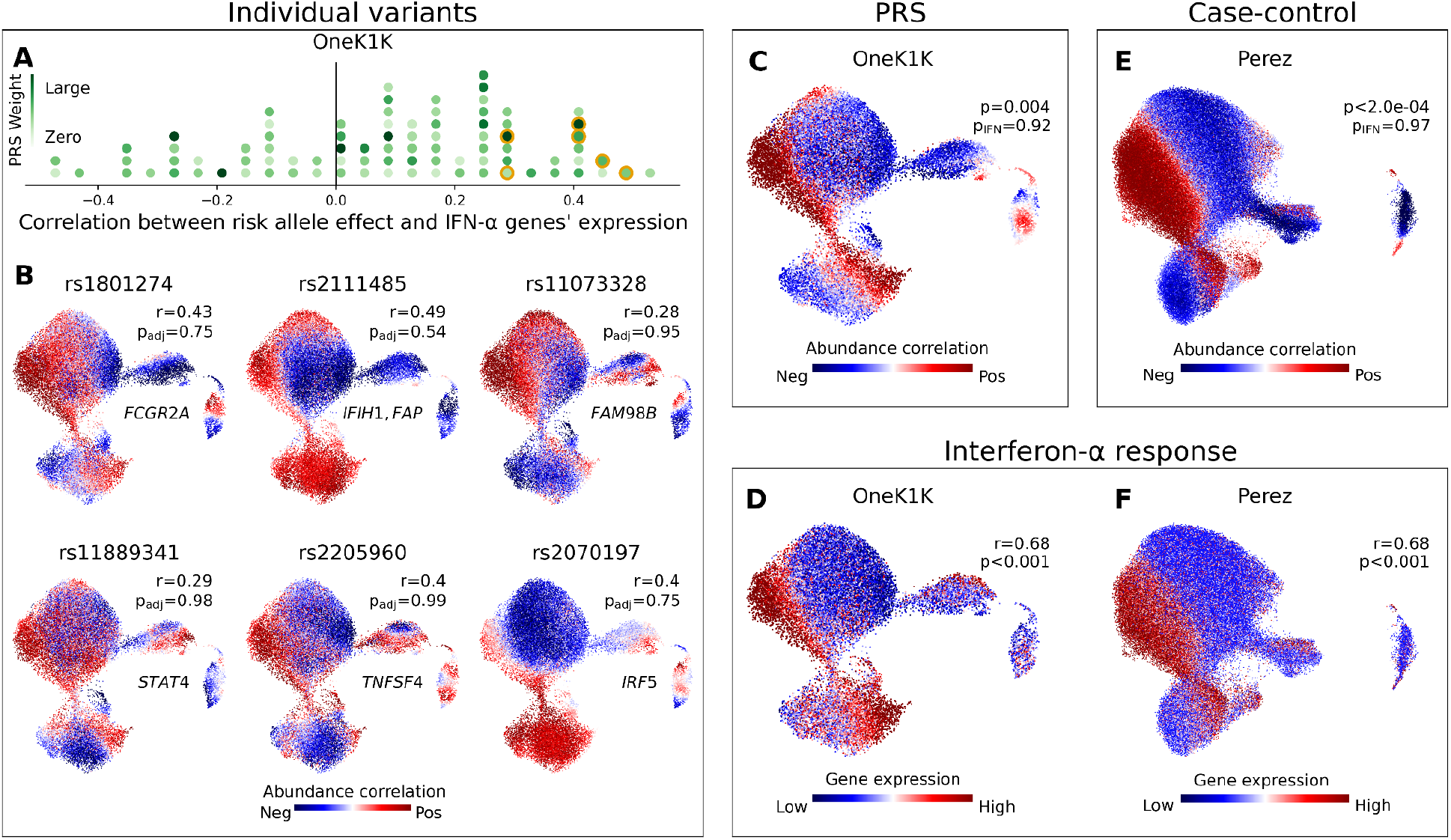
Polygenic risk scores aggregate the effects of individual loci to highlight disease-relevant cell states, a valuable point of comparison to single-cell case-control analyses. (**A**) Histogram of SNPs in the SLE PRS. For each SNP, we plot the Pearson’s r correlation across OneK1K myeloid neighborhoods between the SLE risk-associated phenotype and an IFN-α response gene signature. The marker for each risk allele is colored according to its effect weight in the PRS. Six SNPs plotted in (B) are highlighted in orange. (**B**) We show six SNPs in the SLE PRS for which the myeloid cell state abundance correlations to the SLE risk allele correspond closely to an IFN-α response signature. For each selected SNP, we plot a UMAP of OneK1K myeloid cells colored by the abundance correlation per neighborhood to dose of the risk allele. We also report the gene(s) to which the SNP has been mapped, the Pearson’s r correlation between the neighborhood-level phenotype and IFN-α response signature, and the FDR-adjusted CNA global p-value. (**C**) Myeloid cell state abundance shift associated with increasing SLE PRS value in the OneK1K cohort. CNA global p values are shown with (p_IFN_) and without (p) controlling for mean IFN-α response gene expression per individual. (**D**) IFN-α response gene expression per neighborhood among myeloid cells in the OneK1K cohort. Pearson’s r between IFN-α response per neighborhood and the PRS phenotype from (C) is shown, with associated bootstrapped p-value for r>0. (**E**) Myeloid cell state abundance shift associated with SLE disease status in the Perez *et al*. European cohort. CNA global p-values are shown with (p_IFN_) and without (p) controlling for mean IFN response gene expression per individual. (**F**) IFN response gene expression per neighborhood among myeloid cells in the Perez *et al*. European cohort. Pearson’s r between IFN response per neighborhood and SLE phenotype from (E) is shown, with associated bootstrapped p-value for r>0.

Not all tissue composition changes associated with genetic risk for a disease may mirror the disease state itself. The quantity of patients with rheumatoid arthritis (RA) in the OneK1K dataset enabled us to pursue the same analysis with a second autoimmune disease.^66^ We detected a significant case-control difference (all-cells CNA Global p<0.0001) and an association to RA PRS (all-cells CNA Global p_unadj._=0.027) (**Supp. Tables 11-12**). Cell abundance correlations per neighborhood from both tests suggested specific shifts in T cell subtypes, which we then tested directly using the published cell type clusters (**Supp.** Fig. 23). Among CD4+ T cells the naïve-to-effector ratio (T_naïve_/[T_EM_+T_CTL_]) increased with higher RA genetic risk (permutation p<0.05) but was decreased in RA disease relative to controls (t-test p<0.05). When disease-associated and PRS-associated shifts differ, detecting cell states altered by genetic background may help disentangle disease-driving processes from disease sequelae.

### Future applications

GeNA identifies five csaQTLs in the OneK1K cohort, a comparable count to the three abundance-associated loci found in an N=1,000 dataset using flow cytometry.^37^ By comparison, in datasets of N=3,757^15^ and N=563,085,^25^ flow cytometry-based studies have identified 70 and 7,122 independent loci, respectively, associated with the abundance of predefined blood cell states. When applied to published loci, GeNA produces results with strong directional concordance to known associations (71%) (**Methods; Supp.** Fig. 24). Furthermore, power analyses suggest that GeNA’s power increases linearly with sample size for 100<N<1000 (**Methods; Supp.** Fig. 25). As new single-cell datasets and dataset integration efforts increase the sample sizes available for study, GeNA may reveal a broader catalog of csaQTLs across human tissues.

## Discussion

We have introduced a tool to flexibly detect genetic associations to the abundance of granular cell states in high-dimensional single-cell data. In transcriptomic datasets, the effects GeNA detects may be considered *trans*-eQTLs; the examples we highlight effectively reflect the association of one variant with a gene expression program composed of genes in *trans* that define a particular cell state. GeNA identifies several novel associations, highlighting the promise of high-dimensional profiling to reveal genetically associated tissue composition changes hidden in targeted approaches like flow cytometry. Further, cell type proportion GWASs are largely limited to blood, while high-dimensional single-cell profiling can be generated from many tissues. GeNA may enable, for the first time, the detection of csaQTLs in key tissue contexts where disease risk variants take effect.

GeNA has several limitations. First, phenotypes GeNA links to csaQTLs require annotation, whereas cell types predefined using flow sorting or clustering offer fixed biologic connotations. Second, the top NAM-PCs may not capture abundance variation for rare cell types. The choice of which cells are included in the dataset, such as all PBMCs versus myeloid cells only, may impact csaQTL detection. Third, GeNA uses a nearest neighbor graph embedding of single cells and the choice of embedding—which cell features are profiled and what distance metric is used—may impact the representation of cell states available for csaQTL detection. Despite these limitations, we have introduced the first tool to associate genotypes with cell states in high-dimensional single-cell data. As single-cell datasets become more widely available for genotyped cohorts, methods that leverage the rich information these data contain will be crucial to understanding genetic disease risk mechanisms.

## Acknowledgments

We are grateful to our fellow members of the Raychaudhuri Lab, as well as Yang Luo and members of the Alkes Price and Shamil Sunyaev Laboratories, for their helpful feedback. LR is supported by award F30AI157385 from the National Institute of Allergy and Infectious Diseases. JBK is supported by award F30AI172238 from the National Institute of Allergy and Infectious Diseases. LR, JBK and KAL are supported by awards numbered T32GM144273 and T32HG002295 from the National Institute of General Medical Sciences. P.-R.L. is supported by a Burroughs Wellcome Fund Career Award at the Scientific Interfaces. JEP is supported by award 1175781 from the National Health and Medical Research Council and a fellowship from the Fok Foundation. SR is supported by awards numbered R01AR063759 and UC2AR081023 from the National Institute of Arthritis and Musculoskeletal and Skin Diseases, U01HG012009 and R56HG013083 from the National Human Genome Research Institute, and P01AI148102 from the National Institute of Allergy and Infectious Diseases. The content is solely the responsibility of the authors and does not necessarily represent the official views of the National Institutes of Health. Per the agreement for the Oelen *et al.* data, we thank the participants and the staff of Lifelines-DEEP DAG2+ for their collaboration. Funding for that project was provided by the ERC Starting Grant #637640. We also acknowledge the funders of the Lifelines Cohort Study, the sample collections from which the Oelen *et al.* project data have been derived. Finally, we are grateful to all participants in the study cohorts whose data we have analyzed in this paper.

## URLs

An open-source repository containing the implementation of GeNA can be found at github.com/immunogenomics/GeNA/. All code underlying our figures and tables can be found at github.com/immunogenomics/GeNA-applied/.

## Methods

### csaQTL association testing

#### Model input

Consider a dataset of single-cell profiling for *M* cells drawn from *N* total samples, which has already undergone quality control, batch correction and neighborhood construction using CNA^36^ (i.e., we have performed a call to the function cna.tl.nam). We use CNA version 0.1.6. This function defines one neighborhood per cell in the dataset, in which many cells with similar profiles have fractional membership. The function computes an *N* × *M* matrix 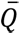, the neighborhood abundance matrix (NAM), which stores, per row, the total membership in each neighborhood across all cells from a given sample. 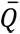 is standardized to have columns with mean zero and variance one. The function also performs principal components analysis of the NAM to yield the decomposition

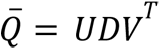

where *U* is a matrix whose *i*-th column contains the *i*-th left principal component, which has one value per sample (sample loadings on NAM-PC *i*); *D* is the diagonal matrix of singular values; and *V* is a matrix whose *i*-th column contains the *i*-th right principal component, which has one value per neighborhood (neighborhood loadings on NAM-PC *i*). GeNA obtains the input features for the csaQTL model from *U*.

#### csaQTL model

For a given SNP, let *G* be a length-*N* vector of alternative allele dose values per individual in the dataset. If we were to assess the relationship between genotype dose values *G* and sample loadings on a single NAM-PC (e.g., NAM-PC1, a length-*N* vector denoted *U*), we could use linear regression, i.e., we could model

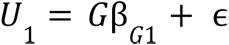

where coefficient β*G*1 reflects the relationship between *G* and *U*_1_, and ɛ represents mean-zero noise. From this model we could compute a Wald test statistic *X*_1_ that follows a chi-squared distribution with one degree of freedom under the null hypothesis, i.e.,

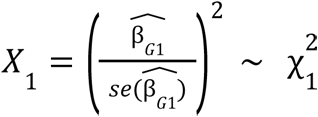

Likewise, for another NAM-PC (e.g., NAM-PC*k*, a length-*N* vector denoted *Uk*), we could quantify the relationship between genotype dose values *G* and sample loadings on NAM-PC*k* using an equivalent linear model that generates an equivalent statistic *Xk*, i.e.,

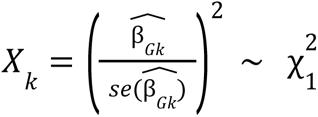

Because vectors *U*_1_,…, *U*_k_ were constructed using PCA, they are independent, i.e., *U*_1_ ⊥… ⊥ *U*_k_. For any pair of random variables *V*1 and *V*_2_ that are independent (i.e., *V*_1_⊥ *V*_2_), any measurable functions of those two random variables (e.g., *f*, *g*) are also independent, i.e., *f V*_1_ ⊥ *g* (*V*_2_), so *X*_1_ ⊥… ⊥ *X*_k_.

By the additive property of independent chi-squares, the sum of independent random variables that follow chi-squared distributions follows a chi-squared distribution with degrees of freedom equal to the sum of the degrees of freedom corresponding to the component random variables. For GeNA’s csaQTL association test, we therefore define a single test statistic *y* that incorporates all of the top NAM-PCs *U*_1_ through *U*_k_ and follows a chi-squared distribution with *k* degrees of freedom under the null hypothesis:

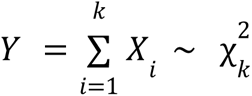

The test statistic *y* quantifies the relationship between *G* and *U*1,…, *Uk*. We can use this model to assess whether genotype values at this SNP among individuals in our cohort are associated with any linear combination of sample loadings on the top *k* NAM-PCs. To obtain an analytical p-value for this association, we compare *y* to a chi-squared distribution for *k* degrees of freedom in order to quantify the probability of a value as large as our observed *y* under the null hypothesis of no association between *G* and *U*_1_,…, *U*_*k*_. PLINK offers a computationally efficient means of obtaining Wald test statistics *X*_1_,…, *X*_k_, given quality-controlled genotype data and vectors *U*_1_,…, *U*_k_. We use PLINK v2.00a2.3LM --glm.

#### Covariates and batch effects

As facilitated by the cna.tl.nam function, GeNA accounts for covariates by residualizing them out of the NAM prior to PCA, which maintains the independence of the NAM-PCs. GeNA also residualizes batch assignments out of the NAM prior to PCA and removes any individual neighborhoods with strong abundance correlations to any batch.

#### Selection of k

By default, GeNA tests two models per SNP, one with a higher and one with a lower value of *k*, and corrects for these multiple tests. GeNA also accepts a user-specified set of *k* values. The two default values of *k* are those that offer the largest amount of variance explained in the NAM below 80% and 50% thresholds. Including two values of *k* helps to account for the possibility that some csaQTL phenotypes might be best captured by fewer NAM-PCs. GeNA reports a final p-value for each SNP that is the minimum across the tested models, adjusted using Šidák correction to account for the number of *k* values tested per SNP. When two values of *k* are tested per SNP, as in the default behavior, this correction is:

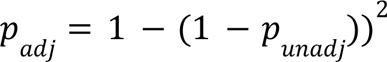

#### Defining the phenotype

For a given SNP, such as a lead SNP for a csaQTL, we apply CNA to define phenotype values for the variant-associated change in tissue cellular composition, fitting a multivariate linear regression model of the form

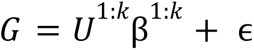

where *U*^1,k^ denotes a N-by-*k* matrix containing the sample loadings on the first *k* NAM-PCs, β^1,k^ is a length-*k* vector with one coefficient per principal component, and ɛ represents mean-zero noise. Phenotype values per sample, γ_n_, are defined using the coefficients for the fitted model, i.e.,

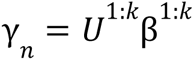

Phenotype values per neighborhood, γ_m_, are defined as the correlation per neighborhood between cell fractional abundance in that neighborhood across samples and *G*.

### OneK1K dataset processing for csaQTL GWAS

#### Dataset overview

The previously-published^1^ OneK1K dataset includes single-cell mRNA-sequencing of 1.27M peripheral blood mononuclear cells (PMBCs) collected from 982 donors of European ancestry, labeled based on 1000 Genomes and Haplotype Reference Populations, who were recruited from hospitals and retirement villages in Australia. Age and sex information is available for all participants. All samples were collected between January and April of a single year, offering approximate control for the season of blood draw as an environmental covariate. Additional clinical metadata, including self-reported clinical diagnoses and medications, is available for approximately 55% of individuals in the cohort. Genotyping for the cohort was also made available through the index publication.

#### Quality control of included individuals

Clinical metadata was shared for 1047 individuals by the dataset authors. Smoking status was one-hot encoded (N,C,P). Most other entries were coded as a binary Y/N. Rare ‘NN’ and ‘n’ entries were treated as ‘N’. Rare ‘y’ entries were treated as ‘Y’. ‘NY’ were treated as NaNs. Seven individuals designated by the dataset authors as ‘ethnic outliers’ were removed. 67 further individuals were removed because they lacked mRNA profiling. Four additional individuals were removed whose genotyping data failed quality control (see below). 969 total individuals were ultimately retained for analysis. One sample per individual was available in the scRNA-seq dataset.

#### Processing of single-cell profiling data

We began with a cells-by-genes counts matrix provided by the dataset publication authors. All cells in the dataset contained ≥660 UMIs, <7.8% mitochondrial reads, and >230 unique genes per cell. We did not modify these thresholds for cell inclusion. We removed a substantial number of additional cells identified as doublets in this dataset and reassigned remaining cells to cluster-based types following the procedure used by the original authors. We have described previously the additional doublet identification and removal steps we applied to these data.^67^ Ultimately, 822,552 total cells were retained. The median count of cells per sample was 848 (min: 201, max: 2650). We retained only genes with nonzero expression in at least three cells.

Each cell had an assignment to one of five major types—T, B, NK, myeloid cells or other—as well as an assignment to one of 30 minor types, such as CD4+ T naïve cells (a T subtype) or plasmacytoid dendritic cells (a myeloid subtype). We generated five single-cell objects from this dataset: an object including all cells that passed QC (“all cells”), as well as objects for T cells only, B cells only, NK cells only and myeloid cells only.

For each of these five data objects we used the following processing (“primary single-cell processing pipeline”): We retained only samples with at least 25 cells (N_all cells_=969 samples, N_T_=968 samples, N_NK_=935 samples, N_B_=910 samples, N_myeloid_=523 samples). We removed expression of HLA genes (“*HLA*-”) and hemoglobin genes (*HBB, HBA2, HBD, HBA1*), to avoid reference mapping biases stemming from high polymorphism, as well as cell cycle genes (S-phase and G2M-phase),^68^ to emphasize other components of expression variation across cells, and contaminating platelet genes (*PF4, PPBP*). We followed a standard scanpy pipeline^69^ to total-count normalize (library-size correct) the data to 10,000 reads per cell, to logarithmize the normalized expression values, and to select variable genes (with min_disp parameter set to yield approximately 2000 variable genes for each data object). Genes with especially high dispersion (d.var.dispersions_norm > 11) were removed. Expression values per gene across cells were scaled to give each gene unit variance. Principal components analysis was used for dimensionality reduction of the cells-by-genes matrix followed by Harmony^70^ for batch correction. The scRNA-sequencing dataset includes sample multiplexing, with 12-14 donors per library resulting in a total of 75 independent pools (“batch”). Sample assignments to pools were the batch assignments provided to Harmony. Harmony was run on the top 20 gene expression principal components with max_iter_harmony = 50 and theta = 2 for all five data objects. For the four major cell type objects (T, B, NK, myeloid), a Harmony sigma parameter value of 0.2 was used (versus the default of 0.1) to encourage softer clustering (with nclust = 50) because all cells within each of these objects belong to the same major type. The resulting harmonized principal components (“hPCs”) were used for construction of a nearest neighbor graph and UMAP using scanpy default parameters.

#### Symphony reference objects

Symphony^71^ is a single-cell reference mapping tool that assigns loadings for each cell in a query dataset within a reference dataset embedding. To enable downstream projection of phenotypes from our discovery dataset into replication datasets for replication analysis, we constructed OneK1K Symphony reference objects for each of our five single-cell data objects. Symphony typically takes as input the raw cells-by-genes expression matrix for a reference dataset as well as batch information and performs normalization, dimensionality reduction and batch correction (following the methodology of the related package Harmony), while storing intermediate objects for use in mapping a query dataset to the resulting reference embedding of hPCs. Because we had performed quality control and processing of the OneK1K dataset in Python, we assembled our Symphony reference objects by exporting the intermediate objects that Symphony requires as reference object components. Specifically, in order to build a Symphony reference object from Python-generated components, we export 1) the means and standard deviations used to normalize and scale the cells-by-variable-genes expression data, 2) loadings for each cell and each variable gene on the components that result from PCA of the cells-by-variable-genes expression matrix, 3) soft-cluster assignments produced by Harmony for each cell (in this case, the object ‘ho.R’ from harmonypy), 4) cell loadings on the batch-corrected hPCs from harmonypy, and 5) cell type assignments for each cell.

#### Genotype data processing

We obtained SNP array-based genotypes from the OneK1K study authors^1^ and performed quality control (QC) of these data. Briefly, we aligned the alleles to forward strands based on the hg19 reference genome and removed any duplicated SNPs. We retained samples with a call rate > 99% and confirmed the absence of any outlier samples for heterozygosity. We also confirmed the included individuals mapped to European ancestry using the 1000 Genomes phase 3 reference.^72^ We removed palindromic SNPs, SNPs with frequency differences >35% in comparison to the 1000 Genomes reference or with a call rate ≤98%. After QC, genotyping was available for 972 samples across 492,431 variants. These data were then phased using SHAPEIT2.^73^ We finally imputed phased genotypes by using Minimac3^74^ and the 1000 Genomes phase 3 reference. As post-imputation QC, we retained only variants with Rsq>0.7. We used PLINK to compute the top genotype PCs for use in controlling for genetic ancestry.

### Assessing performance in simulations

#### Calibration

To assess calibration (type I error), we sampled 400 SNPs from among all those on chromosome 22 that passed genotyping QC in the OneK1K dataset. Specifically, 40 SNPs were sampled at random with equal probability within each decile of MAF (above 0.05) among chromosome 22 SNPs. For each selected SNP, observed genotype values were permuted 200 times to generate simulated genotypes with no true association to the corresponding samples’ single-cell profiling. These 200 simulates per SNP were tested for csaQTLs within each single-cell data object (all cells, T, B, NK, and myeloid). In total, each of the 200 simulated genotypes per SNP is therefore included in 5 association tests. To generate an estimated overall false positive rate per SNP, we compute the number of p<0.05 results across the 200 permutations times 5 association tests for that SNP (1000 total p-values per SNP). We report the mean and standard error of these false positive rates per SNP across all 400 SNPs.

To assess calibration across cell types and *k* values, we ran GeNA again twice per object (T, B, NK, myeloid, all cells), each time directing GeNA to consider only one of the two values of *k* that GeNA selects by default for that cell type. To assess test calibration further at our smallest included MAF, we then ran an equivalent analysis using instead 400 SNPs sampled at random with equal probability among all SNPs on chromosome 22 with MAF 0.05-0.055.

#### Statistical power

To assess the statistical power of our csaQTL method we defined real cell state abundance traits that vary across individuals in the OneK1K dataset, following our previously-published methodology.^36^ To obtain traits that reflect increased expression of a gene set across all cells, we used the top 10 gene expression principal components across cells as our gene expression programs. Trait values per sample were defined as the mean loading on the selected expression PC across all cells in the sample. To obtain traits that reflect differential abundance of a cluster-based cell type, we quantified the fraction of all cells in each sample that had been assigned to a given cluster-based cell type. We included only clusters that were uncorrelated with batch (Pearson’s r^2^ < 0.25 to any batch) and that included representation of at least 50 cells from at least 100 samples. We used these same clusters to obtain traits that reflect increased expression of a gene set specifically within a given cluster-based cell type. For a given cluster, we used the top 3 gene expression principal components among cells within that cluster as our gene expression programs. Trait values per sample were defined as the mean loading on the selected cell type-specific expression PC across all cells in that cluster in the sample. We defined phenotypes using this approach for each of our five single-cell objects (all cells, B, T, NK, and myeloid), yielding 94 total phenotypes.

For each phenotype, we used the observed phenotype values per individual in the OneK1K cohort to create simulated genotypes that have a true association to the phenotype. Specifically, we generated a set of genotype values in {0, 1, 2} equal to our count of individuals with overall MAF of 0.25 and genotype frequencies consistent with Hardy-Weinberg equilibrium. We assigned the genotype values of 2 to individuals with the highest phenotype values, the genotype values of 0 to individuals with the lowest phenotype values, and genotype values of 1 to the remaining individuals.

We introduced noise to these simulated genotype values by permuting genotype values at random across individuals for a subset of individuals selected at random. For each phenotype, we permuted genotype values for 1%, 10%, 20%, 30%, 40%, 50%, 60%, 70%, 80% and 100% of samples. For each count of samples to permute, we generated 100 genotype permutations (simulates). For each simulate, we selected the given number of samples at random with equal probability and permuted genotype values among those samples at random. As the count of samples included in the permutation increases, the phenotypic variance explained by the resulting simulated SNP decays. We generated 94 x 10 x 100 = 94,000 total simulates.

We ran GeNA to conduct one csaQTL survey for associations per single-cell data object (T, B, NK, myeloid, allcells) using as input all SNPs defined to associate with true phenotypes within that data object. We ran GeNA with default parameters and controlled for age, sex, batch and gPC1-6 to generate an observed p-value for each simulate. We pooled simulates for each trait into bins based on the percent of phenotypic variance explained by genotypic variance for that simulate. Our bins corresponded to [0, 0.02, 0.04, …, 0.38, 0.4] % variance explained ±1% (i.e., the second bin contained all simulates with variance explained greater than 0.01% and less than 0.03%). To ensure that our first bin reflected true null simulates only, we included only simulated genotypes that resulted from a permutation of 100% of the samples, fully breaking the genotype-phenotype relationship. Within each bin, we estimated statistical power per trait as the fraction of association tests that resulted in a p-value less than a threshold of 5×10^-8^. We excluded from each bin any trait with fewer than 20 simulates in the corresponding range of variance explained. Finally, we computed the mean and standard error of statistical power across traits within each bin. We followed this same process to estimate statistical power for p-value thresholds of 0.05, 5×10^-5^ and 5×10^-20^.

We then subdivided our results into three phenotype categories (cell type abundance, global gene expression program, cell type-specific gene expression program) and estimated statistical power using the same process, with one modification: in order to include a power estimate for a given trait in a given variance-explained bin, we required a minimum of 7 available simulates rather than 20.

To evaluate the relationship between sample size and statistical power, we generated versions of our five OneK1K single-cell objects containing 80%, 60%, 40% and 20% of the total number of samples included in the full dataset. For each downsampled object, samples were selected for inclusion at random with equal probability from among all samples included in the full-size object. We defined traits for the downsampled objects and estimated GeNA’s statistical power to detect genetic associations to those traits following the same procedures described for the full dataset.

#### Estimated variance explained for previously-studied cell state abundance traits

As a reference point for the expected variance explained by a SNP in a cell state abundance trait we used the results published by Orrù *et al..*^15^ Specifically, we computed the 5th and 95th percentiles of ‘Heritability explained’ reported by these authors across their genome-wide significant loci (Supplementary Table 3: “Associations observed at p<5×10-8 and variant features”) for absolute count and relative abundance traits, rounded to three decimal places.

## csaQTL GWAS in the OneK1K dataset

We applied GeNA to the five single-cell objects we generated from the OneK1K dataset, controlling for age, sex, batch, and the top six genotype principal components. We use estimated alternative allele dose values per SNP, rather than best-guess genotype calls.

### Defining loci and lead SNPs

For all SNPs with associations that passed our genome-wide significance threshold (p<5×10^-8^), we sorted the SNPs in descending order by p-value. We retained the SNP with the lowest p-value as the lead SNP for the first locus and removed any other SNPs within a 1MB window centered on the lead SNP or with LD>0.8 to the lead SNP (computed using genotyping within the cohort). We then selected among the remaining SNPs the SNP with the lowest p-value as the lead SNP for the second locus, and so on. To define suggestive associations, we considered all SNPs with p-values <1×10^-6^ and used this same procedure to identify lead SNPs per suggestive csaQTL.

### Fixed-phenotype summary statistics

Many methods that take GWAS summary statistics as input assume the results per-SNP are defined in relation to a fixed phenotype (e.g., case control status for a given disease). However, the summary statistics output from GeNA can reflect different linear combinations of NAM-PCs across different SNPs (i.e., associations to different phenotypes per SNP). Therefore, to match the assumptions of available tools, we define a set of “fixed-phenotype summary statistics” for each of our csaQTLs, using per-sample values γ_n_ for the lead SNP to define associations in the local region around the lead SNP, i.e., we fit a linear model of the form

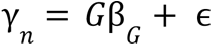

where γ_n_ is the length-*n* vector of sample-level phenotype values as defined above corresponding to the lead SNP, coefficient β_G_ reflects the relationship between *G* and γ_n_, and ɛ represents mean-zero noise. The fixed-phenotype summary statistics reflect relative differences in association across nearby SNPs to the phenotype defined using the lead SNP.

#### Cluster-based cell type proportion GWAS in the OneK1K dataset

For comparison to the csaQTL GWASs with GeNA, we ran one GWAS using cluster-based cell state abundance traits for each major cell type (T, B, NK, or myeloid) and one for all cells in the dataset. For each published cluster in the OneK1K dataset, we computed two cluster-based cell type proportion traits: one reflecting the fractional abundance of cells in the cluster relative to the corresponding major type (e.g. T, B, NK, or myeloid) and one reflecting the fractional abundance of cells in the cluster relative to all cells in the sample. We also included four traits corresponding to the fractional abundances of major cell types (T, B, NK, or myeloid) out of all cells. We removed any traits with values of 0 for 200 or more samples, which eliminated traits for rare cell types like the ASDC and pDC clusters. For any pair of traits with high correlation (Pearson’s r^2^ > 0.7) we removed one trait in order to eliminate redundancy across the tested traits. For the remaining 28 traits, we applied inverse-normal transformations, following published methodology used to test genetic associations to flow cytometry-based cell state abundance traits.^15^ We tested for associations to each trait using PLINK v1.90b6.21 (--linear), controlling for the same covariates as in the csaQTL GWASs: age, sex, and the top six genotype principal components. Within a given cluster-based GWAS (T, B, NK, myeloid, or all cells), we used a genome-wide significance threshold corrected for the total number of tested traits (5×10^-8^ / n_traits). Specifically, we used p<5×10^-9^ to account for the 10 traits in the T cell cluster GWAS, p<1.67×10^-8^ to account for the 3 traits in the myeloid cell cluster GWAS, p<1.67×10^-8^ to account for the 3 traits in the B cell cluster GWAS, p<5×10^-8^ for the single trait in the NK cell cluster GWAS, and p<4.55×10^-9^ to account for the 11 traits in the all-cells cluster GWAS. We defined our loci and lead SNPs as described for the GeNA results.

#### Characterization of GeNA csaQTLs

### Overview

To characterize a neighborhood-based association test result, whether generated by GeNA or CNA, we employ the existing toolkit of approaches used to interpret trajectories and clusters in high-dimensional single-cell data, now applied to identify biologic factors that may correspond to the observed distribution of cell abundance correlation values across neighborhoods. For example, we can identify genes whose expression correlates with the neighborhood-level phenotype, using the anchor cell for each neighborhood as an estimate for the transcriptional state represented by that neighborhood. Taking these correlation values per gene as an input ranked list, we can then apply gene set enrichment analysis to identify expression programs that characterize the cell states associated with the locus. We can also review the distribution of neighborhood-level phenotype values across cluster-based cell type labels to determine whether, for a given cell type, the abundance correlations to the locus are consistently positive (suggests expansion of that whole type), negative (suggests depletion of that whole type), or heterogenous (suggests a cell state abundance shift within that type). In some cases the phenotype may clearly correspond to an increase in expression of one gene expression program or the abundance of one particular cell type cluster. In other cases, the locus-associated phenotype may reflect multiple simultaneous changes.

### Permutation-based verification of genome-wide significant loci

We permuted the observed genotype values for each lead SNP 10^6^ times across samples and tested these permuted genotypes for association to cell state abundance shifts using the same procedure as the discovery csaQTL GWAS (in the same major cell type—NK or myeloid cells—as the observed csaQTL and with the same covariates). Our permutation p-value per lead SNP is the fraction of these permuted genotypes that attain a p-value less than the genome-wide significance threshold used in the discovery csaQTL GWAS.

### Locus Zoom plots

We applied ANNOVAR^75^ to determine the location of each SNP in a given csaQTL relative to nearby genes and to reveal any known functional consequences of the effect alleles for each SNP. Plots showing the association test results across SNPs at each locus, as well as gene locations and annotations of colocalizing QTLs, were generated with LocusZoom.^76^

### Identification of cell features that correspond to the neighborhood-level phenotype

For each csaQTL, we computed the correlation between normalized and scaled expression for each variable gene included in the single-cell object and the neighborhood-level phenotype values. We used the resulting Pearson’s r correlation values per gene as a ranked list for input to gene set enrichment analysis conducted using FGSEA^77^ and considering all the MSigDB Hallmark^78,79^ gene sets as candidates for enrichment.

For each csaQTL and corresponding enriched gene set implicating a cytokine response as part of the csaQTL-associated phenotype, we followed up with a direct association test between genotype values for the csaQTL lead SNP and estimated cytokine response levels per individual. We summed (normalized and scaled) expression per cell across all genes in the gene set that were retained among the variable genes for the single-cell data object. We then computed usage of the gene set per sample as the mean value across all cells in the sample.

We quantified the variance explained in cytokine response per sample by the lead SNP and performed a one-tailed t-test to evaluate the significance of the increase in cytokine response with increasing alternative allele dose, after controlling for age, sex, and gPC1-6.

### Testing csaQTLs with masked expression of cis-genes

For each csaQTL lead SNP and its respective single-cell object (NK or myeloid cells), we used the Gencode Release 38 assembly (mapped to GRCh37; gencode.v38lift37.annotation.gtf)^80^ to define *cis-*genes as those with any bases overlapping a two mega-base window centered on the lead SNP. We then applied identical processing as in the discovery dataset (“primary single-cell processing pipeline”) to create a *cis-*masked single-cell object, with one key change: after variable gene selection, we removed expression information for all *cis-*genes that had been included in the variable gene set for the discovery GWAS. This ensures that no information for any *cis-*gene informs the resulting data object. We then used the *cis-*gene-masked single-cell object and GeNA to test the lead SNP for a csaQTL association. We report the set of masked genes (*cis*-genes removed from the variable gene set) for each lead SNP in **Supplementary Table 10**, along with the p-value from the *cis-*masked csaQTL test.

We assessed whether expression for these *cis-*genes had corresponded strongly to the csaQTL phenotype in the discovery dataset. Specifically, for each variable gene in the discovery dataset, we computed a Pearson’s correlation r value between (normalized and scaled) expression of that gene per cell and the csaQTL phenotype value in the neighborhood anchored by that cell. For each *cis-*gene we report in **Supplementary Table 10** the resulting Pearson’s r as well as the percentile of r^2^ value for that gene among all the variable genes.

To compare the csaQTL-associated phenotypes in the discovery dataset and *cis-*masked dataset, we computed a Pearson’s correlation r value between the csaQTL per-sample phenotype values in the discovery dataset and in the *cis-*masked dataset. To compare neighborhood-level phenotypes, we quantified the consistency between the discovery and masked datasets in the cell-level features that corresponded to the neighborhood-level phenotype. Specifically, we computed a Pearson’s correlation r value between the discovery and masked datasets for the variable-genes-length vector of gene expression correlations to the neighborhood-level csaQTL phenotype.

### Testing discovered csaQTLs in custom OneK1K subcohorts

For each csaQTL, we re-ran GeNA on the lead SNP using single-cell objects which we constructed using only donors with available clinical metadata and excluding donors representing specific clinical states. For example, for the csaQTL that colocalizes with a risk association to asthma, we retained only individuals individuals with a known absence of asthma diagnosis. Sample inclusion criteria for each custom subcohort are described in **Supp. Table 9**. Using only cells from the retained samples, we constructed each custom single-cell object as described above (“primary single-cell processing pipeline”). As in the discovery analysis, we controlled for age, sex, batch, and gPC1-6 when applying GeNA to these objects. To evaluate whether a csaQTL that was evident in the relevant subcohort associated with a phenotype that was consistent with the result in the discovery dataset, we computed Pearson’s r value between the sample-level phenotypes for the discovery cohort and subcohort csaQTLs (**Supp. Table 9**).

## Colocalization of csaQTLs with molecular trait and disease risk loci

### eQTL analysis in the OneK1K dataset

For each csaQTL lead SNP, we tested for eQTL associations to all *cis*-genes within a 2MB window centered on the SNP within each single-cell object (T, B, NK, myeloid and all cells). Following existing methodology,^7,67^ we first tested genotype associations to pseudobulk gene expression per sample. For each single-cell object, we quantified pseudobulk expression for each gene in each sample. After normalizing the library size within each cell using log_2_(counts per ten thousand+1) normalization, we computed the mean normalized expression of each gene in each sample, across all cells from the sample. We retained only genes with nonzero expression in more than half of the samples. We then performed a rank-based inverse normal transformation for all genes. We used the probabilistic estimation of expression residuals (PEER) method^81,82^ (v1.0) implemented in R to infer 20 hidden determinants of pseudobulked gene expression across samples (“PEER factors”) and generate covariate-corrected expression residuals, accounting for these 20 PEER factors as well as sex, age, and gPC1-6. We used linear regression to test whether the lead SNP was associated with residualized expression for each *cis-*gene. Specifically, for each pair of csaQTL lead SNP and *cis-*gene, we fit a model

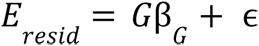

where *G* is a samples-long vector of genotype dose values at the SNP, coefficient β_G_ reflects the relationship between *G* and *Eresid*, and ɛ represents mean-zero noise. Significance was determined with Wald p-values.

For each each gene-SNP pair with a p<5×10^-4^ association in the pseudobulk eQTL model, we further tested the relationship between genotypes at the csaQTL and expression of the eGene at single-cell resolution using the Poisson mixed-effects (PME) model published by our group,^7^ implemented in the lme4^83^ (v.1.1-29) R package with parameters family = “poisson”, nAGQ=0, and control = glmerControl(optimizer = “nloptwrap”). We controlled for eight donor-level fixed-effect covariates: age, sex, and gPC1-6. We also included seven cell-level fixed-effect covariates: the natural log of the number of unique molecular identifiers (nUMI) per cell, the percent of reads per cell mapped to mitochondrial genes, and cell loadings on the top five gene expression principal components defined prior to batch correction. Age and log(nUMI) were scaled to unit variance. We also included random effects for donor (sample) and sequencing batch. Specifically, for each csaQTL lead SNP and *cis-*gene pair, we fit a full model

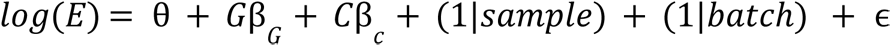

where *E* is a cells-long vector of UMI counts for the gene of interest, *G* is a vector of genotype dose values at the SNP of interest, coefficient β_G_ reflects the relationship between *G* and *log*(*E*), *C* is a cells-by-15 matrix of fixed-effect covariate values, vector β*c* reflects the relationships between the covariates in *C* and *log*(*E*), θ is an intercept, and ɛ represents mean-zero noise. This full model was compared to a null model lacking the *G*β*G* term using a likelihood ratio test, with a resulting p-value computed by comparing the resulting test statistic against a chi-squared distribution with one degree of freedom. To quantify colocalization of a csaQTL and an eQTL we defined in the OneK1K dataset, we applied the coloc^41^ R package (v5.1.0.1) with default parameters, using PP.H4.abf to quantify the posterior probability of a shared causal variant. We provided coloc with fixed-phenotype summary statistics from our csaQTL GWAS and summary statistics from our pseudobulk-based OneK1K eQTL analysis.

### Colocalization of published blood eQTLs and pQTLs to GeNA csaQTLs

To supplement our targeted eQTL analyses in the OneK1K dataset, we also reviewed summary statistics for whole-blood eQTLs published by eQTLgen^84^, eQTLs for major blood cell types published by BLUEPRINT^85^ and DICE,^86^ additional NK cell eQTLs published by Gilchrist *et al.*^87^ and Schmiedel *et al.*,^53^ and pQTLs published by Sun *et al*..^11^ We downloaded eQTLgen and DICE eQTL summary statistics directly from the DICE and eQTLgen databases, respectively. The other published summary statistics were obtained from the European Biomedical Informatics eQTL Catalogue.^88^ For each csaQTL lead SNP and reference set of summary statistics, we tested colocalization for all published loci with lead SNP p<5×10^-4^ and LD>0.8 between the csaQTL and eQTL lead SNPs. Linkage disequilibrium was calculated using the OneK1K cohort genotyping. To quantify colocalization of a csaQTL and published molecular QTL, we applied the coloc^41^ R package (v5.1.0.1) with default parameters, using PP.H4.abf to quantify the posterior probability of a shared causal variant. We provided coloc with fixed-phenotype summary statistics from the csaQTL GWAS and the published QTL summary statistics.

### Differential expression of eGene by cell subtype

We assessed differential expression of *BCL2A1* on the basis of cell membership in the CD16+ monocyte cluster using a similar Poisson-based approach. We included the same covariates as in the single-cell eQTL model above apart from cell loadings on gene expression principal components, which capture major cell state distinctions, leaving 10 included covariates. We fit a full model

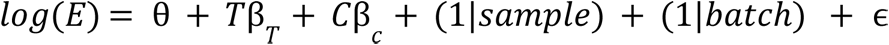

where *E* is a cells-long vector of UMI counts for the gene of interest, *T* is a vector with values 1 for all cells assigned to the type of interest (i.e., CD16+ monocytes) and 0 otherwise, coefficient β*T* reflects the relationship between *T* and *log*(*E*), *C* is a cells-by-10 matrix of fixed-effect covariate values, vector β*c* reflects the relationships between the covariates in *C* and *log*(*E*), θ is an intercept, and ɛ represents mean-zero noise. This full model was compared to a null model lacking the *T*β_*T*_ term using a likelihood ratio test, with a resulting p-value computed by comparing the resulting test statistic against a chi-squared distribution with one degree of freedom.

### Colocalization of csaQTLs and disease risk associations

Published genetic associations to disease risk that correspond to our csaQTLs were queried using the GWAS Catalog.^3^ To quantify colocalization, we applied the coloc^41^ R package (v5.1.0.1) with default parameters, using PP.H4.abf to quantify the posterior probability of a shared causal variant. We provided coloc with summary statistics for the published disease GWAS and with fixed-phenotype summary statistics from our csaQTL GWAS.

For psoriasis, we report the p=4.17×10^-9^ association identified by Tsoi *et al*. at the chromosome 12 locus in their meta-analysis (lead SNP rs11053802-T with LD 0.50 to rs3003).^43^ Summary statistics were not available from that study, so for colocalization analyses we employed summary statistics for this region available from Stuart *et al.*, which performed a psoriasis GWAS on a combined cohort of individuals with European and South Asian ancestries.^42^

#### Projection of neighborhood-based tissue cellular composition phenotypes to independent datasets

### Notation and assumptions

We previously described our methodology to define neighborhoods, compute an NAM and define NAM-PCs in the discovery dataset.^36^ To maintain notational consistency with that methodology, we use the following notation. The discovery dataset contains *N* samples and *M* cells, and therefore also *M* neighborhoods because each neighborhood in the discovery dataset is ‘anchored’ on its own discovery dataset cell. The *N* × *M* matrix 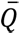 is the discovery dataset neighborhood abundance matrix (NAM), which stores the fractional density of cells from each discovery dataset sample in each discovery dataset neighborhood and is standardized to have columns with mean zero and variance one. Principal components analysis of the NAM yields the decomposition

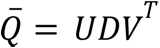

where *U* is a matrix whose *i*-th column contains the *i*-th left principal component, which has one number per sample (sample loadings on NAM-PC *i*); *D* is the diagonal matrix of singular values; and *V* is a matrix whose *i*-th column contains the *i*-th right principal component, which has one number per neighborhood (neighborhood loadings on NAM-PC *i*).

Notation for the replication dataset is differentiated by a prime marker: the replication dataset contains *N*’ samples and *M*’ cells from which we will construct an NAM 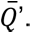’. Importantly, because we seek to define the distribution of replication dataset samples over discovery dataset neighborhoods, 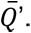’ will have dimensions *N*’ × *M*.

Finally, let *y* be a length-*N* vector containing the attribute values (e.g., allele dose or case-control status) per sample and let *k* be the number of NAM-PCs included in the discovery dataset association test that defined the phenotype of interest, i.e., when we fit a model

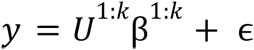

where *U*^1,k^ denotes the first *k* columns of *U*, β^1,k^ is a length-*k* vector with one coefficient per principal component, and ɛ represents mean-zero noise.

### Membership of each replication dataset cell among discovery dataset neighborhoods

We use our published reference mapping algorithm Symphony^71^ to generate a reference object for the discovery dataset (see “*Making symphony reference objects*” above) and situate each replication dataset cell within the discovery dataset embedding (“mapQuery” function). Symphony requires raw or normalized values per cell in the replication dataset for the same set of cell features (e.g., transcript counts per gene) as were used to define the discovery dataset embedding. Symphony performs batch correction for user-specified covariates in the replication dataset. Quality control of the replication dataset cells and samples should be performed prior to Symphony mapping. For each replication dataset cell now situated in the discovery dataset embedding, we use the nn2 (Nearest Neighbour Search) function from the RANN package, an R wrapper for Arya and Mount’s Approximate Nearest Neighbours (ANN) C++ library,^89^ to identify the 15 nearest discovery dataset cells and compute the similarity (1/distance) of these 15 cells to the replication dataset cell. These 15 closest neighbors anchor the neighborhoods in which this replication dataset cell will be assigned non-zero initial membership, with fractional membership proportional to the degree of similarity.

Let *A* be an *M*’ × *M* matrix representing the initial membership of each replication dataset cell *m*’ in each discovery dataset neighborhood *m*. If the anchor cell for *m* was one of the 15 nearest neighbors to *m*’, then the value stored at the *m*’-th row and *m*-th column of *A* is the similarity between *m*’ and *m*. All other values of *A* are zero. We normalize the rows of *A* to sum to one to give each cell equal influence, i.e.,

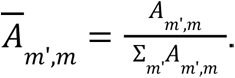

### Cell density distributions per replication sample across discovery dataset neighborhoods

For every sample *n*’ among the total *N*’ replication samples let *C*(*n*’) be the set of cells belonging to the *n*’-th sample. We then define an *N*’ × *M* matrix by taking the sum of cell density per neighborhood across all cells from a given replication sample, i.e.,

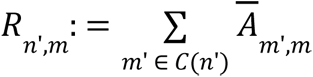

We normalize the rows of *R* to sum to one, i.e.,

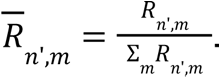

Following methodology we previously described and published in CNA,^36^ we can then diffuse the distribution of fractional abundance for each sample across neighborhoods using pairwise neighborhood similarity values stored in the discovery dataset nearest neighbor graph. More specifically, we apply the cna.tl.diffuse_stepwise function with identical stopping criteria as in CNA. After diffusion, most neighborhoods will have some degree of representation from most samples. The rows of the resulting *N*’ × *M* matrix *Q*’ are normalized to each sum to one, yielding the NAM for replication samples, 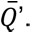’.

### Defining a phenotype value per replication dataset sample

To obtain *U*’^1:*k*^, the loadings for the replication samples on the first *k* discovery dataset NAM-PCs, we take the dot product of the replication dataset NAM, the loadings of the discovery dataset neighborhoods on the first *k* NAM-PCs and the diagonal matrix of *k* singular values from the discovery dataset, i.e.,

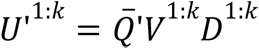

Finally, to obtain estimated phenotype values for each replication sample, 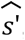, we combine information about sample loadings across the *k* NAM-PCs, weighted according to the coefficient fitted values that defined the phenotype in the discovery dataset, i.e.,

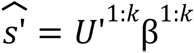

#### Testing for an attribute-phenotype association in the replication dataset

In order to evaluate whether the projected phenotype values per replication sample, 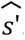, associate with *y*’, the sample attribute of interest (e.g., allele dose or case-control status per sample) in the replication dataset, we can fit a linear model, i.e.,

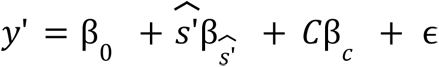

where β_0_ is an intercept, 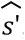 is a length-*N*’ vector of projected phenotype values per replication dataset sample, β*C* is a vector with one coefficient for each covariate (e.g., age, sex), *C* is a *N*’ × number of covariates-dimensional matrix storing covariate values per replication sample, and ɛ represents mean-zero noise. The coefficient 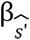 represents the relationship between the ground truth attribute values per sample in the replication dataset (*y*’) and the projected phenotype values per replication dataset sample 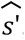. We use a one-tailed Student’s t-test to evaluate whether 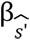 > 0, which would indicate that samples in the replication dataset with higher attribute values also tend to have a stronger presence of the projected phenotype in their single-cell profiled tissue.

#### Processing of Perez *et al.* dataset

Perez *et al.*^38^ generated single-cell profiling and genotype data from N=261 individuals, including 162 donors with systemic lupus erythematosus (SLE) and 99 donors without SLE.

#### Genotype quality control and imputation

Genotyping data were downloaded from DBGaP (accession phs002812.v1.p1). We performed imputation separately for 49 ImmVar samples assayed with the Omni Array and the remaining 209 samples assayed with the LAT Array. This cohort split by genotyping array was revealed by the genotyped SNPs available for each sample. We excluded five samples with self-reported ancestries other than European and Asian. Using PLINK within each genotyping cohort, assayed variants with <1% missingness across samples were retained (696,384 SNPs for LAT array and 766,172 SNPs for Omni), after which all samples had <1.5% missingness across variants. No duplicate SNPs or SNPs with reversed or ambiguous strand orientation relative to the hg19 reference downloaded from UCSC (chromFa.tar.gz) were present. Only SNPs with MAF>1% were retained (585,456 SNPs for LAT array and 610,590 SNPs for Omni). Sample heterozygosity was computed with PLINK and two Omni cohort samples were removed with heterozygosity greater than three standard deviations above the cohort mean. Variants were removed from the Omni Array cohort (375 variants) and LAT Array cohort (226 variants) with observed allele frequencies discordant with the multipopulation average allele frequency among 1000 Genomes Project samples.

Visual examination of a joint PCA plot affirmed that Omni and LAT array samples colocated by self-reported ancestry (European or Asian) to the corresponding major ancestral population among 1000 Genomes Project samples. For each genotyping cohort, these joint principal components were generated after retaining only SNPs included in the 1000 Genomes Project data with MAF>5%, missingness <5%, and pruned to approximate linkage equilibrium (PLINK parameter --indep 50 5 2). Identity by descent (IBD) was computed pairwise for samples within each self-reported ancestry group within each genotyping cohort using PLINK. Seven samples were identified to have high relatedness (PI_HAT>0.125) with at least one other sample in the dataset. We removed the minimum set of samples necessary to eliminate at least one from each pair of samples with high IBD. When either one of two samples could be removed to attain this outcome, we retained the sample with lower genotype missingness. In total, four samples were removed from the LAT Array cohort based on IBD (2 of Asian ancestry, 2 of European), leaving 200 total samples from the LAT array (103 of Asian ancestry and 97 of European) and 47 individuals of European ancestry genotyped on the Omni array. The resulting post-QC genotype data was used for imputation. By chromosome and within genotyping cohort, phasing was completed using SHAPEIT version v2.r727,^90^ and imputation was completed using Minimac3.^74^

### Genotype PCs

Genotype PCs were computed separately with PLINK for the Asian (LAT array only) and European (Omni and LAT arrays) cohorts using post-QC but pre-imputation genotypes, after retaining only SNPs with MAF>5% pruned to approximate linkage equilibrium (PLINK parameter --indep 50 5 2). For the European cohort, SNPs shared between the LAT and Omni arrays (148,945 SNPs, before filtration) were used. Within the European cohort, gPCs capture within-ancestry genotypic variation rather than array batch (**Supp.** Fig. 26).

### Constructing two ancestry-based cohorts

Samples of Asian ancestry (N=103), all of which were genotyped on the LAT array, were isolated after imputation. Only SNPs with MAF>1% within this cohort and R2>0.7 from LAT array imputation were retained. To create the second ancestry cohort (European), post-imputation SNPs shared across both arrays were retained, samples of Asian ancestry were excluded, and data from the remaining samples was merged (N=144). SNPs were not filtered on the basis of MAF or R2 within the replication cohort, lest lead SNPs from the discovery dataset be eliminated. Instead, replication was tested for all lead SNPs from the discovery dataset, and the MAF and R2 for these SNPs in the replication cohort was reviewed.

### Single-cell mRNA-seq data sourcing and sample quality control

Post-quality-control single-cell transcriptomic profiling was downloaded from GEO (accession: GSE174188). The dataset was subdivided by ancestry cohort, as defined above. Samples from individuals that failed genotype QC were excluded. Samples from SLE patients with a status of “treated” or “flare” were removed, leaving only SLE samples of the status “managed.” The Perez *et al.* dataset includes several donors with duplicate samples and includes 23 sample batches, some of which have very few samples. We removed sample duplicates—to yield one sample per donor—while maximizing the average number of samples per retained batch.

### Single-cell objects for csaQTL replication testing

After sample-level quality control, we retained only cells with expression of at least 200 genes and <10% mitochondrial reads. We generated one single-cell object for NK cells and another for Myeloid cells, using the cell type labels (“cg_cov”) provided by the study authors, separately for the European ancestry and Asian ancestry cohorts, yielding four single-cell objects total. For each single-cell object, we retained only genes expressed in at least three cells. We used Gencode version 19 to map the available gene names to EnsembleIDs. We provided both “batch_cov” and “Processing_Cohort” (nested batch variables from GEO object) for Symphony batch correction, along with SLE case-control status and sex. Because European ancestry individuals were genotyped on one of two arrays, while all Asian ancestry individuals were genotyped on the same array, we also provided a genotype array covariate for Symphony correction within the European ancestry single-cell objects.

### Covariates for csaQTL replication testing

In our linear models to test associations between a csaQTL lead SNP and a phenotype projected from the discovery dataset into each replication cohort, we included covariates for age, sex, and genotype PCs 1-4. For the European ancestry cohort we also included a genotyping array covariate. The number of gPCs to include was selected using an elbow plot for each cohort (**Supp.** Figs. 26-27).

### Single-cell objects for sex-associated phenotype projection-replication example

We followed the same process as just described to generate NK, Myeloid, B, T, and all cells single-cell objects for both the Asian ancestry and European ancestry cohorts (ten objects total) with one exception: because we sought to evaluate replication for a sex-associated phenotype we did not control for sex during Symphony batch correction or association replication testing.

#### Processing of Oelen *et al.* dataset

### Overview

Oelen *et al.*^40,91^ generated single-cell mRNA-sequencing of 1.3 million PBMCs from N=120 healthy individuals of European ancestry in the Lifelines Cohort Study for whom genotype data had previously been made available by Tigchelaar *et al.*.^91^ The authors generated multiple samples from each donor in culture. For each individual in the Oelen *et al.* cohort, the study authors profiled cells from an untreated sample as well as samples exposed to pathogens at each of two timepoints.

### Genotype data for csaQTL replication testing

Genotyping data was obtained from the University Medical Center Groningen under a Lifelines DEEP DAG2+ Project Data Access Agreement, including 7,249,882 variants genome wide mapped on the GRCh37 reference genome. Genotypes with a posterior probability (GP) <0.9 were considered missing. All of our variants used for replication testing had a missing call rate ≤10% and MAF ≥5%. We used rs55908509 as proxy for the chromosome 19 csaQTL lead SNP because these two variants are in complete LD (1.0) in European-ancestry populations and genotypes for the lead SNP itself were not available. Genotype PCs were computed with PLINK after retaining only SNPs with MAF>5% and 95% genotyping rate (--geno 0.05) pruned to approximate linkage equilibrium (--indep 50 5 2).

### Single-cell objects for csaQTL replication testing

Demultiplexed and doublet-QC’ed single-cell mRNA count files were obtained via the European Genome-Phenome Archive under a Lifelines DEEP DAG2+ Project Data Access Agreement. We retained only untreated (“UT”) samples, of which one was available for each of 104 donors. We used Gencode version 29 to map the available gene names to EnsembleIDs. We retained only cells with expression of at least 200 genes and <10% mitochondrial reads. We used the provided cell type labels to generate one single-cell object for NK cells and another for Myeloid cells. For each single-cell object, we retained only samples with at least 25 cells (N_NK_=103 samples, N_myeloid_=104 samples) and we retained only genes expressed in at least three cells. We provided both “batch” and “chem” (10x Genomics v2 versus v3 chemistry reagents) for Symphony batch correction.

### Covariates for csaQTL replication testing

In our linear models to test associations between a csaQTL lead SNP and a phenotype projected from the discovery dataset into this cohort, we included covariates for age, sex, and genotype PCs 1-5. The number of gPCs to include was selected using an elbow plot (**Supp.** Fig. 28).

#### Processing of Randolph *et al.* dataset

### Overview

Randolph *et al.*^39^ generated single-cell mRNA-sequencing of 255,731 PBMCs from N=90 healthy male donors of either European (N=45) ancestry or African (N=45) ancestry, along with genotype data. In the published study, the authors generated two samples from each donor, exposed one sample to influenza virus and exposed the other sample to mock-condition media as a negative control. Samples were frozen after collection and before infection or mock infection exposure and profiling. Cells were profiled from all samples after six hours of exposure.

### Genotype data for csaQTL replication testing

Genotyping data was obtained through the Sequence Read Archive (accession PRJNA736483). Low-pass whole genome sequencing was available for 89 donors at a total of 78,111,311 variants genome wide. Genotypes with a posterior probability (GP) <0.9 were considered missing. All of our variants used for replication testing had a missing call rate ≤10% and MAF ≥5%. Genotype PCs for each ancestry cohort were computed with PLINK after retaining only SNPs with MAF>5% and 95% genotyping rate (--geno 0.05) pruned to approximate linkage equilibrium (--indep 50 5 2).

### Single-cell objects for csaQTL replication testing

Demultiplexed and doublet-QC’ed mRNA-seq count files were obtained through GEO (accession GSE162632). We retained only control samples (SOC_infection_status==”NI”) for donors with genotyping data available. We used Gencode version 29 to map the available gene names to EnsembleIDs. We retained only cells with expression of at least 200 genes and <10% mitochondrial reads. For each ancestry cohort, we used the provided cell type labels to generate one single-cell object for NK cells and another for Myeloid cells. For each single-cell object, we retained only samples with at least 25 cells (N_NK, EUR_=41 samples, N_myeloid, EUR_=44 samples, N_NK, AFR_=38 samples, N_myeloid, AFR_=42 samples) and we retained only genes expressed in at least three cells. We provided batch assignments (“batchID”) to Symphony for batch effect correction.

### Covariates for csaQTL replication testing

In our linear models to test associations between a csaQTL lead SNP and a phenotype projected from the discovery dataset into these cohorts, we included covariates for age and the top genotype PCs within the given cohort. A sex covariate was not included because all donors were male. For the European ancestry cohort we included gPCs 1-5. For the African ancestry cohort we included gPCs 1-2. The number of gPCs to include was selected using an elbow plot for each cohort (**Supp.** Figs. 29-30).

#### Replication testing of csaQTLs across five cohorts

### Replication testing per cohort

We evaluated replication of the five csaQTLs detected by GeNA in five cohorts with PBMC scRNA-sequencing: one Asian ancestry (N=103) and one European ancestry cohort (N=144) from Perez *et al.*,^38^ one European ancestry (N=41) and one African ancestry cohort (N=38) from Randolph *et al.*,^39^ and one European ancestry cohort (N=103) from Oelen *et al.*.^40,91^ Data acquisition and single-cell dataset processing of these cohorts is described above. For each csaQTL, we projected the csaQTL phenotype into a corresponding dataset for each cohort (i.e., Myeloid cells for the csaQTL on chromosome 15, and NK cells for the other loci). We tested the projected phenotype for association to allele dose per sample for the corresponding lead SNP, also as described above (“*Testing for an attribute-phenotype association in the replication dataset”*). We excluded cohorts on a per-csaQTL basis if genotype information was not available for the lead SNP (e.g., did not pass genotype QC) or all samples had equal allele values. Across all cohorts, when testing replication we controlled for age, sex, and top genotype PCs, along with dataset-specific covariates where relevant. Specific covariates included for each cohort are described above in the “Processing” sections per dataset.

### Meta-analysis of csaQTL replication across cohorts

We used inverse-variance weighted meta-analysis to combine results across cohorts for each csaQTL, with a one-tailed Student’s t-test to evaluate the hypothesis that the meta-analysis effect size is greater than zero (i.e., significant and directionally concordant to the effect in the discovery dataset).

#### Polygenic risk score analyses

### Selection of diseases

Perez *et al.* represents the first large case-control study for an autoimmune disease—systemic lupus erythematosus (SLE)—with both single-cell profiling of peripheral blood and genotyping data available. RA is the one autoimmune disease in the OneK1K cohort with sufficient representation (N>10) to enable a disease case-control analysis for direct comparison of peripheral blood cell states associated in abundance with higher RA genetic risk to peripheral blood cell states that are differentially abundant in patients with RA relative to controls.

### Defining custom OneK1K subcohorts

The OneK1K cohort includes a cross-section of clinical states. To avoid PRS values acting as a proxy for clinically documented disease status when we test for cell state abundance differences associated with PRS values, we included PRS values only for individuals with documented clinical metadata and a known absence of the disease for which the PRS was constructed (OneK1K_PRS: SLE_ and OneK1K_PRS: RA_ cohorts). We also defined a OneK1K subcohort including only individuals with a known absence of any autoimmune disease (OneK1K_PRS: ADs_). The resulting sample sizes for each single-cell object are reported in **Supp. Table 11**.

### Selection of PRSs

Published and validated polygenic risk scores were obtained from the PGS Catalog.^92^ Among the available SLE PRSs in the PGS Catalog developed and validated in European-ancestry cohorts, we selected the study with the largest source population: Chen et al. “Main” PRS, PGS ID PGS000771.^59^ Likewise, among the available RA PRSs in the PGS Catalog developed and validated in European-ancestry cohorts, we selected the study with the largest source population: Privé et al. penalized regression model, PGS ID PGS001875.^66^ Among the 95 total variants in the SLE PRS, 87 were available in our QC’ed genotype data (92%). Among the 256 total variants in the RA PRS, 251 were available in our QC’ed genotype data (98%). We removed all MHC variants (defined as variants between positions 28477797 and 33448354 on chromosome 6, per the NIH Genome Reference Consortium)^93^ from each PRS. In the SLE PRS, 86 variants remained after removal of 1 MHC SNP. In the RA PRS, 190 variants remained after removal of 61 MHC variants. We computed the correlation between PRS value per-person for each disease and each of the 29 available clinical variables (diagnoses and medications) and found that the PRSs were not correlated with any known clinical feature within the OneK1K_PRS:RA_, OneK1K_PRS:SLE_, or OneK1K_PRS:ADs_ cohorts (Pearson’s r^2^ < 0.01 for all).

### Association testing to PRS values in OneK1K_PRS:RA_ and OneK1K_PRS:SLE_

We tested for associations to each disease PRS within each major celltype (all cells, T, NK, B, and myeloid) for the corresponding cohort (OneK1K_PRS:RA_ or OneK1K_PRS:SLE_) using CNA while controlling for batch, age, sex, and gPC1-6, and with with the “ks” parameter value vector defined as above (using thresholds for 50% and 80% of variance explained). We used the Benjamini-Hochberg method of FDR correction to account for multiple hypothesis testing across cell types for each disease PRS. For associations that passed a nominal significance threshold of p<0.05, we tested the PRS for association in OneK1K_PRS:ADs_.

### SLE case-control analysis

To define lupus case-control differences within our neighborhood-based framework, we used the single-cell profiling and genotype data from the Perez *et al*. European cohort. Using the published assignments of cells to cluster-based minor cell types (“cg-cov” attribute in the GEO data object), we assigned each cell to one of four major types: T, B, NK, or myeloid. We generated four single-cell data objects, each containing cells from one of these major types, following our “primary single-cell processing pipeline” as described above with the following modifications: We retained only cells with expression of at least 200 genes and <10% mitochondrial reads. We excluded cell cycle genes but retained HLA, hemoglobin and platelet genes. We provided both “batch_cov” and “Processing_Cohort” (nested batch variables) for Harmony correction.

We defined lupus case-control differences in cell state abundance across neighborhoods using CNA for each single-cell object, controlling for batch (“batch_cov”), age and sex. We then computed correlations in expression per variable gene to the neighborhood-level SLE-associated phenotype for each major cell type. We observed that in all four major cell types *ISG15* and *IFI44L* were among the top 2-7 genes most positively correlated in expression with the SLE phenotype across neighborhoods. We summed the (normalized and scaled) expression of these two genes per neighborhood and computed the correlation of this interferon signature to the SLE phenotype. We computed a p-value for whether this correlation was significantly greater than zero by bootstrapping over samples. We then computed the mean value (across cells) per sample of this interferon signature and re-tested the SLE case-control analysis using CNA with the addition of this interferon response covariate.

### Interpretation of SLE PRS-associated phenotype

To characterize the cell state abundance shift associated with increasing SLE PRS value, we computed the correlation across neighborhoods between the per-neighborhood SLE PRS phenotype and expression for each variable gene. These values were used as a ranked list for input to gene set enrichment analysis with FGSEA in which we tested all MSigDB Hallmark gene sets. The interferon alpha response gene set was identified as the top enriched gene set. We summed the (normalized and scaled) expression of all genes in this gene set available among the variable genes of the myeloid OneK1K_PRS: SLE_ data object. We computed the correlation between these interferon-alpha signature values and the neighborhood-level SLE PRS phenotype as well as a corresponding p-value for whether this correlation was significantly greater than zero using bootstrapping over samples.

To evaluate whether the PRS association was driven by a strong effect from one SNP or aggregated similar but small effects across multiple SNPs, we tested associations for each SLE PRS SNP within the Myeloid OneK1K_PRS: SLE_ object using CNA while controlling for batch, age, sex, and gPC1-6, and with with the “ks” parameter value vector defined as above (using thresholds for 50% and 80% of variance explained). We used the Benjamini-Hochberg procedure to control the false discovery rate across SNPs. For each SNP, we computed a Pearson’s r correlation between the per-neighborhood abundance correlations to that SNP and the interferon-alpha response expression signature. We used the GWAS Catalog^3^ to identify mapped genes for each SNPs shown in **Fig. 4**.

### OneK1K_CC: RA_ dataset and RA case-control association test

A total of N=16 OneK1K participants were labeled with a diagnosis of RA. To ensure that our control individuals had a known absence of RA, we removed all individuals who lacked clinical metadata. We also removed those individuals with any autoimmune disease besides RA. From the remaining control candidates, we sampled at random with equal probability a group equal in size to the RA cohort (**Supp. Table 12**). We otherwise followed our “primary single-cell processing pipeline” as described above to generate single-cell objects per major cell type (T, B, NK, myeloid) for the RA case-control association tests. We applied CNA to test for case-control differences, controlling for age, sex and batch, and with the “ks” parameter value vector defined as above (using thresholds for 50% and 80% of variance explained).

### Interpretation of RA-associated and RA PRS-associated cell state abundance phenotypes

To interpret the RA PRS and RA disease associated phenotypes, we reviewed the distribution of per-neighborhood phenotype values within each of the published cluster-based cell types. For the type-based shift suggested by the CNA result (i.e., CD4+ T_naïve_/[T_EM_+T_CTL_] ratio), we performed a follow-up direct association test. To test an association between RA case-control status and a cluster-based trait, we used a one-sided t-test to evaluate a difference in trait values among RA and control patients specifically in the direction suggested by CNA. To test an association between a cluster-based trait and RA PRS value, we used permutation across samples, which are independent in our dataset unlike values across neighborhoods. In other words, we computed the CD4+ T_naïve_/[T_EM_+T_CTL_] ratio per individual and the observed Pearson’s r correlation between these trait values and the RA PRS values. Then we permuted the PRS values across samples 1000 times and defined our permutation p-value as the fraction of those 1000 trials in which the trait-PRS correlation was more extreme than the observed value.

#### Comparison to results from previous flow cytometry studies

To our knowledge, Orrù *et al.* conducted the largest blood cell state abundance GWAS by trait count (310 cell state abundance traits; N=3,757 Sardinian participants; 70 found loci). We therefore chose this study as our reference for known blood cell type proportion trait genetic associations. We used Supplementary Table 1B (“Characterization of immunophenotypes: Immune traits measured”) to quantify the number of cell state abundance traits studied by Orrù *et al.*. We considered traits assigned by the study authors to the categories Absolute Count and Relative Count to be cell state abundance traits and excluded MFI and morphology traits. We used Supplementary Table 3 (“Associations observed at p<5×10-8 and variant features”) as our reference set of associations from Orrù *et al*.

Orrù *et al.* had defined one set of independent loci per trait and reported their total set of loci as the union of those loci. We further pruned these loci to define a single set of independent loci across all traits. We first excluded any loci for which genotyping was not available in the OneK1K cohort. Then, we saved the locus with the most significant association and removed other loci in a 2 megabase window centered on the retained SNP. We then saved the locus, among the remaining loci, with the next most significant association and so on.

We determined that we could estimate values per person in the OneK1K cohort for 53 cell state abundance traits tested by Orrù *et al*. Specifically, we included phenotypes that could be defined based on cell counts in the available clusters. For example, we included Orrù trait “T cell %lymphocyte,” which we defined per OneK1K participant as the count of cells in T cell clusters relative to the total count of cells across T, B, and NK clusters. We assigned each of these phenotypes to the appropriate one of our five single-cell objects. For example, the “T cell %lymphocyte” trait was assigned to the all-cells object, while Orrù’s “DN (CD4-CD8-) %T cell” phenotype was assigned to the T cell object. We retained only phenotypes that could be captured by the NAM-PCs. Specifically, for each phenotype we fit a model

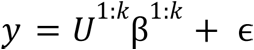

where *y* denotes a length-N vector of the cluster-based true phenotype values per individual, *U* denotes a N-by-*k* matrix containing the sample loadings on the first *k* NAM-PCs, β is a length-*k* vector with one coefficient per principal component, and ɛ represents mean-zero noise.

GeNA identifies for each tested SNP the phenotype most associated with alternative allele dose, which may be some combination or extension of the predefined traits previously found to be associated with this SNP. To ensure consistency in our comparison, for each known locus we defined a phenotype per sample as the linear combination of NAM-PCs that best approximates the previously studied trait. We then tested SNP associations to this fixed phenotype to evaluate directional concordance between associations detectable by our neighborhood-based framework and the known associations. To maintain consistency with our csaQTL GWAS, we selected *k* using an 80% of variance explained threshold (the larger of two *k* values used by GeNA by default). We then defined

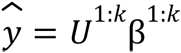

as the closest approximation of the true trait available as a linear combination of the NAM-PCs, and we retained 36 phenotypes with Pearson’s r^2^ ≥ 0.6 between *y* and *y*. Traits that were not retained tended to reflect small cell populations. For example, for Orrù trait “TCRgd %lymphocyte” the 75th percentile of phenotype values was 2%. This trait can only be defined in the all-cells single cell object but γδ T cells represent a small population that does not strongly influence total PBMC cell state abundance variation across individuals and is therefore not well captured by the top *k* NAM-PCs. For each retained Orrù trait, we used PLINK to test associations between *y* and the SNPs found by Orrù to achieve genome-wide significance in association to *y*, i.e.,

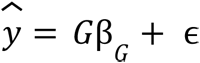

Where *G* is an N-dimensional vector of effect allele dose values per individual, β*G* is our effect estimate of interest for comparison, and ɛ represents mean-zero noise. We controlled for age, sex, and gPC1-6. Fixing 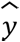 enables us to assess whether the resulting genotype associations to 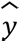 are concordant with past flow cytometry findings.

