## Supplement for "Identifying genetic variants that influence the abundance of cell states in single-cell data"

#### Contents

**Supplementary Tables** **2**

**Supplementary Figures** **8**

**Supplementary Note** **30**

### Supplementary Tables

| Cell Type | CHR | POS | rsID | Effect Allele | Other Allele | Cyto. Band | MAF | P |
|---|---|---|---|---|---|---|---|---|
| Myeloid | 15 | 80263217 | rs3826007 | T | C | 15q25.1 | 0.27 | 2.61e-08 |
| NK | 2 | 111851212 | rs13025330 | T | C | 2q13 | 0.22 | 1.76e-09 |
| NK | 11 | 128070535 | rs519062 | G | A | 11q24.3 | 0.28 | 2.48e-13 |
| NK | 12 | 10583611 | rs3003 | T | C | 12p13.2 | 0.18 | 1.96e-11 |
| NK | 19 | 16441973 | rs56133626 | A | G | 19p13.11 | 0.33 | 1.96e-13 |

**Supplementary Table 1: Genome-wide significant csaQTLs in the OneK1K dataset detected by GeNA**. For each csaQTL, we indicate the GRCh37 build position and rsID of the lead SNP, the effect allele associated with the observed phenotype, the cytogenetic band of the locus as well as the MAF and GeNA p-value.

| Trait | CHR | BP | Effect Allele | P | BETA |
|---|---|---|---|---|---|
| CD16 Mono_frac_Myeloid | 15 | 80267501 | G | 5.732000e-10 | -0.4257 |
| CD16 Mono_frac_allcells | 15 | 80260275 | T | 9.171000e-11 | -0.3212 |
| NK_CD56bright_frac_NK | 2 | 111832065 | A | 3.257000e-11 | -0.3208 |
| NK_CD56bright_frac_NK | 12 | 52595174 | T | 9.888000e-09 | 0.2649 |
| NK_CD56bright_frac_allcells | 2 | 111836333 | G | 7.359000e-11 | -0.3377 |

**Supplementary Table 2: Lead SNPs for csaQTLs that pass a genome-wide significant threshold in a cluster-based GWAS approach performed for comparison against GeNA**. For each cell state cluster defined by Yazar, Alquicira-Hernandez, Wing, *et al.* in the OneK1K dataset, we defined two traits: the fractional abundance of cells in that type out of all cells in the sample (e.g., CD4+ effector memory T cells % of all cells) and the fractional abundance of cells in that type out of all cells in the corresponding major cell type (e.g., CD4+ effector memory T cells % of all T cells). We performed quality control of these traits (e.g., to remove one from each strongly correlated trait pair) and performed one GWAS for each major cell type including all traits defined within that type. For each lead SNP, we list the associated trait and effect allele as well as the chromosome and the GRCh37 build position of the lead SNP.

| Cell Type | CHR | Phenotype Annotation | Novel |
|---|---|---|---|
| Myeloid | 15 | Decrease in CD16+ monocytes % myeloid cells | No |
| NK | 2 | Decrease in CD56bright % NK cells | No |
| NK | 11 | Increase in NK cells activated by TNF-$\alpha$ and IFN-$\gamma$ % NK cells | Yes |
| NK | 12 | Increase in NK cells activated by TNF-$\alpha$, IFN-$\gamma$, IL-2, and IL-6 % NK cells | Yes |
| NK | 19 | Increase in NK cells activated by TNF-$\alpha$ % NK cells | Yes |

**Supplementary Table 3: Phenotypes associated with dose of alternative allele at each csaQTL detected by GeNA**. For each csaQTL, we summarize the phenotype observed in association with higher dose of the effect allele. The analyses that form the basis of these phenotype annotations are discussed below. A directionally-concordant genotype association to the listed phenotype was observed by Orrù *et al.*[1] for the csaQTL at 15q25.1 and a directionally-concordant genotype association to the listed phenotype was observed by Patin *et al.*[2] for the csaQTL at 2q13. Specifically, Orrù *et al.* identified the same lead SNP, rs3826007-T, to associate (p=5x$10^{-9}$) with a decrease in "CD14- CD16+ monocyte %monocyte." Patin *et al.* identified rs12986962-G to associate (p=9x$10^{-19}$) with decreased abundance of CD8a+CD56hi NK cells. rs12986962-G is in LD (0.3) and correlates with rs13025330-T, the GeNA lead SNP.

| Publication | Cohort | N, NK dataset | N, Myeloid dataset | Cells, NK dataset | Cells, Myeloid dataset |
|---|---|---|---|---|---|
| Randolph et al. | AFR | 38 | 42 | 2544 | 6055 |
| Perez et al. | ASI | 98 | 98 | 23817 | 115912 |
| Randolph et al. | EUR | 41 | 44 | 3687 | 7988 |
| Oelen et al. | EUR | 103 | 104 | 16731 | 22441 |
| Perez et al. | EUR | 139 | 140 | 40201 | 154527 |

**Supplementary Table 4: Five cohorts from three previously published datastes of scRNA-seq profiling of PBMCs were used to test replication of the novel csaQTLs identified by GeNA in the OneK1K dataset**. We tabulate the replication cohorts here, indicating the count of donors (**N**) and **Cells** that passed quality control.

| csaQTL | Pathway | P-adjusted |
|---|---|---|
| 11:128070535:A:G | HALLMARK_TNFA_SIGNALING_VIA_NFKB | 8.4e-07 |
| 11:128070535:A:G | HALLMARK_INTERFERON_GAMMA_RESPONSE | 0.04 |
| 12:10583611:C:T | HALLMARK_TNFA_SIGNALING_VIA_NFKB | 5.2e-12 |
| 12:10583611:C:T | HALLMARK_HYPOXIA | 2.3e-06 |
| 12:10583611:C:T | HALLMARK_MTORC1_SIGNALING | 4.4e-03 |
| 12:10583611:C:T | HALLMARK_IL2_STAT5_SIGNALING | 5.8e-03 |
| 12:10583611:C:T | HALLMARK_INFLAMMATORY_RESPONSE | 6.7e-03 |
| 12:10583611:C:T | HALLMARK_INTERFERON_GAMMA_RESPONSE | 6.7e-03 |
| 12:10583611:C:T | HALLMARK_ESTROGEN_RESPONSE_LATE | 0.012 |
| 12:10583611:C:T | HALLMARK_IL6_JAK_STAT3_SIGNALING | 0.012 |
| 12:10583611:C:T | HALLMARK_P53_PATHWAY | 0.013 |
| 12:10583611:C:T | HALLMARK_KRAS_SIGNALING_UP | 0.015 |
| 19:16441973:G:A | HALLMARK_TNFA_SIGNALING_VIA_NFKB | 3.5e-08 |

**Supplementary Table 5: All MSigDB Hallmark gene sets significantly enriched in the cell states associated with three novel csaQTLs revealed by GeNA**. For each novel csaQTL, we list all Hallmark gene sets that passed a p<0.05 threshold after correction for multiple testing with Benjamini-Hochberg adjustment.

| csaQTL | Stimulus | r-squared | P |
|---|---|---|---|
| 12:10583611:C:T | TNF-$\alpha$ | 0.008370 | 0.006119 |
| 12:10583611:C:T | IFN-$\gamma$ | 0.004367 | 0.033220 |
| 12:10583611:C:T | IL-2 | 0.007334 | 0.009544 |
| 12:10583611:C:T | IL-6 | 0.009894 | 0.003338 |
| 19:16441973:G:A | TNF-$\alpha$ | 0.005125 | 0.019800 |
| 11:128070535:A:G | TNF-$\alpha$ | 0.000439 | 0.199147 |
| 11:128070535:A:G | IFN-$\gamma$ | 0.002636 | 0.026951 |

**Supplementary Table 6: Direct association tests between mean cytokine response expression per individual and genotype at the csaQTL lead SNP**. For each csaQTL and corresponding enriched gene set implicating a cytokine response as part of the csaQTL-associated phenotype, we followed up with a direct association test between genotype values for the csaQTL lead SNP and estimated cytokine response level per individual (**Methods**). We report the variance explained by the lead SNP in mean cytokine response expression per individual, as well as the result of a one-tailed t-test evaluating the significance of the association, after controlling for age, sex, and gPC1-6.

| Cell Type | Lead SNP | eGene | Beta | P | csaQTL | Colocalization |
|---|---|---|---|---|---|---|
| NK | 12:10594848:C:A | KLRC1 | -0.4 | 1.0e-41 | NK_12:10583611:C:T | 0.96 |
| NK | 12:10580062:C:T | KLRC2 | -0.86 | 5.2e-64 | NK_12:10583611:C:T | 2.9e-14 |
| NK | 12:10574001:T:C | KLRC3 | -0.59 | 1.1e-117 | NK_12:10583611:C:T | 6.6e-05 |
| B | 12:10561279:C:G | KLRK1 | -0.6 | 2.3e-56 | NK_12:10583611:C:T | 4.0e-05 |
| B | 15:80311721:T:C | BCL2A1 | 0.31 | 2.6e-64 | Myeloid_15:80263217:C:T | 6.5e-06 |

**Supplementary Table 7: eQTLs detected in the OneK1K dataset as candidates to colocalize with csaQTLs**. For each csaQTL, we tested the csaQTL lead SNP for eQTL associations to all *cis*-genes within a 2 megabase window within each single-cell object (T, B, NK, myeloid and all cells) using a model with pseudo-bulked gene expression per sample (**Methods**). For each triple (csaQTL lead SNP, eGene, expression cell type) with a $p<5x10^{-4}$ association in the pseudobulk eQTL model, we further tested the eQTL association using a single-cell-resolution model (**Methods**). For all triples (csaQTL lead SNP, eGene, expression cell type) with $p<1x10^{-6}$ in the single-cell eQTL model, we applied the single-cell eQTL model to all SNPs in a 2 megabase window centered on the csaQTL lead SNP to identify an eQTL lead SNP and we estimated eQTL colocalization with the csaQTL (**Methods**). In this table, we display one row for each triple (csaQTL lead SNP, eGene, expression cell type) that met these criteria for colocalization analysis. Specifically, we report the **csaQTL**, the **Cell Type** in which expression was modeled for the eQTL, the eQTL **Lead SNP**, the eQTL **Beta**, the eQTL **p**-value, and the posterior probability of a shared causal variant between the csaQTL and eQTL (**Colocalization**).

| Cell Type | Lead SNP | eGene | Beta | P | csaQTL | Coloc. |
|---|---|---|---|---|---|---|
| Whole blood | 15:80260274:A:T | BCL2A1 | -0.24 | 9.3e-52 | Myeloid_15:80263217:C:T | 0.98 |
| NK | 11:128085408:C:T | ETS1 | 0.39 | 4.3e-04 | NK_11:128070535:A:G | 0.40 |
| Stim. CD8+ T | 12:10591281:G:A | KLRC4 | 0.89 | 4.9e-06 | NK_12:10583611:C:T | 0.78 |

**Supplementary Table 8: Published eQTLs that colocalize with the GeNA csaQTLs**. We reviewed eQTL summary statistics from five published studies (**Methods**) to identify eQTLs that colocalize with the csaQTLs. We display one row for each triple (csaQTL lead SNP, eGene, expression cell type) with a published eQTL that colocalizes with a GeNA csaQTL. Specifically, we report the **csaQTL**, the **Cell Type** in which expression was modeled for the eQTL, the eQTL **Lead SNP**, the eQTL **Beta**, the eQTL **p**-value, and the posterior probability of a shared causal variant between the csaQTL and eQTL (**Coloc.**).

| Cell Type | Lead SNP | Associated Condition | Subcohort | N, Discovery | N, Subcohort | P, Subcohort | r-sq., Traits |
|---|---|---|---|---|---|---|---|
| NK | 19:16441973:G:A | Asthma | Known absence of asthma | 935 | 444 | 2.54e-6 | 0.72 |
| NK | 19:16441973:G:A | Type 1 Diabetes | Known absence of autoimmune disease | 935 | 454 | 6.08e-4 | 0.75 |
| NK | 12:10583611:C:T | Psoriasis | Known absence of autoimmune disease | 935 | 454 | 2.99e-4 | 0.72 |
| NK | 2:111851212:C:T | Ovarian Cancer | Known absence of ovarian cancer | 935 | 513 | 5.59e-3 | 0.72 |
| Myeloid | 15:80263217:C:T | Primary Sclerosing Cholangitis | Known absence of autoimmune disease | 523 | 247 | 7.48e-5 | 0.75 |

**Supplementary Table 9: Testing for csaQTL persistence in custom OneK1K subcohorts that exclude clinical conditions of interest**. For each csaQTL that corresponds to a disease risk locus, we re-tested the csaQTL using GeNA in a custom subcohort of the OneK1K dataset that excluded all individuals lacking clinical metadata and excluding individuals with specific diagnoses. For each csaQTL tested in a custom subcohort (one per row) we display the **Lead SNP** and source csaQTL GWAS (**Cell Type**), the clinical condition associated with genetic risk at the same locus (**Associated Condition**), the inclusion criteria for the custom OneK1K subcohort (**Subcohort**), and the count of individuals in the original GWAS (**N, Discovery**) and subcohort (**N, Subcohort**). We found that all csaQTLs persisted in their custom cohorts, with GeNA $p<6\text{x}10^{-3}$ (**P, Subcohort**) and with consistency between the csaQTL-associated traits detected by GeNA in the discovery and custom subcohort analyses (Pearson's r-squared between sample-level phenotype values in each cohort, **r-sq., Traits**).

| Celltype | CHR:POS | Cis Vargenes | Expr Cor (r) | %tile Expr Cor ($r^2$) | $r^2$ Pheno, Nbhds | $r^2$ Pheno, Samples | GWAS P | P, Masked |
|---|---|---|---|---|---|---|---|---|
| | | BCL2A1 | -0.358 | 97 | | | | |
| | | RP11-81A1.6 | 0.021 | 41 | | | | |
| Myeloid | 15:80263217 | ST20 | -0.016 | 32 | 1.00 | 1.00 | 2.61e-08 | 2.71e-08 |
| NK | 2:111851212 | MIR4435-2HG | 0.025 | 80 | 1.00 | 1.00 | 1.76e-09 | 3.94e-09 |
| NK | 11:128070535 | ETS1 | 0.011 | 55 | 1.00 | 1.00 | 2.48e-13 | 1.65e-13 |
| | | CD69 | 0.164 | 99 | | | | |
| | | KLRC2 | 0.127 | 98 | | | | |
| | | KLRK1 | 0.081 | 95 | | | | |
| | | KLRC3 | 0.082 | 95 | | | | |
| | | LINC02446 | 0.05 | 90 | | | | |
| | | RP11-726G1.2 | 0.03 | 82 | | | | |
| | | CLEC2D | 0.021 | 75 | | | | |
| | | KLRC4 | 0.018 | 70 | | | | |
| | | KLRC1 | -0.014 | 63 | | | | |
| | | CLECL1 | 0.006 | 36 | | | | |
| NK | 12:10583611 | KLRK1-AS1 | 0.006 | 35 | 0.95 | 0.91 | 1.96e-11 | 3.9e-10 |
| | | HSH2D | 0.091 | 95 | | | | |
| | | KLF2 | -0.016 | 68 | | | | |
| | | MYO9B | -0.011 | 59 | | | | |
| | | TPM4 | 0.008 | 50 | | | | |
| | | CTC-429P9.3 | -0.008 | 49 | | | | |
| | | CTC-429P9.5 | -0.003 | 20 | | | | |
| NK | 19:16441973 | BRD4 | -0.002 | 12 | 1.00 | 0.99 | 1.96e-13 | 1.4e-14 |

**Supplementary Table 10: csaQTL association tests for each lead SNP from the discovery GWAS, after masking expression of *cis*-genes**. **Celltype** of the discovery GWAS and **CHR:POS** for each lead SNP are shown, along with the variable genes from the discovery GWAS within a 2 megabase window centered on the SNP (***Cis*-Vargenes**), which were removed from the dataset for the masked test. For each *cis*-variable gene, we also show for the csaQTL in the discovery GWAS the correlation (Pearson's r) between expression of that gene and csaQTL phenotype value per neighborhood (**Expr Cor**). To contextualize these raw correlations among the strengths of all variable genes' expression correlations to the csaQTL per-neighborhood phenotype values, we also show the percentile among all variable genes' correlations (Pearson's $r^2$; **%ile Expr Cor**). To compare csaQTLs in the discovery versus masked versions of the dataset, the correlations are shown between the per-sample (**$r^2$ Pheno, Samples**) csaQTL phenotype values. We also compare the per-gene correlations to the neighborhood-level phenotype in the discovery and cis-masked analyes (**$r^2$ Pheno, Nbhds**). Finally, we compare csaQTL association strength in the discovery (**GWAS P**) and masked (**P, Masked**) datasets.

| Cell Type | Disease PRS | Cohort | N | P | P, adjusted |
|---|---|---|---|---|---|
| Myeloid | SLE | Known absence of SLE | 282 | 0.003996 | 0.03996 |
| B | SLE | Known absence of SLE | 492 | 0.318681 | 0.455259 |
| NK | SLE | Known absence of SLE | 511 | 0.249750 | 0.41625 |
| T | SLE | Known absence of SLE | 531 | 0.369630 | 0.462038 |
| allcells | SLE | Known absence of SLE | 532 | 0.053946 | 0.17982 |
| Myeloid | RA | Known absence of RA | 274 | 0.165834 | 0.331668 |
| B | RA | Known absence of RA | 478 | 0.139860 | 0.331668 |
| NK | RA | Known absence of RA | 498 | 0.527473 | 0.586081 |
| T | RA | Known absence of RA | 517 | 0.605395 | 0.605395 |
| allcells | RA | Known absence of RA | 518 | 0.026973 | 0.134865 |
| Myeloid | SLE | Known absence of any autoimmune disease | 270 | 0.018981 | N.A. |
| allcells | RA | Known absence of any autoimmune disease | 507 | 0.011988 | N.A. |

**Supplementary Table 11: Testing associations to disease PRS in the OneK1K cohort**. Using published polygenic risk scores, we quantified genetic risk for systemic lupus erythematosus (SLE) and rheumatoid arthritis (RA) for each individual in the OneK1K cohort. We tested associations to each PRS within all cells and each major cell type using Covarying Neighborhood Analysis (CNA)[3]. For each association test we report the **Disease PRS** tested in the **Cell Type** and the resulting **P** value from CNA's global association test. We excluded individuals lacking clinical metadata or with a diagnosis of the disease of interest. To account for multiple hypothesis testing, we used the Benjamini-Hochberg FDR correction approach and report the **P, adjusted**. For each association test that passed a nominal p<0.05 threshold, we tested again in a subcohort including only individuals with a known absence of any autoimmune disease.

| Cell Type | N cases | N conrtrols | P |
|---|---|---|---|
| Myeloid | 9 | 9 | <1e-4 |
| allcells | 16 | 16 | <1e-4 |
| NK | 15 | 15 | <1e-4 |
| B | 15 | 15 | <1e-4 |
| T | 16 | 16 | <1e-4 |

**Supplementary Table 12: Case-control association testing for rheumatoid arthritis (RA) in the OneK1K dataset**. For each major cell type, we created a single-cell object containing profiling from all individuals with an RA diagnosis that passed quality control (e.g., at least 25 cells of that type available) and profiling for individuals with a known absence of RA downsampled at random to an equal number as cases. We performed case-control association tests using CNA[3]. All tests attained the minimum possible permutation-based p-value, indicating $p<1x10^{-4}$.

### Supplementary Figures

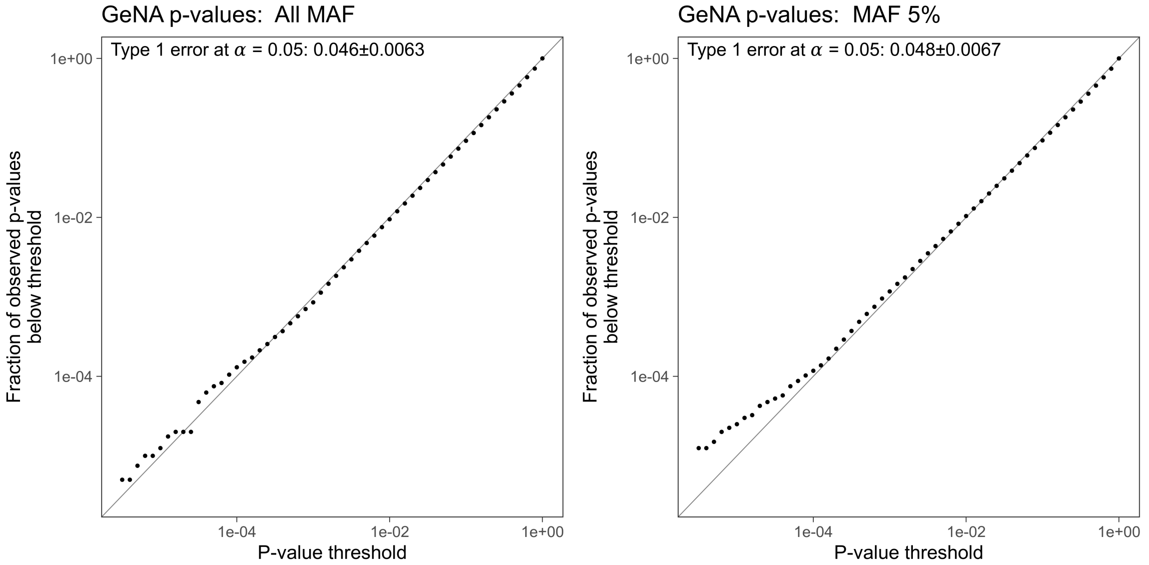


**Supplementary Figure 1: Performance of GeNA on simulations with no true associations between genotypes and cell states**. We plot the cumulative fraction of GeNA p-values (y-axis) that fall below a given p-value threshold (x-axis) for simulated genotypes across our full included MAF spectrum 0.5>=MAF>0.05 (**left**) and for simulated genotypes with 0.055>=MAF>0.05 (**right**). Both x and y axes are log-scaled.

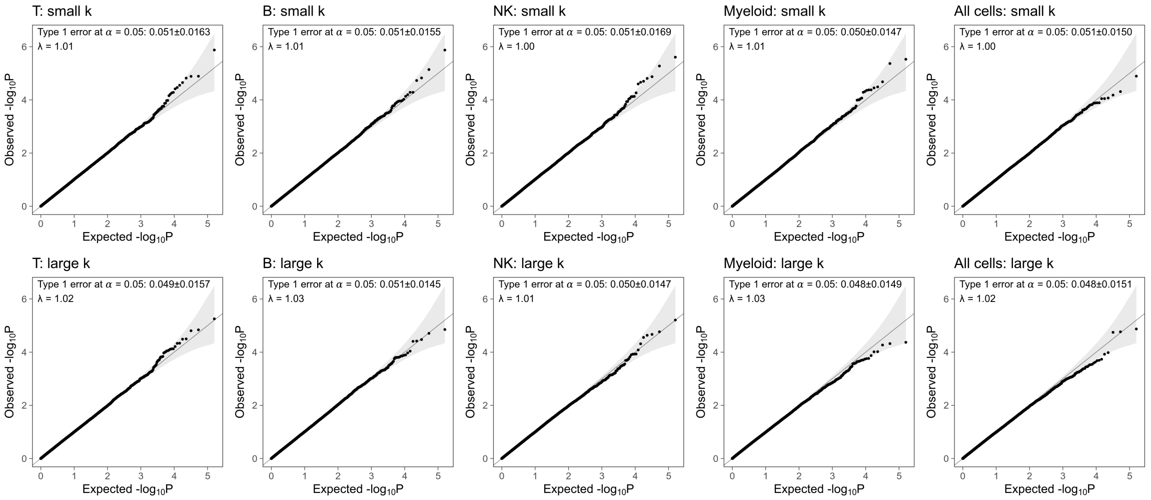


**Supplementary Figure 2: Quantile-quantile plots of null simulations for SNPs sampled from all included MAFs**. We plot GeNA results from simulated genotypes with no true associations to any cell state. We plot results separately by major cell type (T, B, NK, myeloid and all cells) and by value of $k$. GeNA considers two values of $k$ by default for a given dataset. We plot performance separately for each (cell type, $k$) pair for the values of $k$ used by default by GeNA in each major cell type (**Supplementary Figure 4**). We also display the rate of false positive associations and lambda value for each QQ plot.

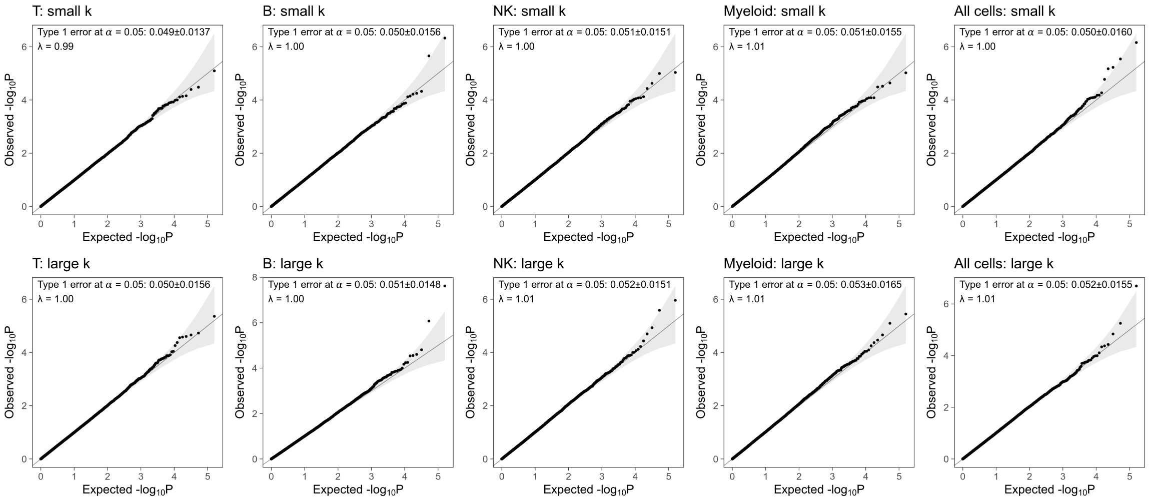


**Supplementary Figure 3: Quantile-quantile plots of null simulations for SNPs with MAF 0.05-0.055**. We plot GeNA results from simulated genotypes with no true associations to any cell state. We plot results separately by major cell type (T, B, NK, myeloid and all cells) and by value of $k$. GeNA considers two values of $k$ by default for a given dataset. We plot performance separately for each (cell type, $k$) pair for the values of $k$ used by default by GeNA in each major cell type (**Supplementary Figure 4**). We also display the rate of false positive associations and lambda value for each QQ plot.

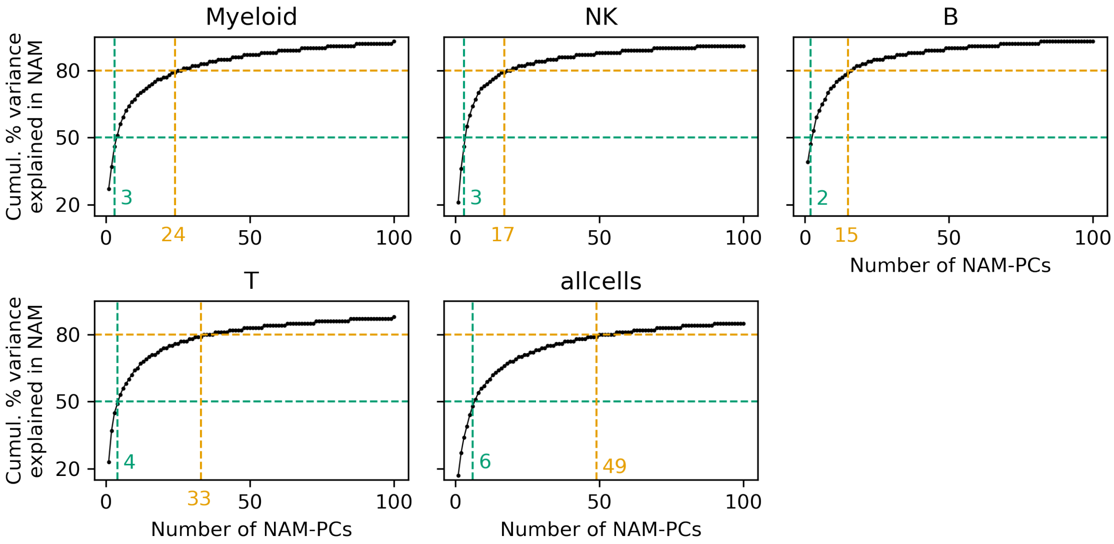


**Supplementary Figure 4: Values of $k$ considered by GeNA by default for each major cell type in the OneK1K dataset**. For each major cell type, two values of $k$, the number of NAM-PCs to include in the csaQTL model, are selected by GeNA using 50% (green) and 80% (orange) thresholds of percent variance explained.

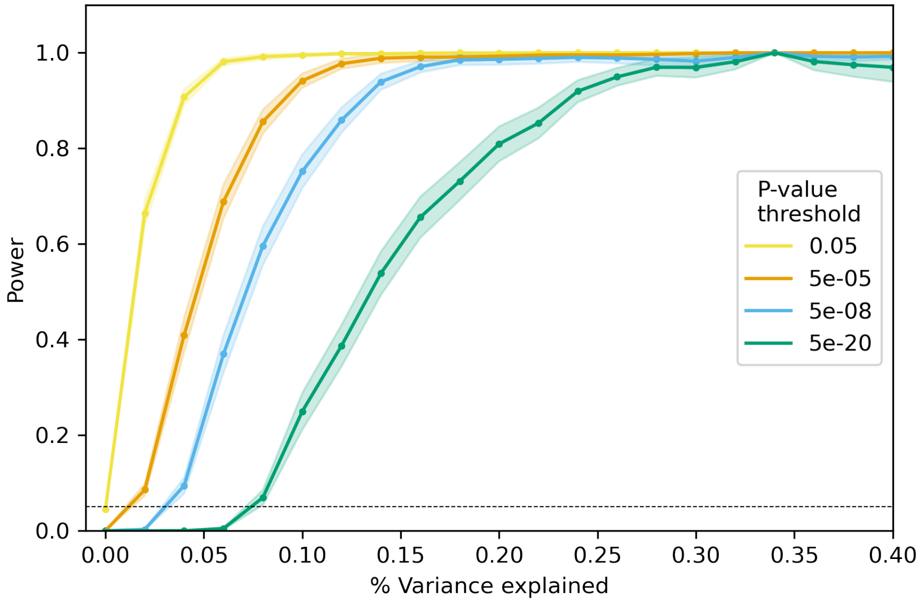


**Supplementary Figure 5: Power to detect associations between simulated genotype values and real single-cell traits**. For 94 real cell state abundance traits that vary across individuals in the OneK1K dataset, we defined simulated genotype values per individual to create true associations to these traits. By including random noise in the simulated genotypes we can use these data to quantify the fraction of true associations detected by GeNA (power) across a spectrum of noise levels. At each noise level, we show the mean and standard error of statistical power across traits for a given p-value threshold. A dashed line is shown at y=0.05.

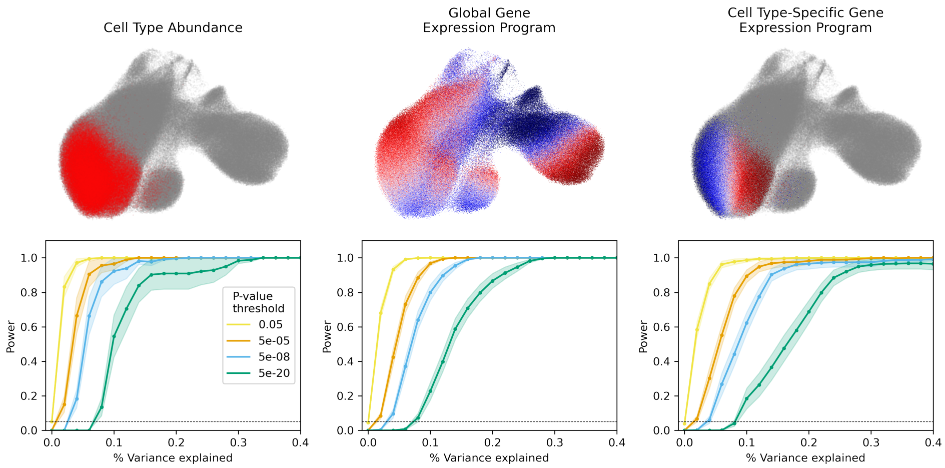


**Supplementary Figure 6: Power to detect associations between simulated genotype values and real cell state abundance traits, split by phenotype category**. We defined real cell state abundance traits within three categories (**left, middle, right**): 1) differential abundance of a cluster-based cell type, 2) increased expression of a gene set across all cells, and 3) increased expression of a gene set within a cluster-based cell type. In this figure, we split our simulation results by phenotype category. At each noise level, we show the mean and standard error of statistical power across the traits in that category for a given p-value threshold. A dashed line is shown at y=0.05. Above each power plot, we include an illustration of one example phenotype from that category. Each neighborhood is colored according to its abundance correlation to the simulated genotype, with larger positive correlations in darker red, larger negative correlations in darker blue, and correlations equal to zero in grey.

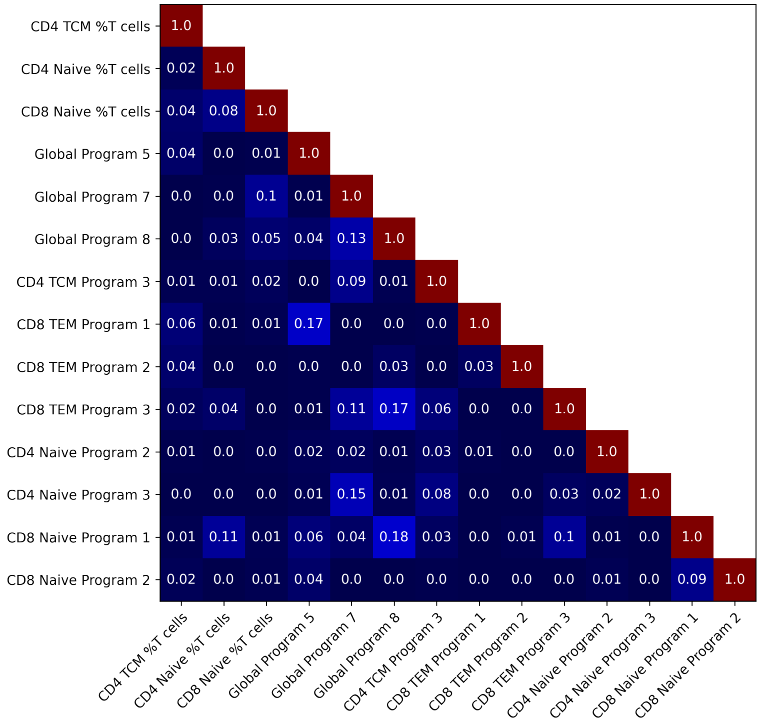


**Supplementary Figure 7: Heatmap of correlations among per-sample values for 14 cell state abundance traits used in our non-null simulated GWAS for T cells**. For each of these 14 traits with pairwise Pearson's $r^2 < 0.2$, we display the $r^2$ correlation in per-sample values between each pair of traits.

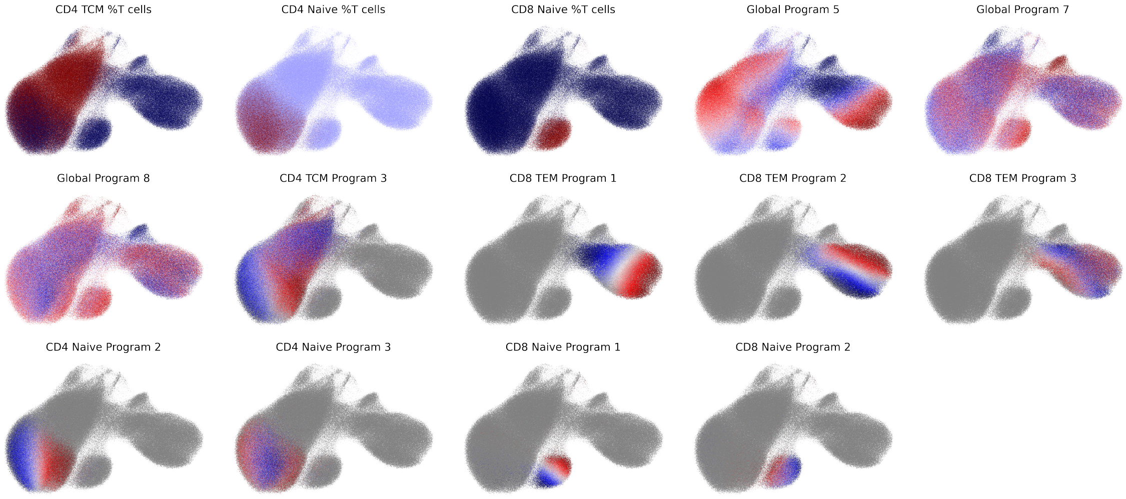


**Supplementary Figure 8: Illustration of 14 real cell state abundance traits used in our non-null simulated GWAS for T cells**. For each trait in **Supp. Fig. 7**, we plot the true cell-level phenotype for which we simulated associated genotypes with varying levels of noise. Each UMAP includes one dot per T cell in the OneK1K dataset. Cells that do not affect the trait are colored grey. For example, for the "CD8 Naïve Program 1" trait, we used a gene expression program that varies substantially across naïve CD8+ T cells. We quantified the usage of that program across all naïve CD8+ T cells and defined the trait value per individual in the dataset as the mean use of that gene expression program across all naïve CD8+ T cells in that individual's sample. Therefore, for the "CD8 Naïve Program 1" trait we color all cells that are not naïve CD8+ T cells grey because they do not influence the trait and we color each naïve CD8+ T cell by its use of the gene expression program. Cells with greater use of the program are colored deeper red and cells with less use of the program are colored deeper blue.

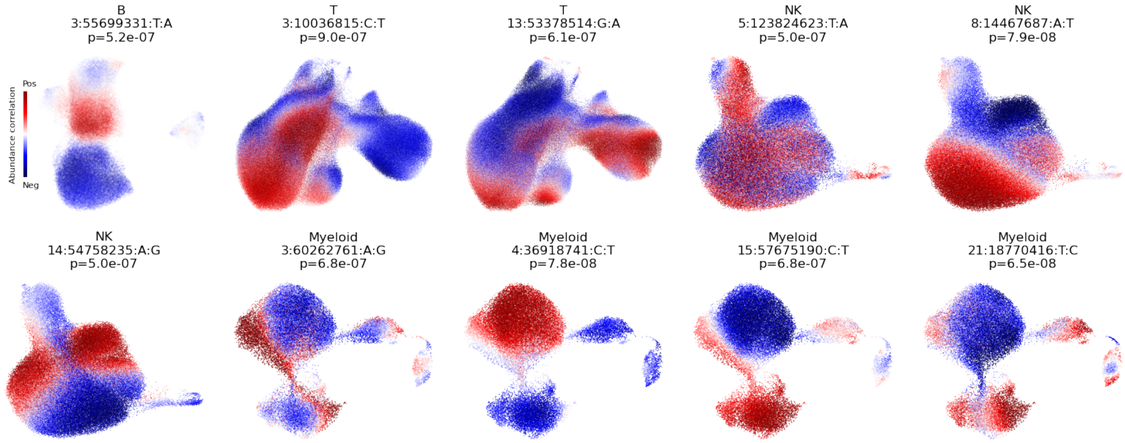


**Supplementary Figure 9: Traits associated with lead SNPs for each suggestive association in our csaQTL GWAS**. We display results for loci that passed a p<1x$10^{-6}$ threshold for suggestive association in our csaQTL GWAS but did not attain genome-wide significance (p>5x$10^{-8}$). For each lead SNP ("CHR:POS:REF:ALT"), we plot a UMAP of the included cells colored by neighborhood-level phenotype and we report the associated p-value and source GWAS (T, B, NK or myeloid cells). For each source GWAS with multiple suggestive loci, we observe associations to distinct traits detected by GeNA in a single genome-wide survey.

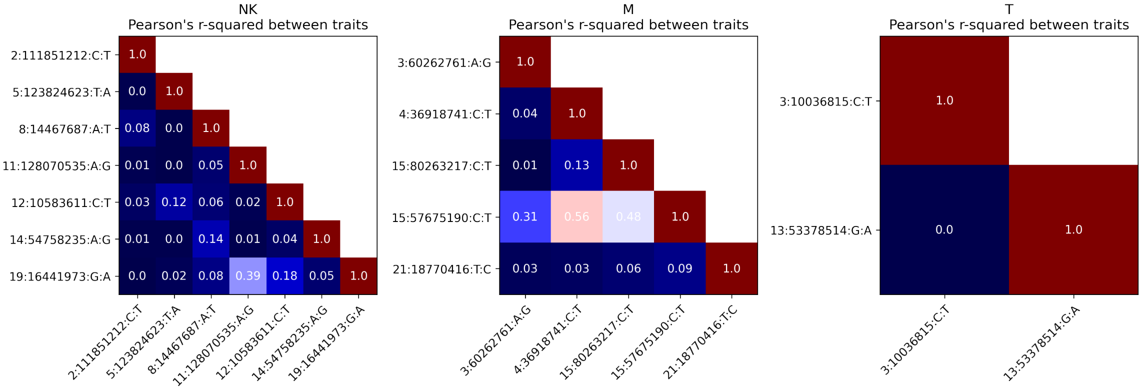


**Supplementary Figure 10: Correlations among csaQTL-associated traits**. For each of our csaQTL GWAS with more than one association that passed a p<1x$10^{-6}$ threshold for suggestive association, we plot pairwise Pearson's r-squared values among the traits associated with the lead SNPs for each suggestive or genome-wide significant locus. Specifically, for each pair of loci, we compute the correlation between the sample-level phenotypes corresponding to each lead SNP. While some phenotypes have shared features, such as the loci on chromosomes 12, 11, and 19 from the NK GWAS, these results highlight the diversity of phenotypes to which csaQTL associations can be detected in a single genome-wide survey using GeNA.

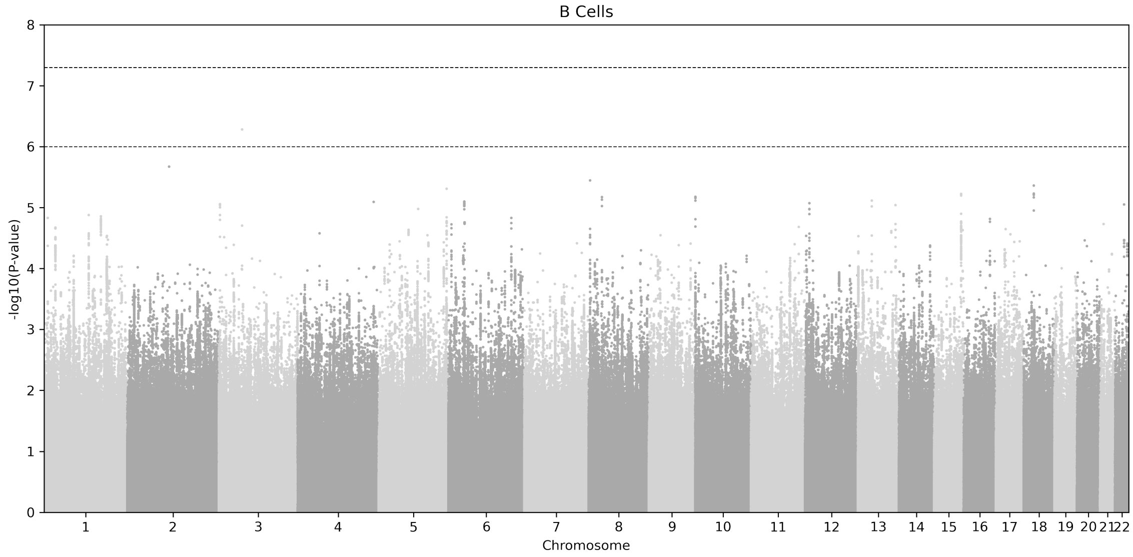


**Supplementary Figure 11: GWAS for csaQTLs within B cells**. Dashed horizontal lines indicate p=5x$10^{-8}$ and p=1x$10^{-6}$ thresholds.

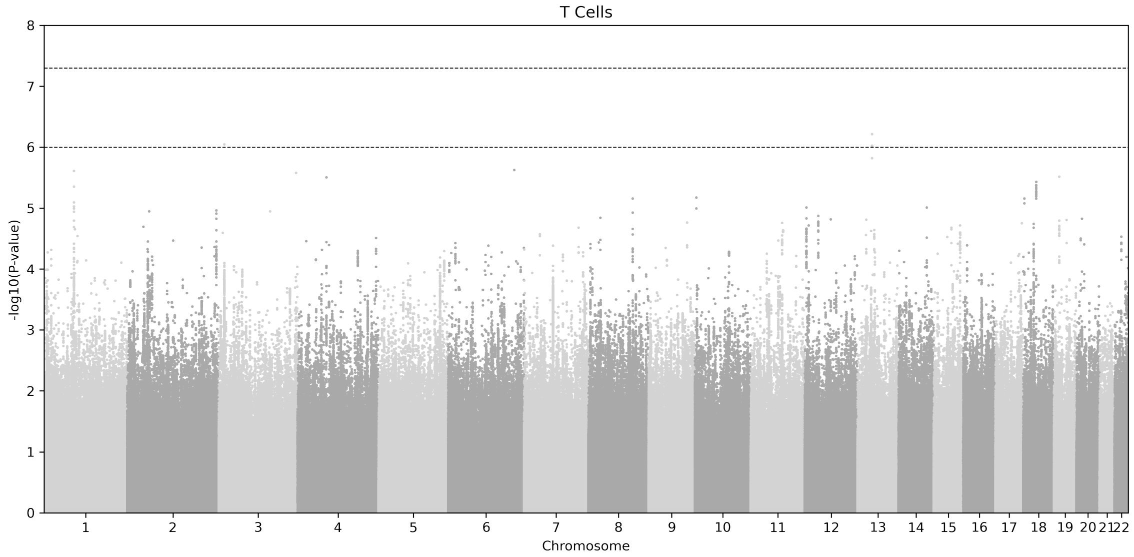


**Supplementary Figure 12: GWAS for csaQTLs within T cells**. Dashed horizontal lines indicate p=5x10$^{-8}$ and p=1x10$^{-6}$ thresholds.

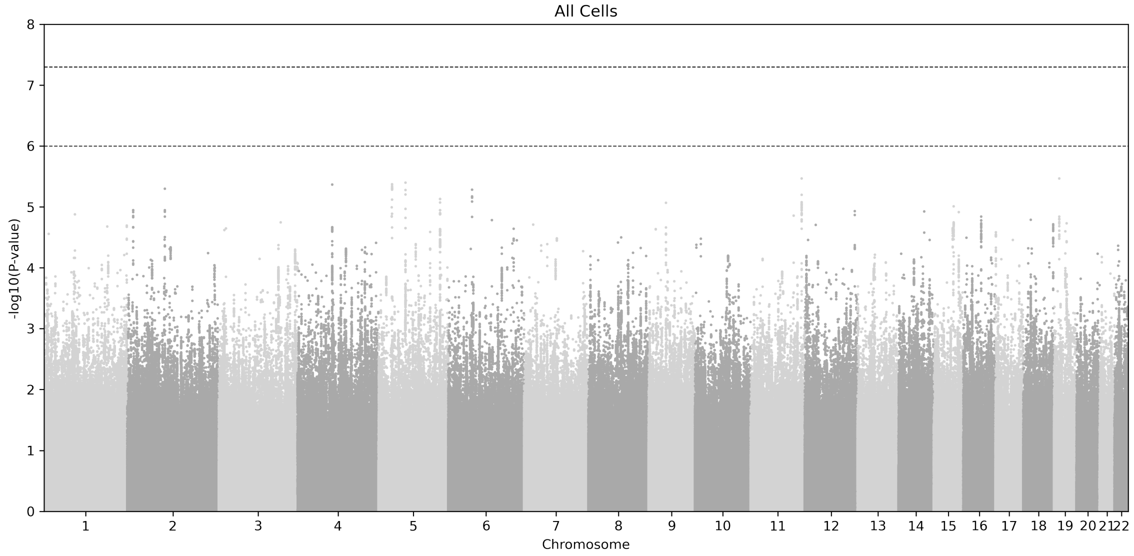


**Supplementary Figure 13: GWAS for csaQTLs across all cells in the dataset**. Dashed horizontal lines indicate p=5x10$^{-8}$ and p=1x10$^{-6}$ thresholds.

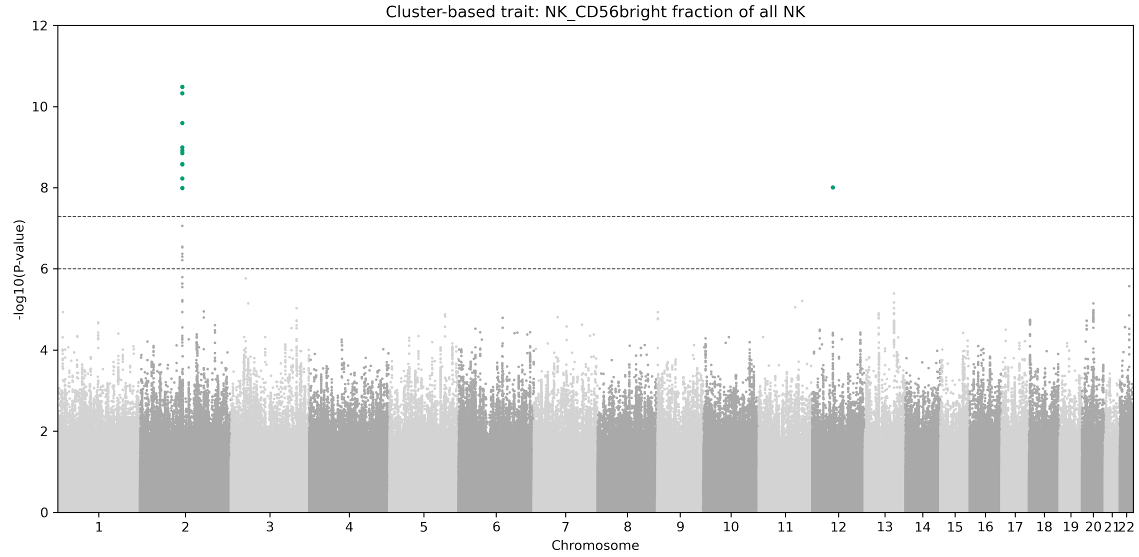


**Supplementary Figure 14: GWAS for CD56$^{br}$ fractional abundance out of all NK cells, as defined by published cell assignments to clusters in this dataset**. Dashed horizontal lines indicate p=5x10$^{-8}$ (genome-wide significant) and p=1x10$^{-6}$ thresholds (**Methods**).

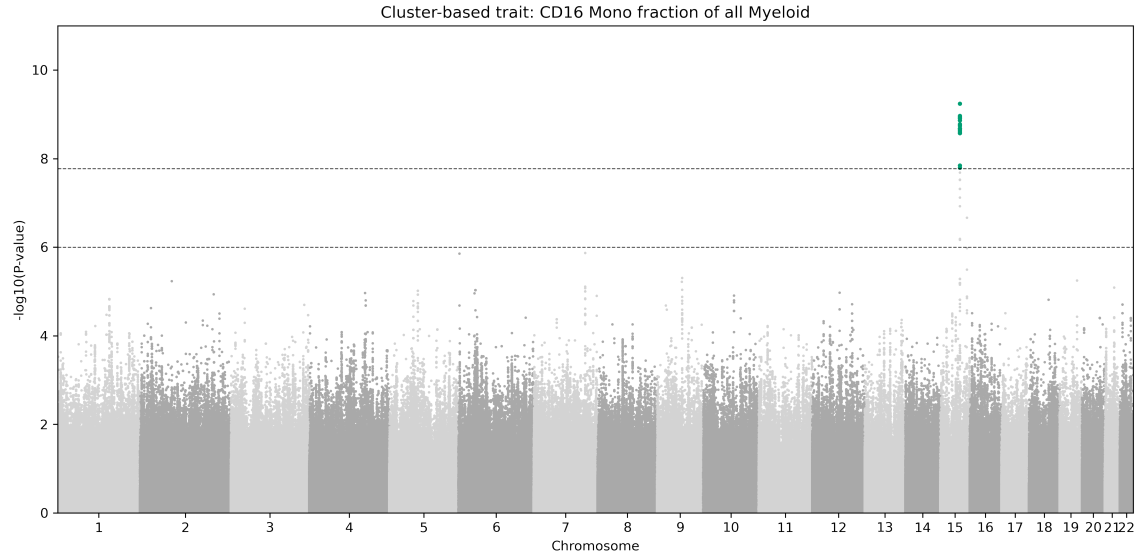


**Supplementary Figure 15: GWAS for CD16+ monocyte fractional abundance out of all myeloid cells, as defined by published cell assignments to clusters in this dataset**. Dashed horizontal lines indicate p=1.67x10$^{-8}$ (genome-wide significant) and p=1x10$^{-6}$ thresholds (**Methods**).

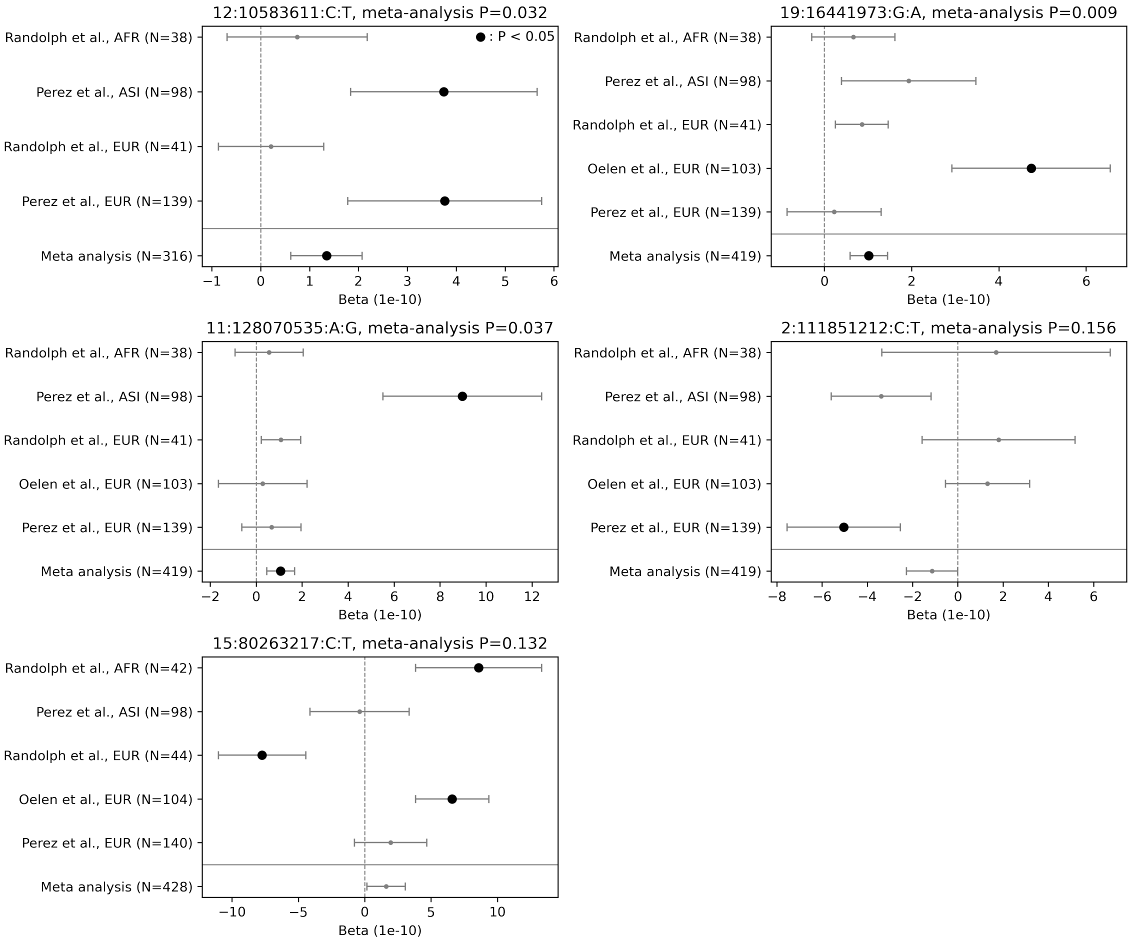


**Supplementary Figure 16: Replication testing within five independent cohorts for each csaQTL identified by GeNA in the OneK1K dataset.** Each plot corresponds to one csaQTL and includes all cohorts in which genotypes for the lead SNP were available. For each replication cohort and SNP pair, we show the replication beta (dot) and one standard error (whiskers). We also show the meta analysis test result associated with each csaQTL across all five replication cohorts. All association tests with $p<0.05$ are shown with a bold marker.

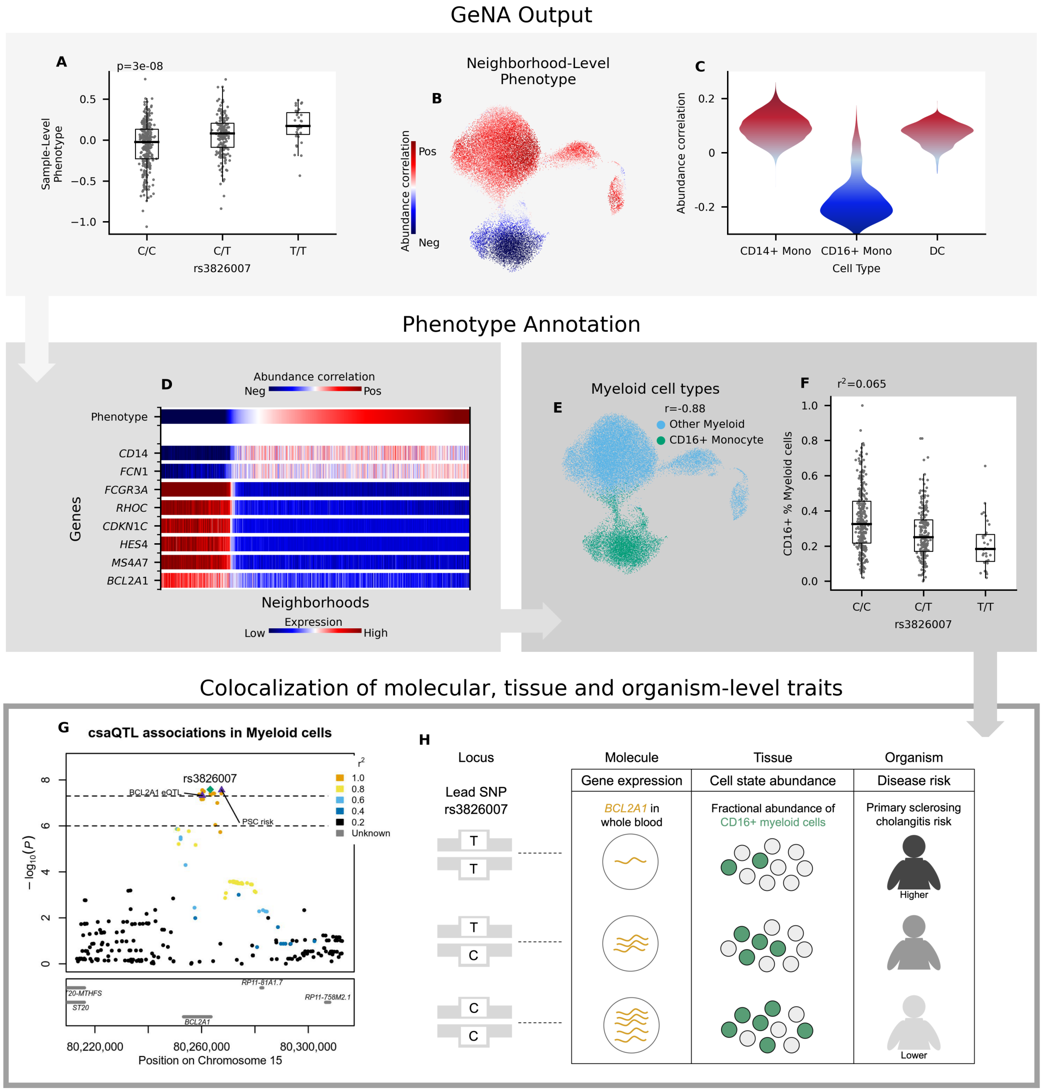


**Supplementary Figure 17: Characterization of the csaQTL at 15q25.1**. (**A**) Boxplot of sample-level phenotype values for each individual, organized by genotype at the lead SNP. We also show the GeNA p-value. (**B**) UMAP of Myeloid cells colored by neighborhood-level phenotype value (i.e., correlation between cell abundance and dose of alternative allele per neighborhood). (**C**) Violinplot of neighborhood-level phenotype value distribution within CD14+ monocytes, CD16+ monocytes and dendritic cells. (**D**) Heatmap of expression across neighborhoods for genes with strong correlations in expression to the csaQTL neighborhood-level phenotype. Neighborhoods are arrayed along the x-axis by phenotype value. (**E**) UMAP of myeloid cells colored by cell type assignment to the CD16+ monocyte cluster. We also show the Pearson's r value between neighborhood-level phenotype values and a binary encoding of CD16+ monocyte cluster membership per cell. (**F**) Boxplot of cluster-based CD16+ monocyte % myeloid cells trait value per donor by genotype. The csaQTL lead SNP explains 6.5% of variance in this phenotype. (**G**) Locus zoom plot with one marker per tested SNP, genomic position along the x-axis, and GeNA p-value on the y-axis. Each SNP marker is colored by LD value relative to the lead SNP. The lead SNP is labeled with a green diamond. The *BCL2A1* eQTL lead SNP and primary sclerosing cholangitis risk lead SNP are labeled with purple triangles. (**H**) Diagram of genotypes for the csaQTL lead SNP and colocalizing associations to molecular, tissue and organism-level traits at this locus.

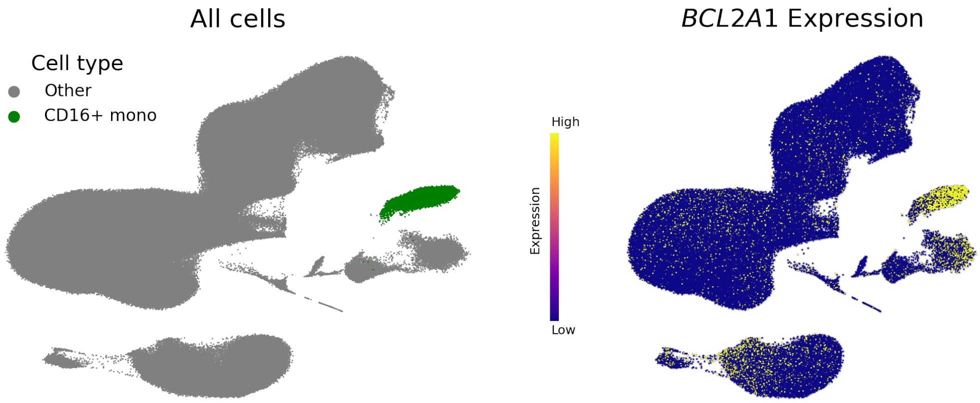


**Supplementary Figure 18: Among peripheral blood mononuclear cells, CD16+ monocytes preferentially express *BCL2A1*.** (**Left**) UMAP of all cells in the OneK1K dataset with cells in the CD16+ monocyte cluster colored green and all other cells colored grey. (**Right**) UMAP of all cells in the OneK1K dataset colored according to (normalized and scaled) expression of *BCL2A1*.

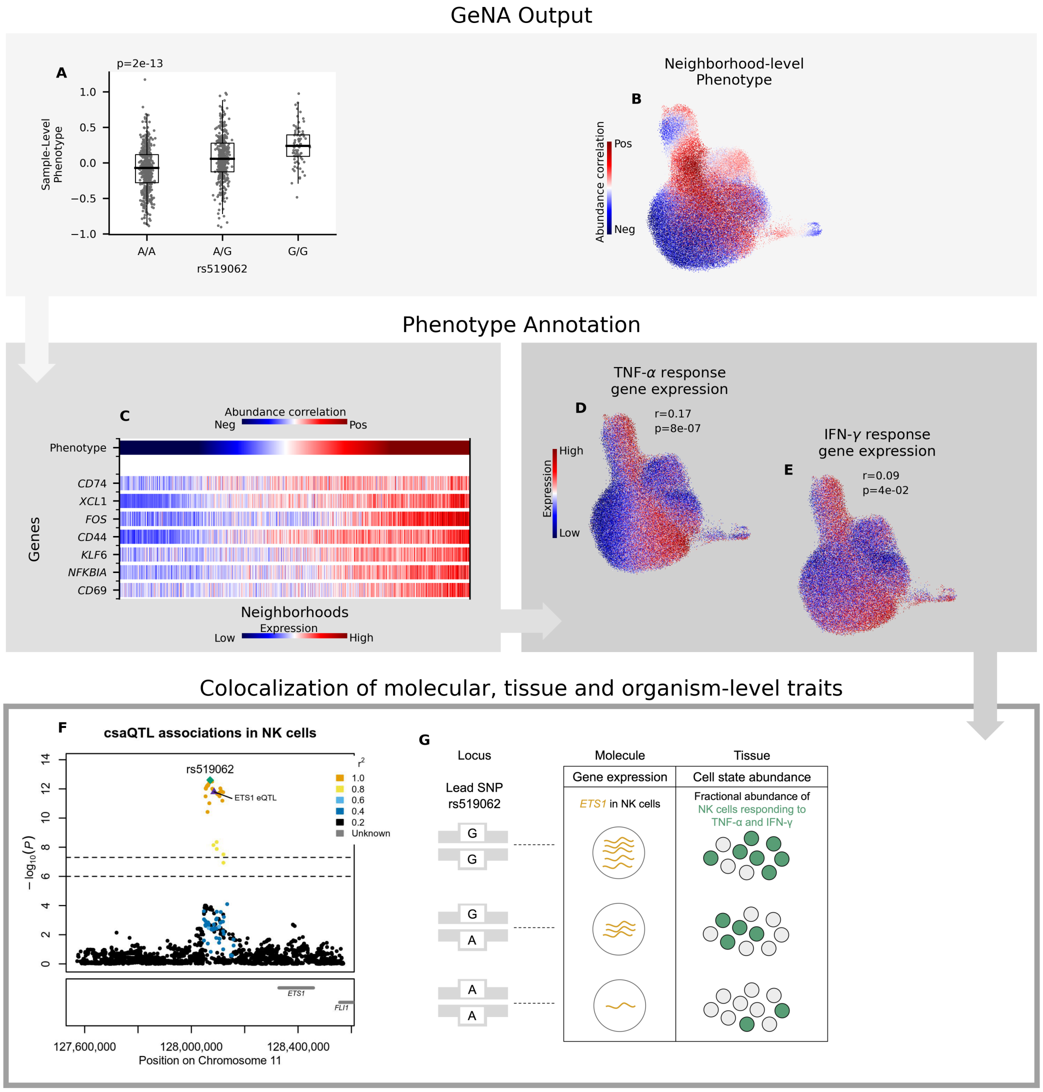


**Supplementary Figure 19: Characterization of the csaQTL at 11q24.3**. At 11q24.3, lead SNP rs519062-G associates with expansion of activated NK cell states expressing TNF-$\alpha$ and IFN-$\gamma$ response genes. Transcription factor *ETS1* is encoded by the nearest gene to all SNPs in the locus. Schmiedel *et al.*[4] report a suggestive eQTL in NK cells (lead SNP p=4x10$^{-4}$; Pr$_{coloc}$=40%) for which rs519062-G associates with increased *ETS1*. Interestingly, primary human NK cells with experimental *ETS1* knockout exhibit decreased IFN-$\gamma$ production in response to stimulation[5]. (**A**) Boxplot of sample-level phenotype values for each individual, organized by lead SNP genotype, with GeNA p-value. (**B**) UMAP of NK cells colored by neighborhood-level phenotype values. (**C**) Heatmap of expression across neighborhoods for genes with strong expression correlations to the neighborhood-level phenotype. Neighborhoods are ordered along the x-axis by phenotype value. The phenotype-correlated genes include general markers of NK activation (*CD69*, *NFKBIA*) as well as TNF-$\alpha$ (*FOS*, *CD44*, *KLF6*) and IFN-$\gamma$ (*CD74*, *XCL1*) response. (**D-E**) Gene set enrichment analysis identified significant activation of TNF-$\alpha$ and IFN-$\gamma$ response in association with the csaQTL phenotype. We show UMAPs of NK cells colored by summed expression of (D) TNF-$\alpha$ response genes and (E) IFN-$\gamma$ response genes. We report the Pearson's r across neighborhoods between phenotype values and summed expression for the gene set, with FDR-adjusted enrichment p-value. (**F**) Locus zoom plot with one marker per tested SNP, genomic position along the x-axis, and GeNA p-value on the y-axis. Each marker is colored by LD value relative to the lead SNP. The csaQTL lead SNP is a green diamond. The *ETS1* eQTL lead SNP is a purple triangle. (**G**) Diagram of genotypes for the csaQTL lead SNP and colocalizing associations to molecular and tissue traits.

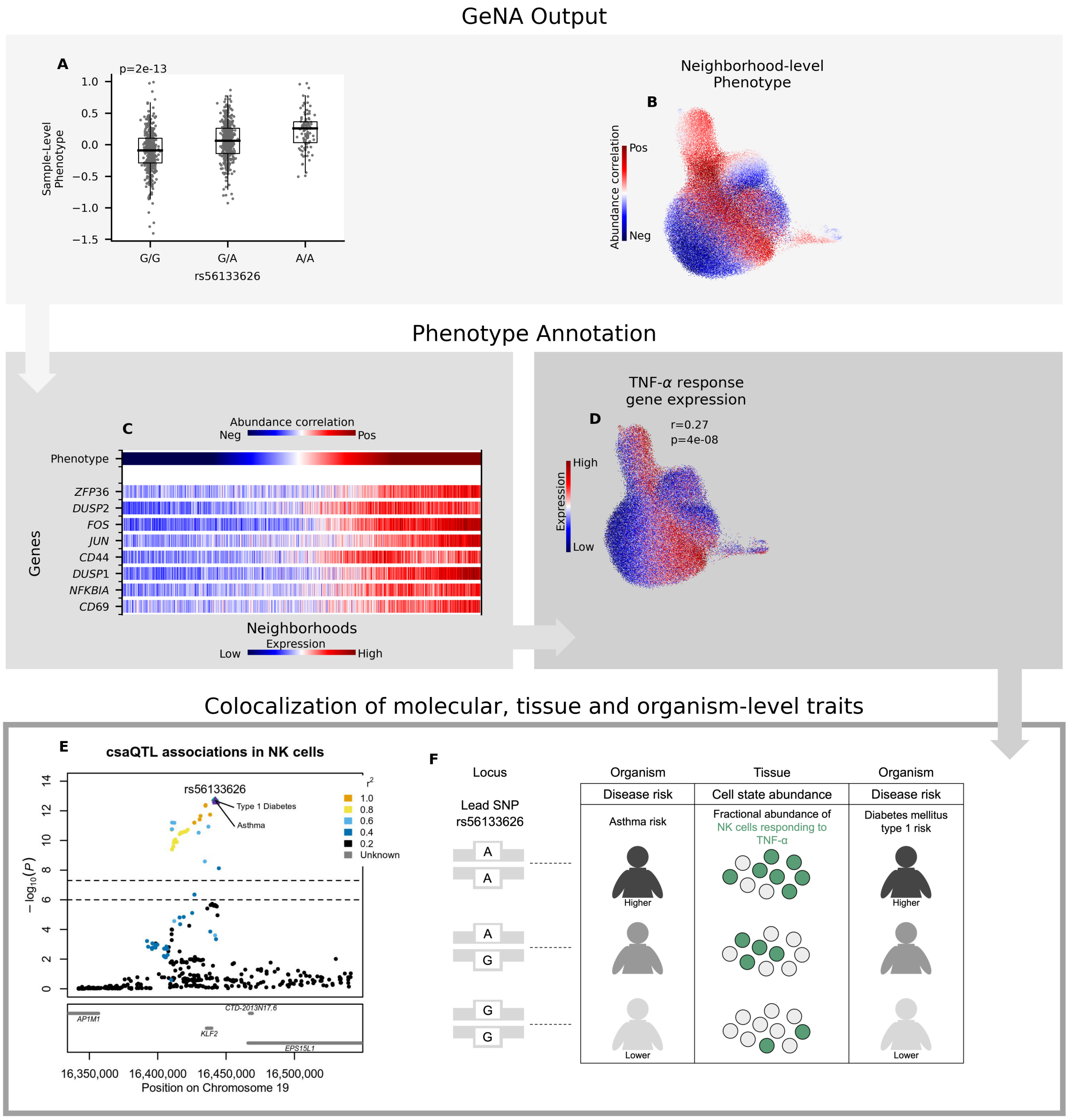


**Supplementary Figure 20: Characterization of the csaQTL at 19p13.11**. At 19p13.11, lead SNP rs55908509-A associates with expansion of activated NK cells expressing TNF-$\alpha$ response genes. All SNPs in the locus lie near the *KLF2* gene, but no colocalizing *KLF2* eQTLs were identified. (**A**) Boxplot of sample-level phenotype values for each individual, organized by lead SNP genotype, with GeNA p-value. (**B**) UMAP of NK cells colored by neighborhood-level phenotype values. (**C**) Heatmap of expression across neighborhoods for genes with strong expression correlations to the neighborhood-level phenotype. Neighborhoods are ordered along the x-axis by phenotype value. The phenotype-correlated genes include general markers of NK activation (*CD69*, *NFKBIA*) as well as TNF-$\alpha$ (*DUSP1*, *CD44*, *JUN*, *FOS*, *DUSP2*, *ZFP36*) response. (**D**) Gene set enrichment analysis identified significant activation of TNF-$\alpha$ response in association with the csaQTL phenotype. We show a UMAP of NK cells colored by summed expression of TNF-$\alpha$ response genes. We report the Pearson's r across neighborhoods between phenotype values and summed expression for the gene set, with FDR-adjusted enrichment p-value. (**E**) Locus zoom plot with one marker per tested SNP, genomic position along the x-axis, and GeNA p-value on the y-axis. Each marker is colored by LD value relative to the lead SNP. The csaQTL lead SNP is a green diamond. The lead SNPs for type 1 diabetes and asthma risk are annotated with purple triangles. (**F**) Diagram of genotypes for the csaQTL lead SNP and colocalizing associations to tissue and organism-level traits.

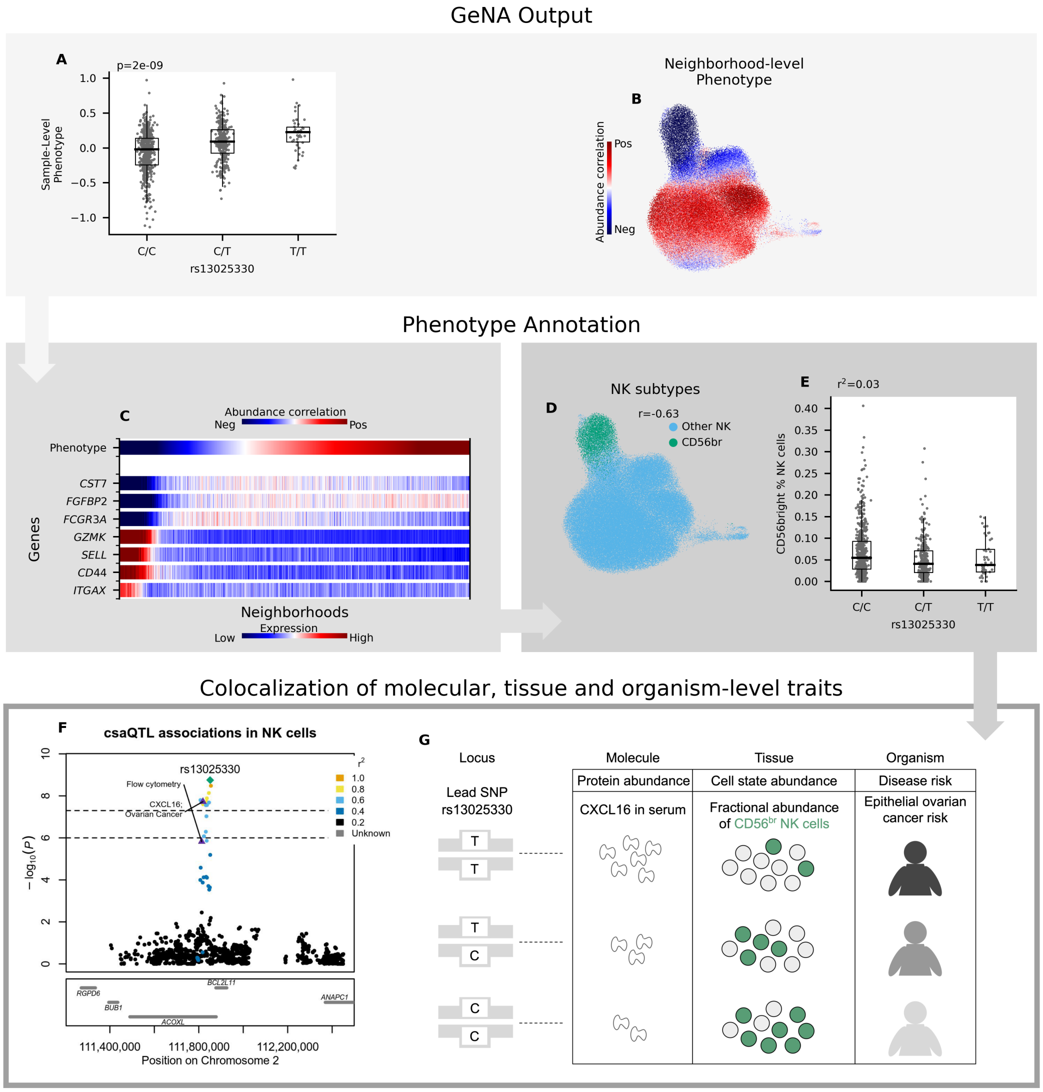


**Supplementary Figure 21: Characterization of the csaQTL at 2q13**. At 2q13, CD56$^{br}$ fraction of NK cells decreases in association with increasing dose of lead SNP rs13025330-T. The csaQTL colocalizes with a published pQTL for the abundance of chemokine CXCL16 in peripheral blood (pQTL lead SNP p=4.6x10$^{-18}$, Pr$_{coloc}$=84%); rs13025330-T associates with increased CXCL16 [6]. (**A**) Boxplot of sample-level phenotype values for each individual, organized by lead SNP genotype, with GeNA p-value. (**B**) UMAP of NK cells colored by neighborhood-level phenotype values. (**C**) Heatmap of expression across neighborhoods for genes with strong expression correlations to the neighborhood-level phenotype. Neighborhoods are ordered along the x-axis by phenotype value. The phenotype-correlated genes include markers of the CD56$^{br}$ (*ITGAX*, *CD44*, *SELL*, *GZMK*) and CD56$^{dim}$ (*FCGR3A*, *FGFBP2*, *CST7*) subtypes. (**D**) UMAP of all NK cells, colored according to cell assignment to the CD56$^{br}$ cluster. We also show the Pearson's r between CD56$^{br}$ cluster assignment status and csaQTL phenotype values across neighborhoods. (**E**) A boxplot of CD56$^{br}$ fractional abundance out of all NK cells per individual, organized by genotype at the lead SNP. We also show the variance explained by the lead SNP in the fraction of NK cells assigned to the CD56$^{br}$ cluster. (**F**) Locus zoom plot with one marker per tested SNP, genomic position along the x-axis, and GeNA p-value on the y-axis. Each SNP marker is colored by LD value relative to the lead SNP. The csaQTL lead SNP is labeled with a green diamond. The ovarian cancer risk and CXCL16 abundance pQTL lead SNPs are labeled with purple triangles, as is the lead SNP from the replicating association to CD56$^{br}$%NK previously found using flow cytometry.

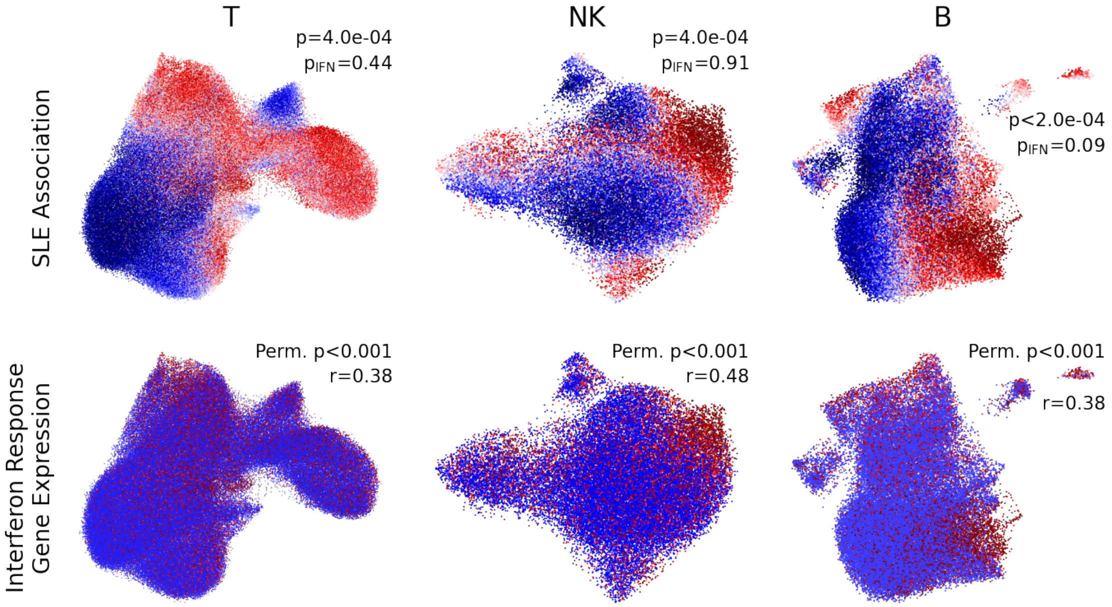


**Supplementary Figure 22: Re-analysis of the Perez *et al.* [7] SLE dataset within a neighborhood-based framework using CNA.** Re-analysis of the Perez *et al.* (2022) SLE case-control PBMC scRNA-seq dataset in our neighborhood-based framework enables a direct comparison of SLE PRS-associated cell states against cell states that differentiate patients with SLE from controls. The significant lupus case-control differences present in Perez *et al.* across major cell types (CNA Global p $<5\text{x}10^{-4}$ for T, B, NK, myeloid) correspond closely to interferon response gene expression across neighborhoods (Pearson's r 0.38-0.68 between SLE-associated phenotype values and interferon response gene expression per neighborhood, all with confirmatory bootstrapped p<0.001 for r>0). In fact, SLE case-control differences are eliminated in this model after controlling for average interferon response gene expression per sample (all CNA Global $p_{IFN}$ >0.08; **Methods**), suggesting that increased interferon response may be the primary perturbation present in peripheral blood cell state abundances during lupus disease. (**Top**) Consistent with the published results in Perez *et al.*, these data show strong cell state abundance shifts associated with SLE disease status across T, NK, and B cells with strong correspondence to interferon signaling. CNA global p-values shown with ($p_{IFN}$) and without (p) controlling for mean interferon response gene expression per individual. (**Bottom**) Interferon response gene expression per neighborhood among T, NK, and B cells. Pearson's r between interferon response per neighborhood and SLE phenotype is shown for each cell type, with associated bootstrapped p-value for r>0.

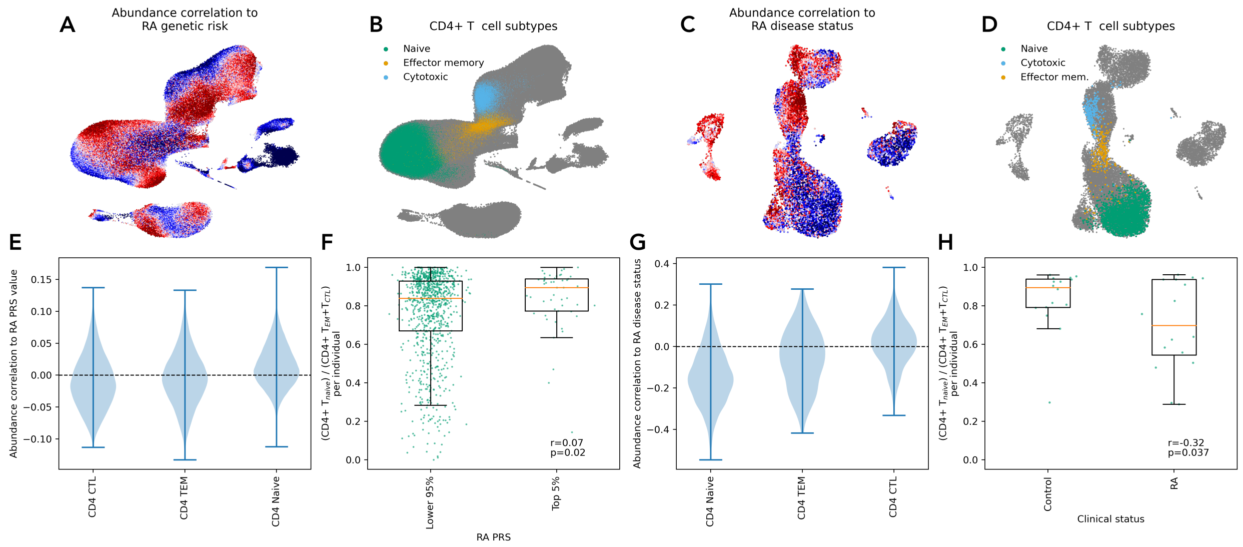


**Supplementary Figure 23: T cell state abundance shifts associated with increasing RA genetic risk and associated with RA disease status relative to controls.** Naïve-to-effector T cell ratios were computed per individual in the OneK1K cohort using cell assignments to the published T cell clusters. (**A**) UMAP of all cells from the OneK1K subcohort used to test for a cell state abundance association to RA PRS value (**Methods**). The UMAP is colored according the cell state abundance shift neighborhood-level phenotype associated with RA PRS value. (**B**) UMAP of all cells from the OneK1K subcohort used to test for a cell state abundance association to RA PRS value. The UMAP is colored according to cell assignments to naïve, effector memory and cytotoxic CD4+ T cell clusters. (**C**) UMAP of all cells from the OneK1K subcohort used to test for an RA case-control association (**Methods**). The UMAP is colored according the cell state abundance shift neighborhood-level phenotype associated with RA. (**D**) UMAP of all cells from the OneK1K subcohort used to test for an RA case-control association. The UMAP is colored according to cell assignments to naïve, effector memory and cytotoxic CD4+ T cell clusters. (**E**) The distribution of cell abundance correlations per neighborhood to RA PRS value within each highlighted cluster (naïve, effector memory and cytotoxic CD4+ T cells). (**F**) $T_{naive}/[T_{EM}+T_{CTL}]$ ratio among individuals with a known absence of RA, shown separately for individuals with the top 5% highest RA PRS values and for individuals with lower PRS values. We report the Pearson's r correlation. We also show a permutation-based p-value for $r>0$. (**G**) The distribution of cell abundance correlations to RA disease status within each highlighted cluster (naïve, effector memory and cytotoxic CD4+ T cells). (**H**) $T_{naive}/[T_{EM}+T_{CTL}]$ ratio among CD4+ T cells for individuals with RA as compared to healthy controls. We report the Pearson's r correlation. We also show a t-test p-value corresponding to the relationship between RA case-control status and this naïve-to-effector ratio. In (F) and (H), each boxplot displays the median (orange), Q1 and Q3 quartiles along with Q1–1.5xIQR (lower whisker) and Q3+1.5xIQR (upper whisker). Values per individual are shown in green with jittering along the x-axis.

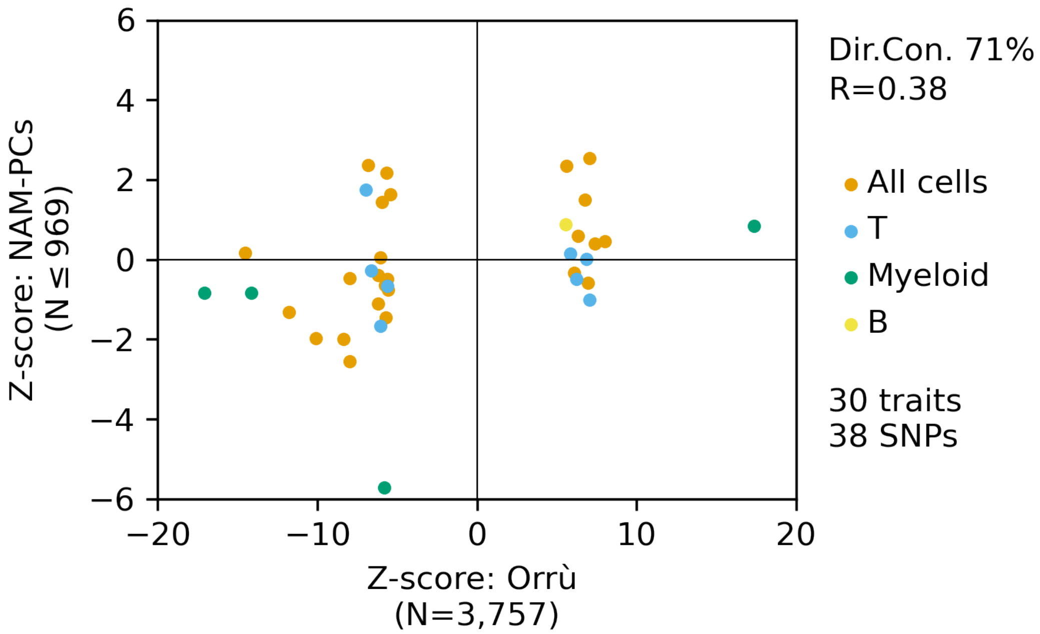


**Supplementary Figure 24: Comparison of effects captured by NAM-PCs to published associations, with respect to specific phenotypes**. Z-scores for published SNP associations to specific cell state abundance phenotypes quantified using flow cytometry by Orrù *et al.* are shown on the x-axis. For each SNP-trait pair, a corresponding Z-score is shown on the y-axis reflecting an association test in the OneK1K dataset between genotype and the best approximation of that phenotype that can be captured by NAM-PCs (**Methods**).

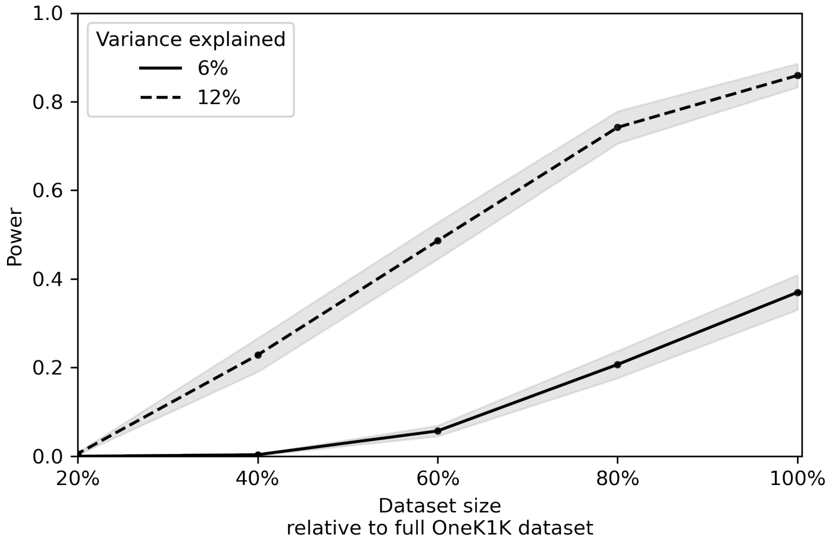


**Supplementary Figure 25: GeNA's statistical power increases linearly with the number of samples included in the single-cell dataset**. We downsampled the OneK1K dataset at random to 80%, 60%, 40% or 20% of the total donor count and repeated our power analysis simulation for each downsampled dataset. Here we plot statistical power by dataset size for simulated genotypes that explain 6% or 12% of variance in the associated cell state abundance shift trait.

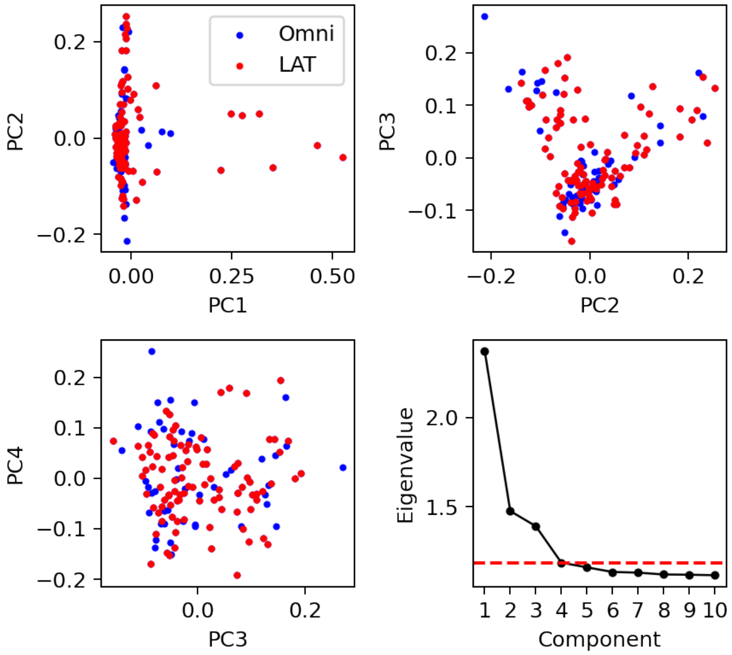


**Supplementary Figure 26: Genotype principal components for the Perez *et al.* [7] European-ancestry samples.** European-ancestry samples genotyped on the Omni and LAT arrays were merged after imputation. gPCs were constructed for this cohort using post-QC pre-imputation shared SNPs. The distributions of these samples, colored by genotyping array cohort, on gPCs 1-4 confirm that the gPCs capture within-ancestry genotypic variation, rather than reflecting genotyping array batch. We also display an elbow plot of eigenvalues by principal component. The red dashed line indicates the threshold used to select the number of gPCs included in our models.

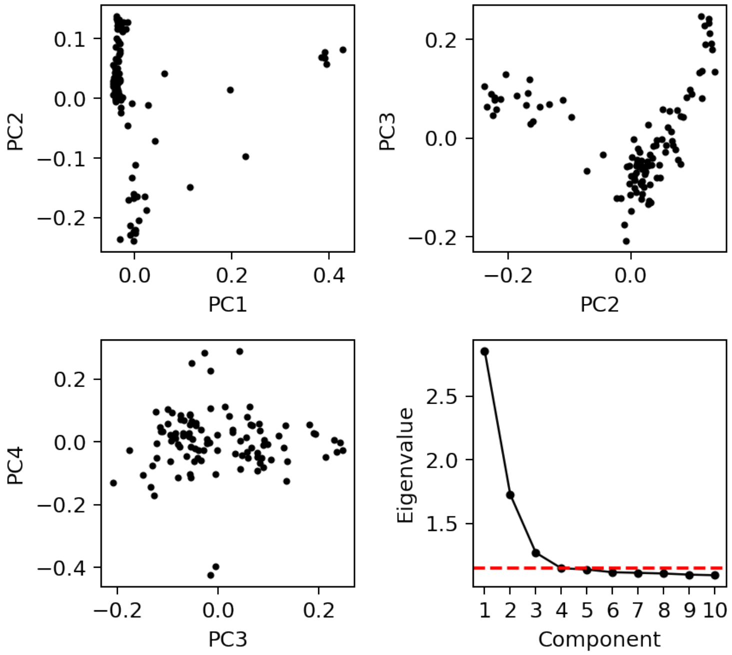


**Supplementary Figure 27: Genotype principal components for the Perez *et al.* [7] Asian-ancestry samples.** The distributions of these samples on gPCs 1-4 are shown along with an elbow plot of eigenvalues by principal component. The red dashed line indicates the threshold used to select the number of gPCs included in our models.

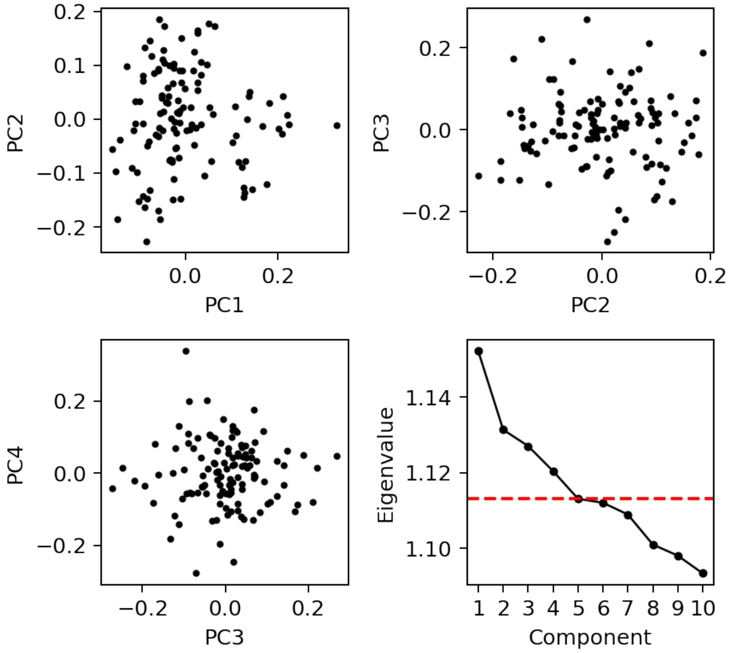


**Supplementary Figure 28: Genotype principal components for the Oelen *et al.* samples.** [8] The distributions of these samples on gPCs 1-4 are shown along with an elbow plot of eigenvalues by principal component. The red dashed line indicates the threshold used to select the number of gPCs included in our models.

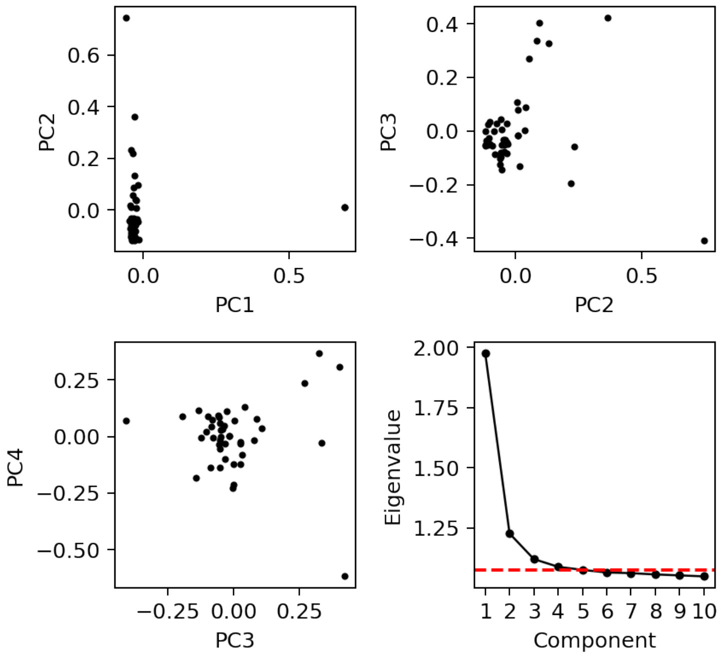


**Supplementary Figure 29: Genotype principal components for the Randolph *et al.* [9] European-ancestry samples.** The distributions of these samples on gPCs 1-4 are shown along with an elbow plot of eigenvalues by principal component. The red dashed line indicates the threshold used to select the number of gPCs included in our models.

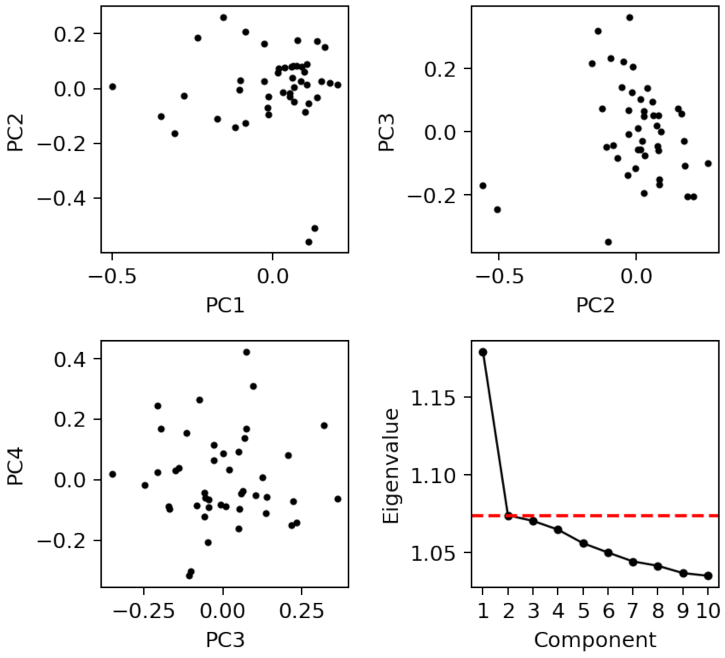


**Supplementary Figure 30: Genotype principal components for the Randolph *et al.* [9] African-ancestry samples.** The distributions of these samples on gPCs 1-4 are shown along with an elbow plot of eigenvalues by principal component. The red dashed line indicates the threshold used to select the number of gPCs included in our models.

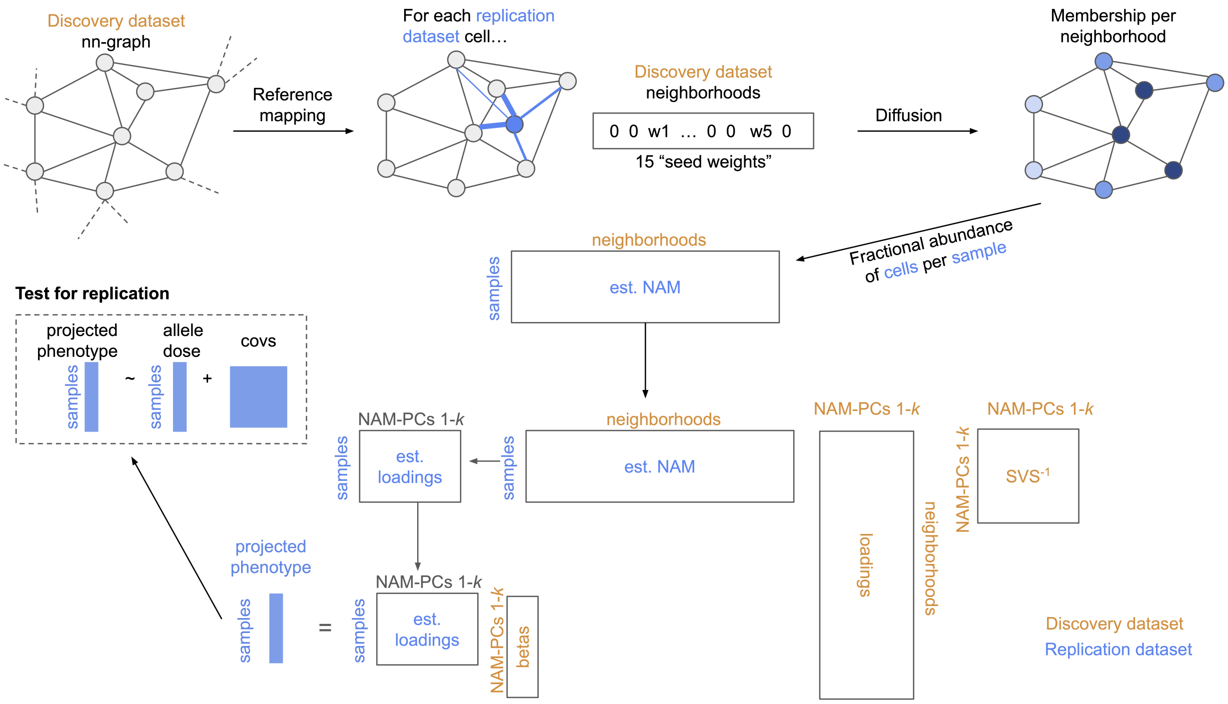


**Supplementary Figure 31: Schematic representation of our our approach to project a neighborhood-based phenotype into an independent dataset for testing of association replication**. We use a published reference mapping algorithm [10] to project each cell from the replication dataset (blue labels) into the embedding used for construction of the nearest neighbor graph from the discovery dataset (orange labels). For each replication dataset cell, we store its distance to the 15 nearest discovery dataset cells; these represent the seed weights of this replication dataset cell in the discovery dataset neighborhoods, of which there is one per discovery dataset cell. We use diffusion in the nearest neighbor graph, as we have previously described [3], to obtain from these seed weights the fractional membership of each replication dataset cell within all discovery dataset neighborhoods. For each replication dataset sample, the sum of neighborhood memberships across all cells in the sample yields the fractional abundance of that sample across discovery dataset neighborhoods. Row-wise stacking these per-sample vectors into a matrix produces an estimated Neighborhood Abundance Matrix (NAM) containing the distribution of each replication dataset sample across discovery dataset neighborhoods. We can then use the stored products of the discovery dataset NAM SVD to obtain loadings for each replication dataset sample on the discovery dataset NAM-PCs, as shown. Combining the replication dataset sample loadings on the discovery dataset NAM-PCs with the fitted coefficients that define the phenotype in the discovery dataset produces an estimated phenotype value per replication dataset sample, which we can use to test for association to the allele of interest (or case-control status) controlling for relevant covariates.

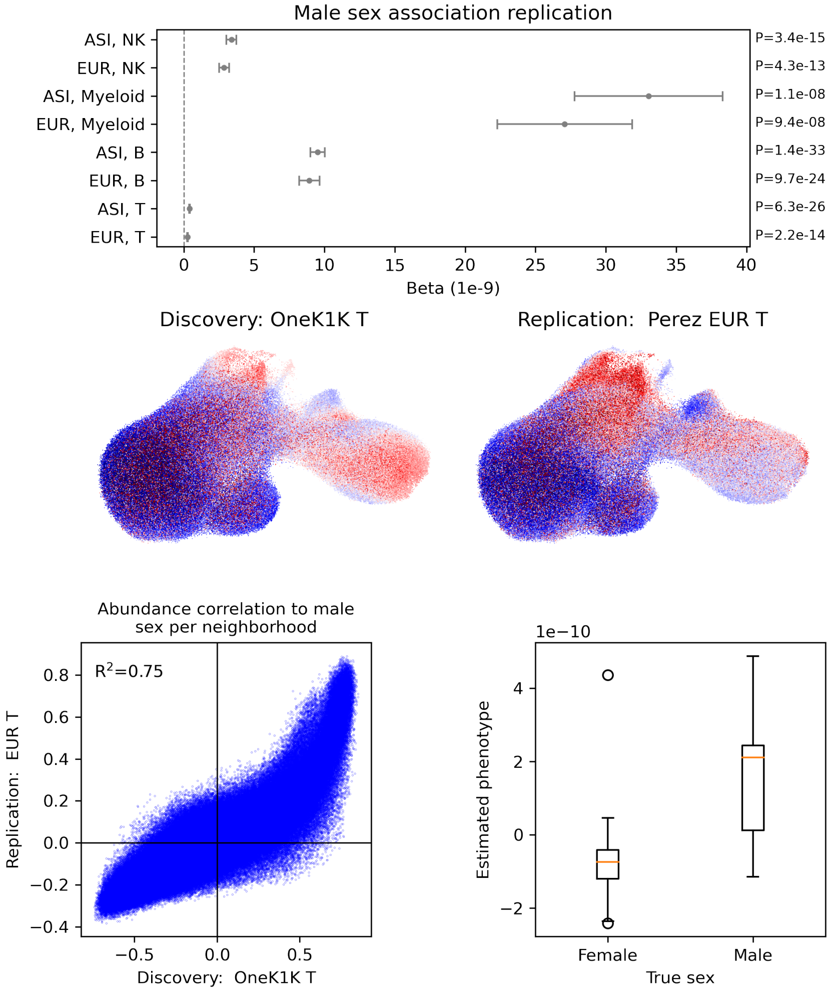


**Supplementary Figure 32: A sex example demonstrates the neighborhood-based phenotype projection and replication testing process**. (**Top**) Phenotypes associated with male sex within the OneK1K cohort were defined for NK, Myeloid, B and T cells. Each of these phenotypes was projected into the relevant cell subset of the Perez *et al.* dataset [7] and tested for association to ground-truth sex labels per sample. (**Middle**) Using the sex-associated phenotype within T cells as an example, we show per neighborhood the empiric correlations between cell abundance per sample and sex label for the discovery (left) and validation (right) datasets. Deeper red colors indicate larger positive correlations, while deeper blue colors indicate larger negative correlations. (**Bottom**) Within the replication dataset, we plot the projected per-neighborhood sex phenotype values against the empiric correlations per neighborhood between replication dataset cell abundance per sample and sex label (left; Pearson's $r^2$ shown). We also plot the distribution of estimated per-sample phenotype values for individuals in the replication dataset within each sex category (right).

### Supplementary Note

GeNA supports flexible detection of genotype-associated cell states in high-dimensional single-cell data. For some found csaQTL-associated phenotypes (e.g., if the SNP associates with abundance of naïve B cells), clustering and differential expression alone may suffice to define an equivalent phenotype in an independent replication dataset. Even in this simple case, however, 'naïve B' cluster boundaries may not be equivalent in the discovery and replication datasets. This challenge increases with more complex phenotypes (e.g., the SNP associates with a phenotype defined by one cell state abundance shift within CD4+ naïve T cells, another cell state abundance shift that spans CD8+ T memory types, and a depletion of MAIT cells). Defining an equivalent phenotype value per individual in a replication cohort that reflects this same pattern of change in global T cell composition could be prohibitively difficult without transferring neighborhood-scale phenotype information from the discovery dataset to the replication dataset. To address this challenge, we developed an approach to project a neighborhood-based cell state abundance phenotype from a discovery dataset to a replication dataset (**Methods**).

As an overview of our approach: consider a single-cell profiling discovery dataset in which we have identified an association between a sample attribute (e.g. sex, or allele dose for a genetic variant) and a tissue cellular composition phenotype using either CNA or GeNA. In order to evaluate replication of this association, we need to define a value per sample in a replication dataset that reflects an equivalent phenotype. The phenotype was originally defined in the discovery dataset through a linear association test between the attribute values (e.g., allele doses) per sample and sample loadings on the discovery dataset NAM-PCs. Using the fitted coefficient values per NAM-PC from the linear model, we can estimate replication sample phenotype values as long as we can estimate replication sample loadings on the discovery dataset NAM-PCs. We can estimate replication sample loadings on the discovery dataset NAM-PCs using the outputs from PCA on the discovery dataset NAM as long as we can generate a NAM for the replication dataset that stores the fractional cell abundance for each replication sample across discovery dataset neighborhoods. A schematic overview of this process is shown in **Supplementary Figure 31**.

Rather than define new neighborhoods for the replication dataset, we map replication dataset cells into the discovery dataset embedding and quantify the fractional abundance of each replication sample across discovery dataset neighborhoods. Further, rather than define new axes of inter-sample variation specific to the replication dataset, we determine the replication dataset sample loadings on the discovery dataset NAM-PCs. Using the fitted coefficient values per NAM-PC that define the phenotype in the discovery dataset, we obtain phenotype values per replication dataset sample and can test the association of the attribute of interest (e.g., allele dose or case-control status) to the phenotype within the replication dataset.

We demonstrate projection and replication testing with a non-genotype example: sex (**Supplementary Figure 32**). In a neighborhood-based framework, we have previously shown that sex associates with simultaneous changes in blood cell type relative abundances as well as the usage of a sex chromosome gene expression program across cell types [3]. The sex-associated cell state abundance shift therefore represents a multi-factorial phenotype that includes both coarse (cell type abundance changes) and granular (differential expression of sex chromosome genes) components. A sex-associated phenotype defined within the OneK1K dataset and projected into the Perez et al.[7] PBMC sc-mRNA-seq dataset yields replicating associations ($p<2x10^{-7}$) to ground-truth sex labels across all major cell types (T, B, NK, myeloid) in both ancestry cohorts (Asian and European, N=98 and N=140 samples respectively).
